# Light-microscopy based dense connectomic reconstruction of mammalian brain tissue

**DOI:** 10.1101/2024.03.01.582884

**Authors:** Mojtaba R. Tavakoli, Julia Lyudchik, Michał Januszewski, Vitali Vistunou, Nathalie Agudelo, Jakob Vorlaufer, Christoph Sommer, Caroline Kreuzinger, Barbara Oliveira, Alban Cenameri, Gaia Novarino, Viren Jain, Johann Danzl

## Abstract

The information-processing capability of the brain’s cellular network depends on the physical wiring pattern between neurons and their molecular and functional characteristics. Mapping neurons and resolving their individual synaptic connections can be achieved by volumetric imaging at nanoscale resolution with dense cellular labeling. Light microscopy is uniquely positioned to visualize specific molecules but dense, synapse-level circuit reconstruction by light microscopy has been out of reach due to limitations in resolution, contrast, and volumetric imaging capability. Here we developed light-microscopy based connectomics (LICONN). We integrated specifically engineered hydrogel embedding and expansion with comprehensive deep-learning based segmentation and analysis of connectivity, thus directly incorporating molecular information in synapse-level brain tissue reconstructions. LICONN will allow synapse-level brain tissue phenotyping in biological experiments in a readily adoptable manner.

**One-Sentence Summary:** Hydrogel expansion enables molecularly informed reconstruction of brain tissue at synaptic resolution with light microscopy.

---

The brain constitutes an incredibly dense, complex, and fine-grained arrangement of neurons with their support cells into a functional network enabling all brain function. Imaging approaches are uniquely positioned to decode the brain’s spatial organization. To determine how individual neurons are connected and to reconstruct the circuitry underlying information processing, i.e. to determine connectomes, demands the capacity to accurately trace cellular circuit components, including axons and dendritic spines, resolve the individual synaptic connections, and assign them to specific neurons.

Light microscopy holds tremendous potential for unifying synapse-level circuit reconstruction with in-depth molecular characterization. However, it is conventionally limited in resolving power by light-wave diffraction to a few hundred nanometers (best case 200-300 nm laterally and typically substantially worse (∼1000 nm) along the optical (*z*-)axis), far too coarse grained to distinguish densely labelled cellular structures in brain tissue. Electron microscopy (EM) with its nanometer-scale resolution and comprehensive structural contrast is currently the only technology allowing dense connectomic analysis, i.e. comprehensive reconstruction of the cellular circuit components (*1–3*), with which enormous strides have been made in mapping connectivity in organisms as diverse as worm (*4*), fly (*5–7*), mouse (*8*) or humans (*9*, *10*). These advances were facilitated by technological progress in automated data collection as well as deep-learning analysis, making the challenge of densely annotating all cellular structures tractable (*2*, *3*). EM sample preparation and readout are not usually compatible with direct visualization of specific molecules in circuit reconstruction experiments, requiring correlation with light microscopy to obtain molecular information (*11–13*). This contrasts with the crucial functional reliance of neurons and synapses on their specific molecular machineries, reflected in enormous molecular diversity both at cellular and synaptic levels. EM reconstructions do allow inferring connectivity via chemical synapses from structural features with high accuracy (*14–18*). However, they are incomplete in the sense that in pure EM data, synapses cannot be further differentiated molecularly and important information related to signaling between cells, like e.g. the distribution of receptor molecules playing important roles also beyond classical synaptic transmission, remains hidden.

Super-resolution optical imaging offers resolution beyond the optical diffraction limit either by increasing instrument resolution (*19–22*) or expanding samples to increase distances between features (*23*) but has mostly been limited to studying sparse subsets of brain cells or the distribution of molecules devoid of cellular context. To visualize brain tissue comprehensively, fluorophores have been applied extracellularly to cast super-resolved “shadows” of all cells in living tissue when read out with stimulated emission depletion (STED) microscopy (*19*) in super-resolution shadow imaging (SUSHI) (*24*). Combining this with two-stage machine learning allowed reconstructing the cellular architecture at 3D-nanoscale resolution with an approach termed LIONESS (*25*). Fixation-compatible extracellular labeling with the CATS approach (*26*) visualized brain tissue architecture across spatial scales, both using STED or expansion microscopy (ExM). While LIONESS and CATS allowed reconstructing tissue architecture at nanoscale detail, they have not provided the traceability and accuracy required for synapse-level circuit reconstruction. ExM (*23*) uses embedding in a swellable hydrogel followed by disruption of the tissue’s mechanical cohesiveness and expansion, thus increasing effective resolution. Expansion factor (exF) and corresponding resolution enhancement have been increased from ∼4-fold (*23*, *27*) to ∼8-10-fold (*28–32*) in single-step approaches and to ∼16x and beyond with iterative application (*33–35*) of two swellable hydrogels. Indiscriminate (“pan-“) labeling for protein density using amine-reactive fluorophore derivatives, such as N-hydroxysuccinimidyl(NHS)-esters, has revealed cellular ultrastructure with single-step (*30*, *36*) and iterative (*35*) expansion and visualized the complexity of brain tissue (*30*, *37*). However, despite these advances, it has thus far not been possible to achieve light microscopy imaging of brain tissue at the resolution and signal-to-noise ratio required for tracing the finest neuronal processes and dense connectomic reconstruction.

Here we report on a technology to densely reconstruct brain circuitry with light microscopy at synaptic resolution. For this, we engineered a high-fidelity iterative hydrogel expansion scheme paired with protein-density staining and high-speed diffraction-limited readout that enables manual neuronal tracing and deep-learning based automated cellular segmentation (**Fig. 1A**). We show traceability of the finest neuronal structures, including axons and dendritic spines, simultaneous molecular measurement, deep-learning based prediction of molecule locations, and connectivity analysis at single synapse resolution. While direct comparison in the same sample with volume EM is not possible for expanded specimens, we provide a series of metrics for validation. This includes validation against independent ground truth based on a sparse positive cellular label and quantification of manual traceability of spines. We furthermore provide comparison of statistical data on neuronal connectivity with previous EM measurements, which is an established method that has been used to compare EM datasets among each other (*38*). This technology, which we term LICONN for *Light Microscopy based Connectomics*, offers molecularly informed reconstruction of brain tissue in a broadly accessible manner.

**Fig. 1.**
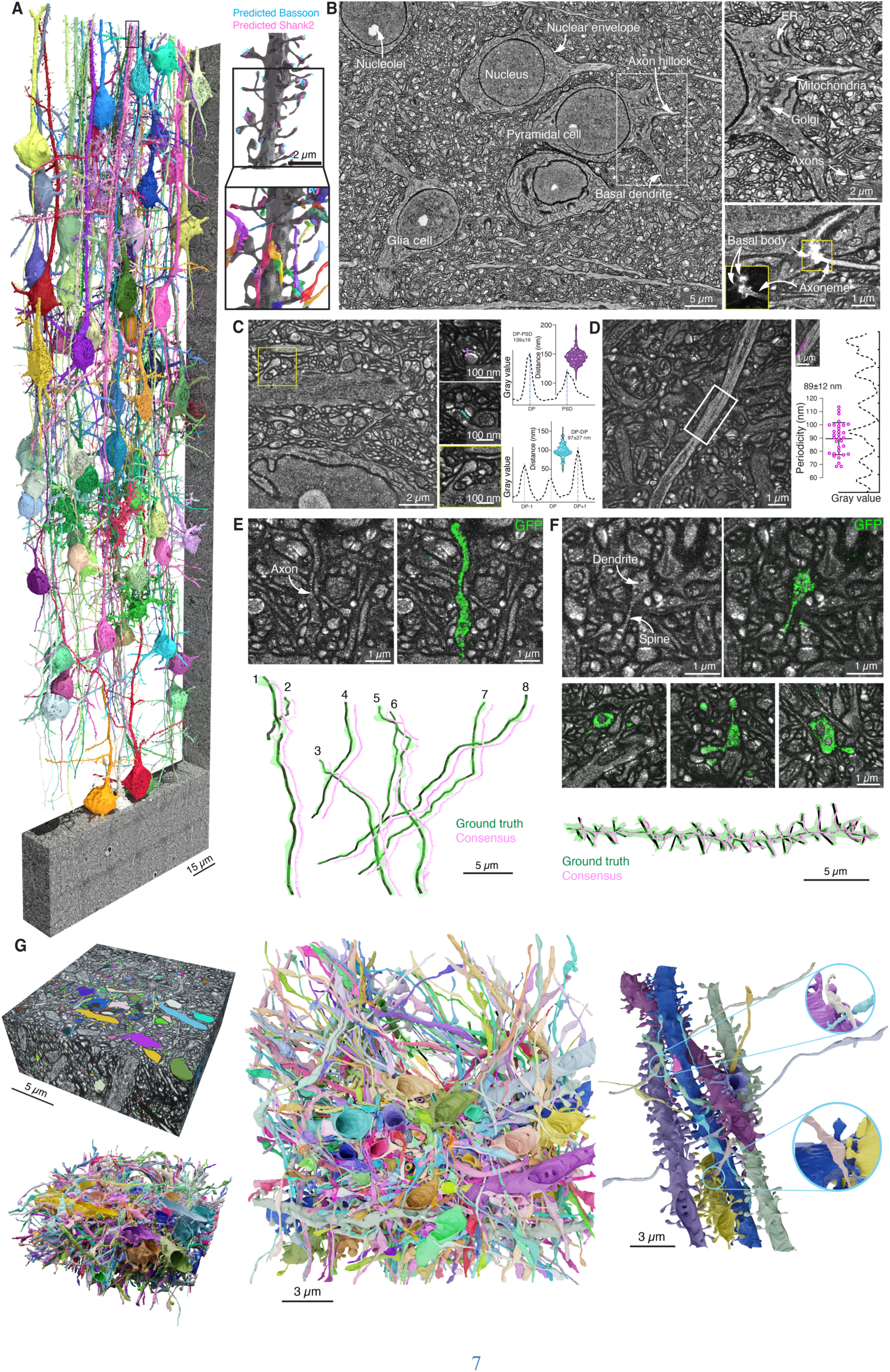
Dense connectomic reconstruction of mammalian brain tissue with light microscopy. (**A**) ∼1 Million cubic micrometer (native tissue scale) LICONN volume of mouse primary somatosensory cortex spanning layers II/III-IV (396 x 109 x 22 µm^3^ original tissue scale, 0.95 Million µm^3^ before and 3.5*10^9^ µm^3^ after ∼16x hydrogel expansion). Length scales and scale bars refer to biological size before hydrogel expansion throughout. 3D-renderings of 79 example cells from a dense reconstruction produced by a flood-filling network (FFN; see https://neuroglancer-demo.appspot.com/#!gs://liconn-public/ng_states/fig1a.json for original data and segments). Magnified views: *Top*: 3D-rendering of a dendrite (black) from a pyramidal neuron, corresponding to the box at the top of the main panel. Additionally, deep-learning based predictions of the location of the synaptic molecules Bassoon (cyan, pre-synapses) and Shank2 located at dendritic spines (magenta, excitatory post-synapses) are displayed. *Bottom*: Same dendrite with synaptically connected axons as inferred from the prediction of molecules associated with synaptic sites. Imaging time: 6.5 h (effective voxel rate 17*10^6^ voxel/s (17 MHz)). Displayed cells were completed by manual agglomeration of the FFN segmentation. Imaging data representative of expansion and multi-tile imaging in *n*=10 replicates across >3 animals. (**B**) Single imaging plane in a subregion of panel A with cell bodies embedded in dense neuropil. Magnified views: *Top*: Intracellular structure in a pyramidal neuron with surrounding neuropil. *Bottom*: Primary cilium in a different region of the same dataset. Raw spinning-disk confocal imaging data with Contrast Limited Adaptive Histogram Equalization (CLAHE) applied (comparison of raw and CLAHE-processed data: **fig. S12**). (**C**) Single-plane view of dendrite with spine (yellow box, raw data with CLAHE). In the bottom, a cell nucleus is visible with nuclear pores revealed as bright densities. Small panels: *Top*: excitatory synapse with dense, protein-rich punctate features at pre-synapse and bar-like feature at post-synapse (raw data without CLAHE). The line indicates direction of line intensity profiles (*top right*) for measuring the distance between pre- and post-synaptic dense features with corresponding violin plot (DP: dense projection, PSD: post-synaptic density) (139±19 nm, mean±s.d., 192 synapses, 3 technical replicates across *n*=2 animals). *Middle*: Single bouton contacting two spines, evident from the respective high-density features. The line indicates direction of line intensity profiles (*bottom right*) for measuring the distance between dense features at pre-synapses with corresponding violin plot (97±27 nm, mean±s.d., 192 synapses). *Bottom*: Highlighted spine. (**D**) Periodic, protein-dense structure at the circumference of a subset of neurites (without CLAHE). Example line profile (direction indicated in inset) and plot of periodicity (89±12 nm, mean±s.d., 32 distance measurements across *n*=2 animals). (**E**) Manual tracing of axons. *Top, left*: Zoomed view of a single plane of LICONN volume (∼19 x 19 x 19 µm^3^) from primary somatosensory cortex in *Thy1-eGFP* mouse with cytosolic eGFP expression in an axon (arrow). *Right*: Overlay of structural channel (gray) with eGFP visualized via immunolabeling. *Bottom*: Renderings (green) and skeletons (black) of 8 eGFP-expressing axon stretches (Ground truth, based on eGFP and structural channels) in a LICONN dataset, compared to skeletons generated by two annotators blinded to eGFP channel (magenta, Consensus). Skeletons are offset for visual clarity. Additional datasets: **figs. S13** and **S14**. Representative of *n*=12 datasets (3 technical replicates in n=3 animals). (**F**) Manual tracing of dendrites. *Top, left*: Single plane (zoomed view) of structural channel in LICONN volume (∼19 x 19 x 19 µm^3^) from hippocampal CA1 in *Thy1-eGFP* mouse with eGFP-expressing dendrite (arrow). *Right*: Overlay of structural channel (gray) with eGPF signal visualized by immunolabeling (green). *Middle*: Cross sections of dendrites and spines with eGFP expression embedded in dense neuropil. *Bottom*: Skeleton (black) generated by taking eGFP- and structural channels into account (ground truth), enclosed in 3D-rendering of dendrite (green). Spines are indicated as branches. Additional overlay with a skeleton (magenta) generated from structural LICONN channel by two annotators blinded to eGFP. Additional datasets: **fig. S15**. Representative of *n*=5 datasets (3 technical replicates in *n*=2 animals). (**G**) *Left*: Single-tile LICONN volume (gray) from hippocampal CA1 with sparse, manual cellular annotations (color, 658 structures) and 3D-rendering of the annotated structures. *Middle*: Top view. *Right*: Subset of dendrites and axons with enlarged views of synaptic connections. Enlarged views were generated with a different camera position.

## Results

### Brain tissue expansion for connectomics

Our strategy for dense light-microscopy based connectomics achieves resolution increase exclusively via hydrogel expansion rather than optical super-resolution to take advantage of the speed, optical sectioning capability and wide availability of standard, diffraction-limited, spinning disk confocal microscopes for imaging. Hydrogel embedding (*39*) and expansion homogenize sample refractive index and adjust it to that of water. This facilitates acquisition of tissue volumes extended both laterally and along the optical (*z*)-axis, which is notoriously difficult with other optical super-resolution techniques due to aberrations, scattering and photobleaching. This strategy also allows facile incorporation of multi-color readout of specific molecules. In contrast to typical correlative light/electron microscopy workflows for circuit reconstruction, it delivers molecular information at essentially identical resolution as the structural channel.

To achieve high-fidelity tissue preservation, traceability of neuronal structures and experimental robustness, we developed an iterative expansion technology with truly independent, interpenetrating hydrogel networks and specifically tailored chemical fixation, cellular protein retention, and hydrogel chemistry that obviated hydrogel cleavage and signal handover steps. Available single-step or iterative approaches did not provide the required performance and straightforward modifications to increase expansion factor resulted in unstable hydrogels (**figs. S1, S2, S3, table S1**).

**Fig. 2.**
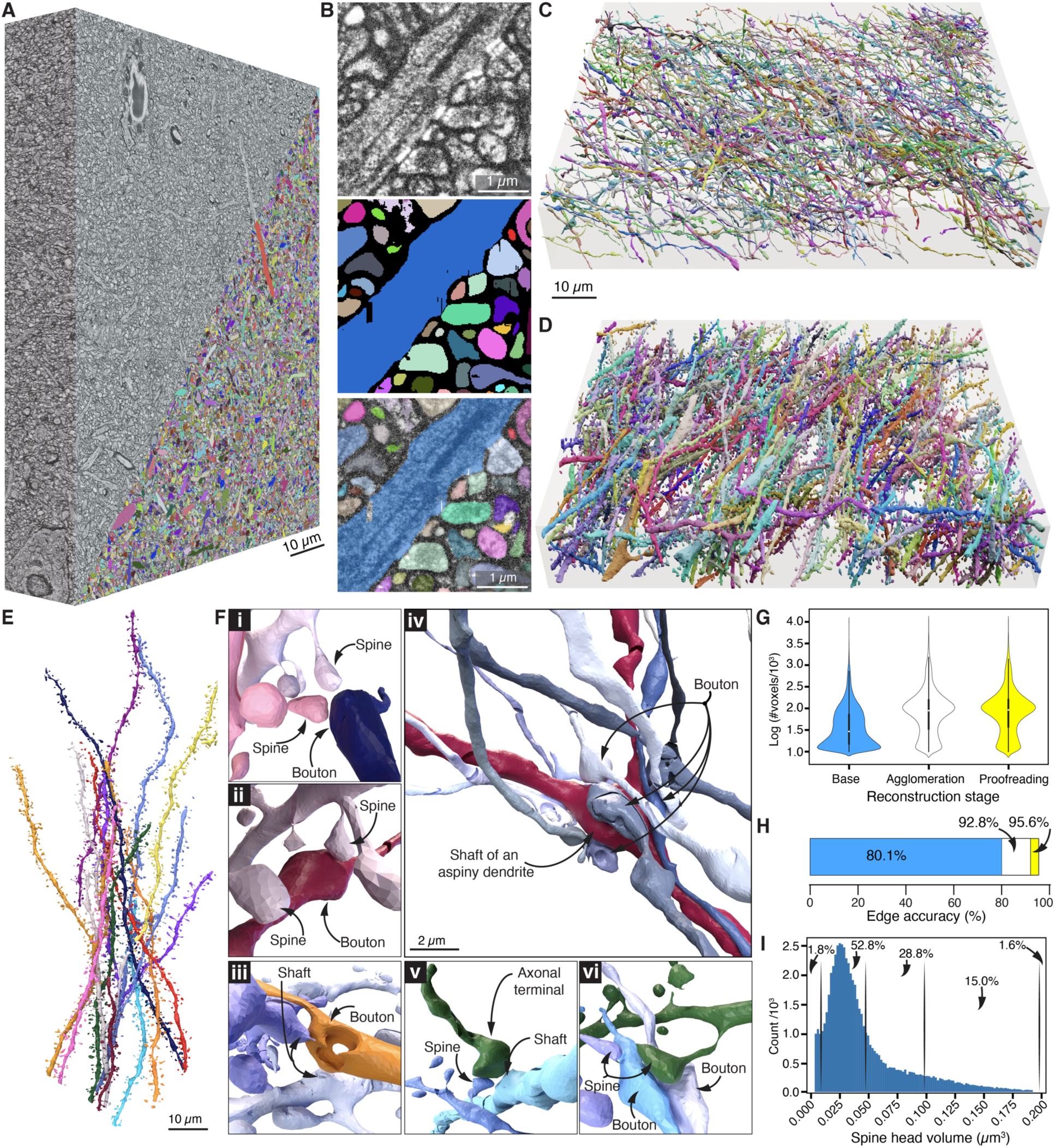
Deep-learning based segmentation. (**A**) Rendering of 85 x 69 x 14 µm^3^ LICONN volume (native tissue scale) from hippocampal neuropil in CA1, overlaid with dense FFN-based segmentation of neuronal structures in the bottom corner. Neuronal structures were comprehensively proofread in this volume by correction of split and merge errors (without any manual painting of voxels; for original data and proofread segmentation see https://neuroglancer-demo.appspot.com/#!gs://liconn-public/ng_states/expid82.json). Data representative of FFN based segmentation in 6 datasets. Comprehensive proofreading of all neuronal structures was performed in one dataset. (**B**) Magnified view from a single plane with raw structural data (CLAHE applied), dense segmentation after proofreading, and overlay. (**C**) Rendering of 5.8% of axons (see https://neuroglancer-demo.appspot.com/#!gs://liconn-public/ng_states/expid82_fig2_axons.json for browsable data) in the volume in (A). (**D**) Rendering of 27.3% of dendrites and a small number of axons (see https://neuroglancer-demo.appspot.com/#!gs://liconn-public/ng_states/expid82_fig2_dends.json for browsable data). (**E**) Rendering of example dendrites. (**F**) Spatial arrangement of selected axonal and dendritic structures, highlighting various types of contacts. (**G**) Segment size (number of voxels on a logarithmic scale) of neuronal structures for the base FFN segmentation (blue), after automated agglomeration (white), and after full manual proofreading of the automated agglomeration (yellow). (**H**) Edge accuracy for the base segmentation (blue), after automated agglomeration (white), and after manual proofreading (yellow). (**I**) Distribution of spine head volumes in the same volume, analyzed for 59,332 spines. Percentage numbers refer to the intervals indicated by the vertical lines (<0.01, 0.01, 0.05, 0.1, 0.2, >0.2 µm^3^).

**Fig. 3.**
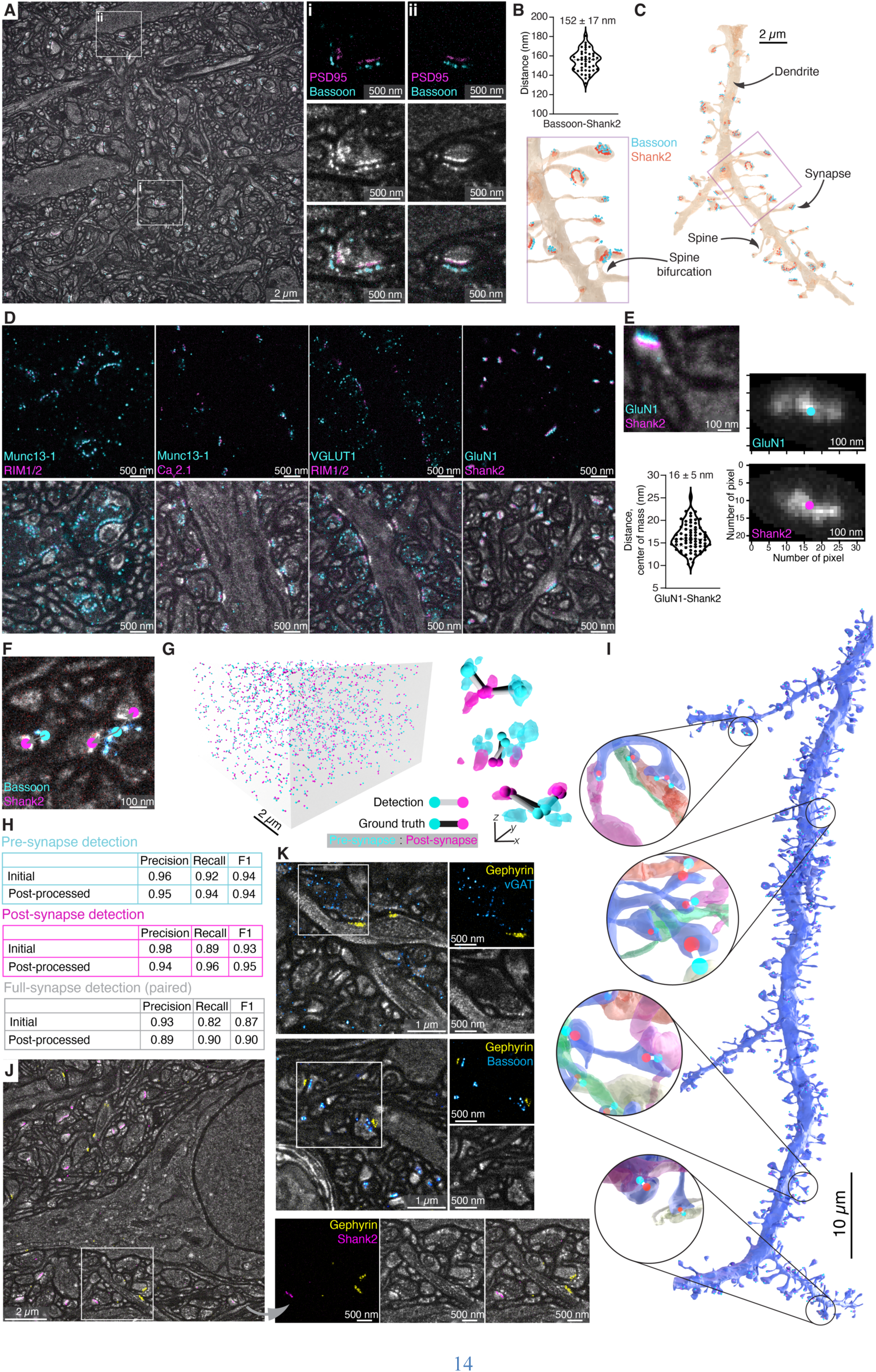
Molecular labeling and detection of synaptic connections. (**A**) Immunolabeling for pre-synaptic Bassoon (cyan) and post-synaptic PSD95 (magenta, excitatory post-synapses) paired with the structural LICONN channel (gray), displaying single plane from a LICONN volume of somatosensory cortex. (i, ii) Magnified views of individual synapses showing the immunolabeling and structural channels separately and as overlay. Bassoon/PSD95 labeling was performed in *n*=3 replicates across 2 animals. **(B)** Violin plot of distances between Bassoon and Shank2 signals (152±17 nm, mean ±s.d., 106 synapses measured across *n*=2 animals). **(C)** 3D-renderings of Bassoon and Shank2 immunolabeling signals mapped onto the corresponding dendrite from an FFN segmentation of the volume in panel A. Inset: magnified view of the boxed region. Bassoon/Shank2 labeling was performed in *n*=5 replicates across 3 animals. **(D)** LICONN with immunolabeling for pre- and post-synaptic markers (*top*) and overlay with structural channel (*bottom*). Single planes from volumetric datasets in the hippocampal CA3 stratum lucidum (leftmost panel) and CA1. Individual immunolabelings were replicated in various combinations between *n*=2 and *n*=6 times according to experimental needs. **(E)** Example of an individual dendritic spine with labeling for Shank2 (magenta) and the NMDA receptor subunit GluN1 (cyan). Plot of the distance between the center of mass of GluN1 and Shank2 distributions (16±5 nm, mean ±s.d., 184 synapses evaluated in *n*=1 animal) with maximum intensity projections of the immunolabeling signal. The respective centers of mass are indicated by points. **(F)** Illustration of computational detection of excitatory synapses based on immunolabeling for Bassoon (cyan) and Shank2 (magenta), overlaid with the structural channel (gray). Immunolabeling signals are converted to the respective point annotations for pre- (cyan) and excitatory post-synapses (magenta). Examples include synapses with 1:1 and 1:2 connections between pre- and post-synapses. **(G)** *Left*: Rendering of ground truth (proofread) immunolabeling based excitatory synapse detections in a 913 µm^3^ volume from the hippocampal CA1, stratum radiatum. *Right:* Magnified views of specific examples of detected synapses, overlaid with 3D renderings of the immunolabeling channels. Synaptic connections are indicated by cyan and magenta balls for pre- and post-synapses, respectively. Gray bars correspond to computationally detected synaptic connections. Black bars indicate manually generated ground truth overlaid for comparison. *Top*: Two boutons contacting a single post-synapse. *Middle*: 1:1 connection. *Bottom*: One pre-synapse connected to two post-synapses. **(H)** Precision, recall and F1 score (range 0-1, combining precision and recall) of computational immunolabeling-based detection of pre-synapses, post-synapses, and fully assembled synapses, evaluated using manually generated ground truth in the dataset in G. Metrics are given both for the base detection and with a post-processing step (materials and methods). See **fig. S22** for additional test dataset. **(I)** 3D-rendering and magnified views of a dendrite (segmented with FFN) from hippocampus and excitatory synaptic connections (bars) detected via immunolabeling for Bassoon (pre-synapses, cyan balls) and Shank2 (post-synapses, red balls). Magnified views additionally include the synaptically connected boutons. **(J)** Single plane from a LICONN volume with immunolabeling for the post-synaptic scaffolding proteins Gephyrin (yellow, inhibitory post-synapses) and Shank2 (cyan, excitatory post-synapses) with the immunolabeling and structural (gray) channels shown separately and as overlay for the boxed region. Inhibitory post-synapses show less pronounced features in the structural channel than their excitatory counterparts. Immunolabelings of inhibitory synapses were replicated at least *n*=5 times. **(K)** *Top*: LICONN with immunolabeling for Gephyrin (yellow) and vesicular GABA transporter (vGAT, cyan), indicating inhibitory pre- and post-synapses, respectively, with immunolabeling and structural (gray) channels displayed separately for the boxed region. *Bottom*: Similar measurement with immunolabeling for Bassoon (cyan, excitatory, and inhibitory pre-synapses) and Gephyrin (yellow, inhibitory post-synapses).

We first transcardially perfused mice with hydrogel-monomer (acrylamide, AA) containing fixative solution equipping cellular molecules with vinyl residues to later co-polymerize with the swellable hydrogel. Optimization of the perfusion solution, with lower monomer concentration (10% AA) than previously used (*27*, *37*) markedly improved the preservation of cellular architecture (**fig. S4**). This likely reflects osmotic effects known to impact tissue preservation, which has been utilized to enhance traceability in EM connectomics (*40*). We harvested and sliced brains, and then exploited the broad chemical reactivity of multi-functional epoxide compounds (*41*, *42*) to further stabilize and functionalize cellular molecules. We used glycidyl methacrylate (GMA) to attach acrylate groups to proteins more broadly than achieved with commonly employed amine-reactive compounds, further improving anchoring into the hydrogel. A molecule featuring three epoxide rings on a branched backbone, glycerol triglycidyl ether (TGE), was introduced to further fix and stabilize cellular molecules during downstream processing. Alternatively, amine-reactive anchoring (*43*, *44*) using N-acryloxysuccinimide (NAS) (*32*) resulted in datasets that were in principle traceable (**figs. S2, S5 and S6**) but the use of epoxides improved definition of cellular ultrastructure and emphasized synaptic features, both of which were desirable for our analysis.

We polymerized an expandable acrylamide/sodium-acrylate hydrogel, integrating functionalized cellular molecules into the hydrogel network. We then used heat/chemical denaturation (*27*) to disrupt mechanical cohesiveness while retaining cellular proteins (**fig. S7**) and, after expanding this first hydrogel ∼4-fold, applied optional immunolabeling to visualize specific proteins. A non-expandable stabilizing hydrogel prevented shrinkage during application of a second swellable hydrogel intercalating with the first two hydrogels. To achieve structural preservation and homogeneous expansion, we optimized the composition of each hydrogel (**fig. S8 and S9**). We further found that chemically inerting any unreacted groups in the hydrogel network after each polymerization step facilitated high-fidelity expansion by abolishing crosslinks between individual hydrogels and ensuring independence. Finally, we utilized protein-density (“pan-protein”) staining with fluorophore NHS-esters to comprehensively visualize cellular structures, mapping (primary) amines abundant on proteins at various amino acids and N-terminals.

These triple-hydrogel/sample hybrids featured ∼16-fold overall expansion and were mechanically robust, facilitating handling and extended imaging (exF=15.44±1.3, mean±s.d., 18 imaging volumes from *n*=6 technical replicates recorded across *n*=3 animals, **fig. S10**; unless otherwise stated, we give length measures in original tissue size by scaling measured post-expansion lengths with this exF). Distortions introduced by the expansion procedure were similar to those in previous work (*33*, *34*, *37*) (**fig. S10**). Spinning-disc confocal imaging with a high-numerical aperture (NA=1.15) water-immersion objective lens for refractive index matching in the green spectral range yielded an expected optical resolution of ∼280 nm laterally and ∼730 nm axially. With ∼16-fold resolution increase by expansion, this translated into ∼20 nm and ∼50 nm effective spatial resolution, respectively, demanding effective voxel sizes in the imaging of ∼10 x 10 x 25 nm3 for adequate spatial sampling. Overall, this procedure resulted in a robust and straightforward workflow and provided the resolution, contrast, and experimental throughput required for connectomic tissue reconstruction.

We analyzed a ∼1 Million cubic micrometer (native tissue scale, 396 x 109 x 22 µm^3^, 0.95*10^6^ µm^3^ before expansion, effective voxel size of 9.7 x 9.7 x 25.9 nm^3^, 3.5*10^9^ µm^3^ after expansion) tissue volume spanning layers II/III - IV in primary somatosensory cortex (**Fig. 1A**, **fig. S11,** see https://neuroglancer-demo.appspot.com/#!gs://liconn-public/ng_states/fig1a.json for original data and segments). 132 partially overlapping subvolumes were imaged (arranged on a 6x22 grid) and an automated algorithm (“SOFIMA” (*45*)) was used to achieve seamless volume fusion. Due to the high degree of parallelization in spinning disc confocal microscopy (∼300 focal points in our system), the ∼4.2*10^9^ cubic microns (post-expansion size, 0.47 Teravoxels, including tile overlap) were imaged within 6.5 h. This corresponded to 17*10^6^ voxels/second (17 MHz) effective voxel rate including overhead due to sample stage movement and tile overlap. Individual neurons with their axons and dendrites were clearly delineated from each other in the densely packed neuropil and showed rich subcellular structure (**Fig. 1B-D, fig. S12**), including e.g. mitochondria, the Golgi apparatus, and primary cilia. Inspecting dendrites and their spines, specialized post-synaptic structures typical of excitatory synapses (**Fig. 1C**), we found putative synaptic transmission sites highlighted by protein-rich, high-intensity features, akin to post-synaptic densities (PSDs) in EM data of chemically fixed specimens (*14*). Similarly, pre-synaptic sites displayed protein-dense nanoscale features, in this case arranged in a lattice-like pattern spaced 97 ± 27 nm, at a distance of 139 ± 19 nm (evaluated at 192 synapses in 3 technical replicates recorded across *n*=2 animals) from the post-synaptic densities (**Fig. 1C**), which was again similar to features observed in EM. We also observed a prominent ring-like periodic pattern at the circumference of a subset of neurites (**Fig. 1D, fig. S12**). The periodicity of 89 ± 12 nm (32 distance measurements from *n*=2 animals) was highly suggestive of the actin/β-spectrin cytoskeletal lattice organizing the distribution of specific proteins (*46*) and hence protein density below the plasma membrane of neurites. Periodicity was consistent with the previously reported value of 182 nm when only one component was selectively labelled (*46*), which further corroborated the faithfulness of the expansion procedure at the nanoscale. Together, this indicated that our expansion and imaging procedure revealed the cellular constituents of brain tissue with high fidelity from tissue- to nano-scale.

### Tracing neuronal structures in LICONN data

Thin, tortuous axons in dense neuropil and dendritic spines with their thin necks are among the most challenging structures for connectomic tracing. To evaluate the reliability of manual tracing, we compared human consensus skeletons to ground truth from sparse eGFP labeling. We obtained “ground truth” neurites (*47*) from cytosolic expression of enhanced green fluorescent protein (eGFP) in a sparse subset of neurons in *Thy1-eGFP* mice and compared those with independently traced skeletons of the same objects manually generated exclusively from the LICONN structural channel (**Fig. 1E,F**). We imaged the positive label in a separate color channel with comparable post-expansion resolution as the structural channel. Two novice human tracers were trained with both the eGFP and the structural LICONN channels of four datasets (∼19x19x19 µm^3^), containing in total two eGFP-expressing axons and two eGFP-expressing dendrite stretches. These datasets were excluded from further analysis. The tracers then received 12 similar LICONN datasets with 37 axon stretches (880 µm cumulative length of eGFP-expressing axons, recorded with *n*=3 technical replicates across *n*=2 animals in cortex), with a seed point in each eGFP-expressing axon while being blinded to the eGFP signal itself. They were asked to independently trace the indicated axons, compare results, and find consensus at locations of disagreement (**Fig. 1E**). Out of the 37 axon stretches analyzed, the structural channel consensus skeletons followed a wrong path in one case compared to eGFP ground truth (1.1 errors/mm, **Fig. 1E, figs. S13 and S14**). In a similar analysis of eGFP-expressing dendrites (on which spine locations were marked by branch points of the respective skeletons), the blinded tracers correctly identified 259 out of 289 spines (**Fig. 1F, fig. S15**, 90%, *n*=3 technical replicates recorded across *n*=2 animals in hippocampus).

Encouraged by the strong consistency between LICONN-derived skeletons and their eGFP ground truth, we returned to the 1 Million cubic micrometer cortical dataset in **Fig. 1A**. We validated traceability of axons and dendrites across tile borders (**fig. S16**). We further sought to exclude the possibility that there was a sizable number of non-traceable spines, employing exhaustive tracing analysis of local volumes. We sampled 39 subvolumes of 2x2x2 µm^3^ size at random locations. An expert annotator marked all spine heads based on morphology and post-synaptic densities, differentiating spines from pre-synaptic boutons by their different nanoscale topology of protein-dense features, and boutons typically tapering into axons at two locations. Spine density was 1.0±0.3/µm^3^ (mean±s.d.), consistent with previous data in the cortex (*1*). The annotator then manually attached spine heads to parent dendrites. Out of 306 spine heads contained in the randomly sampled volumes, 285 (93.1 %) could be unambiguously traced to a dendrite whereas 21 (6.9 %) were marked as uncertain. Overall, this confirmed high traceability of LICONN data and excluded the presence of a large population of non-traceable “orphan” spines.

To next test whether LICONN data allowed for volumetric annotation, we manually reconstructed neuronal structures in a 19.3 x 19.3 x 8.1 µm^3^ LICONN volume (imaged at 9.7 x 9.7 x 12.95 nm^3^ effective voxel size) from the CA1 *stratum oriens* in the hippocampus (**Fig. 1G, Movie S1**). We reconstructed 658 structures, revealing their complex shapes and the interwoven arrangement in the neuropil. This demonstrated the suitability of LICONN data for detailed volumetric annotation. However, manual reconstruction scales poorly, and would be difficult to achieve for the volume in **Fig. 1A**.

### Automated Segmentation with Flood-Filling Networks

Having manually validated traceability and segmentability, and with structural detail and voxel dimensions akin to EM connectomics data, we set out to analyze larger volumes by adopting deep-learning based segmentation algorithms developed for EM connectomics. Specifically, we trained Flood-Filling Networks (FFNs) (*48*), which have achieved state-of-the-art segmentation accuracy on a diverse set of connectomic datasets.

We imaged a 109 x 74 x 22 µm^3^ region from the hippocampal CA1 region (**Fig. 2A, fig. S17**) in a 4x6 tile arrangement with 9.7 x 9.7 x 13.0 nm^3^ effective voxel size. Within this volume, we chose an 83,825 µm^3^ bounding box (85 x 69 x 14 µm^3^, 68.6 Gigavoxel (Gvx) at native imaging resolution) and produced ground truth annotations constituting 540 axons with 17.8 mm cumulative path length (covering 920 Megavoxel (Mvx)) and 314 dendrites with 23.8 mm cumulative path length (3,344 Mvx) using an iterative procedure that interleaved model predictions and manual proofreading (see supplementary materials, materials and methods).

An FFN was trained on the ground truth annotations, applied to the whole bounding box (**Fig. 2**), and evaluated on a set of 99 manually skeletonized neurites (**figs. S18 and S19**)) consisting of 69 axons (1.8 mm cumulative path length) and 30 dendrites (1,041 dendritic spines). Mean spine density along dendrites (1.6±0.3/µm, evaluated on 9 dendrite stretches) in the manual skeletons was similar to previous EM data (*49*). We optimized the FFN base segmentation (see materials and methods) to minimize merge errors, the success of which was confirmed by comparison with the manually generated skeletons (0 mergers, 413 splits, 80.1% overall edge accuracy; a skeleton edge was counted as “accurate” when both its nodes were labelled identically and the label did not correspond to a segment overlapping more than one skeleton, see ref. (*48*) for details). We applied an automated agglomeration procedure (*48*), which increased edge accuracy to 92.8% (**Fig. 2H**), reducing splits by 92.5% (from 413 to 31) with some misattached spines but no major morphological merge errors (i.e., no mergers between axons or dendrites).

We then attempted to eliminate the remaining errors in the automated reconstruction through comprehensive manual proofreading of the entire 83,825 µm^3^ volume (correcting object-level split and merge errors, see materials and methods). We labelled objects in the proofread segmentation as either axon, dendrite, or glia using an automated classifier (see materials and methods) and found 18,268 axons with 342.3 mm cumulative path length (5.8% (by length) of which are displayed in **Fig. 2C**, for segmentation see https://neuroglancer-demo.appspot.com/#!gs://liconn-public/ng_states/expid82_fig2_axons.json), 1,643 dendrites (119.1 mm cumulative path length), and 71,269 dendritic spines (**Fig. 2D,E)**, 27.3% of which are displayed in **Fig. 2D** (for segmentation see https://neuroglancer-demo.appspot.com/#!gs://liconn-public/ng_states/expid82_fig2_dends.json). Using the manually generated skeletons as reference, we evaluated the contribution of the base segmentation, automated agglomeration, and manual proofreading to the increases in segment size and in edge accuracy (**Fig. 2G,H**). Comparing the proofread reconstruction with the manually traced skeletons yielded an overall edge accuracy of 95.6% and revealed one morphological merger (involving a dendrite pair), 29 incorrectly attached spines (2.8% of all spines), 12 uncorrected spine splits (1.1%), and zero splits involving axons or dendrite trunks. The volume furthermore contained glial segments and blood vessels which we did not process further, as the FFN models were trained exclusively on neuronal structures. Dense segmentation thus revealed the complex 3D arrangements of neuronal structures at nanoscale detail (**Fig. 2E,F, Movie S2**) and is available online in a browsable format together with the volumetric imaging data (https://neuroglancer-demo.appspot.com/#!gs://liconn-public/ng_states/expid82.json).

To further exclude the possibility that a substantial fraction of structures was systematically omitted from the automated segmentation, we merged all segments and overlaid these with the imaging data. Visual inspection of multiple regions suggested that signal-containing regions that were not covered by segments largely corresponded to slight deviations in segment shape. The small portion of areas not captured in the automated segmentation contained either intracellular regions or spines.

We therefore quantified the traceability of spines in the volume. First, we randomly sampled 40 subvolumes of 2 x 2 x 2 µm^3^. An annotator was able to trace 281 out of 301 spines (93.4%) to the parent dendrite, using only the raw imaging data. Spine density was 0.94±0.3/µm^3^, similar to previous reports in CA1 (*49*). To quantify missing spines in the FFN segmentation, we then compared ground truth skeletons to the segmentation and found that the remaining edge accuracy error was dominated by spines not labelled by the FFN (83 out of 1041 spines contained in the skeletons, 8.0%). Nearly all spine necks in the automated segmentation were attached to a dendrite; occasionally the FFN neglected to segment voxels corresponding to spine necks, but agglomeration and proofreading were in most cases still able to attach spine heads to the correct dendrite (manual proofreading focused on correcting topological split and merge errors rather than painting voxels). Measured spine head volumes (**Fig. 2I**, 53% within 0.01 and 0.05 µm^3^) were consistent with EM data (*50*) whereas the remaining predicted segments after agglomeration were mostly small irregular shapes (**fig. S19**).

Overall, LICONN enabled FFN-based segmentation with automated accuracy comparable to state-of-the-art EM results (*48*, *51*), as well as manual correction of remaining errors utilizing standard connectomic proofreading workflows.

### Molecularly annotated connectomics

We next sought to take advantage of light microscopy’s ability to visualize specific molecules to directly verify cellular, subcellular, and synaptic identities based on molecular components and to place those molecules in the context of the tissue’s 3D architecture and connectivity. Using post-expansion immunolabeling (applied here after the first expansion step) avoided additional processing of the tissue before expansion and promoted epitope accessibility (*27*, *34*) in the expanded, molecularly decrowded tissue-hydrogel hybrid. It also renders displacement of reporter fluorophores from biological target structures irrelevant by effectively “shrinking” antibodies from their physical size of ∼10 nm to ≲1 nm. While certain epitopes will be denatured by the expansion procedure, our limited screen identified a range of commercial antibodies compatible with the LICONN procedure. The iterative expansion paired with multicolor readout in standard spinning-disc confocal microscopes directly provided super-resolution measurements of structural and molecular channels, which is difficult to achieve when correlating light microscopy with EM connectomics (**fig. S20**).

To demonstrate molecularly informed connectomics with LICONN, we first focused on synaptic proteins. Specifically, we immunolabelled the pre-synaptic scaffolding protein Bassoon, a component of the cytomatrix at active zones, and PSD95, a scaffolding protein in post-synaptic densities of excitatory synapses and visualized them in the context of the structural channel in triple color measurements (**Fig. 3A**). At the high three-dimensional resolution achieved here, Bassoon labeling revealed a lattice-like arrangement of nanoscale spot-like structures, recapitulating the high-protein density features observed at pre-synapses in the structural channel (**Fig. 1C**). Both PSD95 and Shank2, another post-synaptic scaffolding protein, were arranged in more compact, disc-like arrangements that mirrored the post-synaptic densities in the structural channel. Distance measurements between Bassoon and Shank2 yielded 152 ± 17 nm (**Fig. 3B**, mean ± s.d., 106 synapses across *n*=2 animals), comparable to previous measurements on cultured neurons (*52*). Applying the FFN segmentation model from **Fig. 2A-F** enabled us to locate the synaptic molecular machinery in the context of their respective neuronal structures, such as a spiny dendrite rendered in **Fig. 3C**. We labelled additional synaptic proteins in LICONN’s super-resolved structural framework (**Fig. 3D, fig. S21**). As expected, the active zone markers Munc13-1 and RIM1/2 overlapped in space at the pre-synapse. We also visualized P/Q type Ca^2+^ channels (Ca_V_2.1) at pre-synapses and labelled for the vesicular glutamate transporter vGLUT1 to highlight synaptic vesicles, while co-labeling with RIM1/2 concomitantly demarcated the active zone. The distribution of the NMDA glutamate receptor GluN1 showed pronounced overlap with Shank2 at post-synapses, with the centres of mass of their 3D distributions 16.3 ± 4.6 nm (184 synapses, *n*=1 animal) apart (**Fig. 3E**).

### Computational interpretation of synaptic connectivity from molecular labels

Proximity between neurites is a weak predictor of synaptic connectivity (*1*). We therefore used molecular information as ground truth for connectivity and developed an automated synapse identification pipeline based on synaptic immunolabeling (**Fig. 3F-I, fig. S22**). We first computationally annotated pre-synapses and excitatory post-synapses as defined by the presence of Bassoon and Shank2, respectively (**Fig. 3F**). For this, sampling the intensity of the structural channel was useful to distinguish specific synaptic signal from unavoidable immunolabeling background (see materials and methods). We then developed a procedure to automatically match corresponding pre- and post-synaptic sites to full synapses, taking both classical one-to-one and one-to-many matches into account (**Fig. 3G**). Unpaired pre-synapses corresponded to inhibitory synapses, missed Shank2 detections, or infrequent synapses that lacked Shank2 (but that we were still able to identify as excitatory due to the presence of a prominent post-synaptic density). We therefore revisited unpaired pre-synapses and classified neighboring post-synapses as excitatory if a prominent post-synaptic density was present in the structural channel. When comparing purely automated synapse detections with manually validated synapse locations in a test dataset comprising 1059 excitatory synaptic connections, we obtained an accuracy of 95% for the detection of pre- and post-synapses, and 90% for fully assembled synapses (**Fig. 3H**) (F1: range 0-1, combining precision and recall; pre-synapses: F1=0.94; post-synapses: F1=0.95; full synapses: F1=0.90, 913 µm^3^ LICONN volume). We also found detection to be robust against variation of imaging parameters (**fig. S22**). Finally, by integrating FFN-based neuron segmentation with automated synapse detection, we inferred excitatory inputs from axons onto a specific dendrite **(Fig. 3I**). LICONN thus allowed mapping molecularly defined synaptic connectivity onto automated morphological reconstructions.

### Molecular identification of inhibitory vs. excitatory synapses

We then used specific immunolabels to molecularly identify and distinguish excitatory and inhibitory synapses in cellular reconstructions. To identify inhibitory synapses, we labelled for Gephyrin, a post-synaptic scaffolding protein. Gephyrin-positive, inhibitory post-synapses had less pronounced features in the structural LICONN channel than their excitatory, Shank2-positive counterparts and these molecules were mutually exclusive at any individual synapse (**Fig. 3J**). As a consistency check, we also co-labelled vesicular GABA transporter (vGAT) and Gephyrin in LICONN datasets, thereby identifying inhibitory pre- and post-synaptic compartments in the same synapses. As expected, pre-synaptic Bassoon and post-synaptic Gephyrin were closely juxtaposed (**Fig. 3K**). Specific molecular labeling thus distinguished excitatory and inhibitory synapses in LICONN volumes.

### Connectivity analysis

We now sought to relate molecular properties to neurite tracings in order to characterize fundamental parameters of synaptic wiring diagrams, such as inhibition and excitation. We imaged two LICONN volumes in primary somatosensory cortex (143 x 20 x 24 µm^3^ and 179 x 20 x 24 µm^3^, 1 animal), including immunolabeling for postsynaptic scaffolding proteins at excitatory (Shank2) and inhibitory (Gephyrin) post-synapses (**Fig. 4, figs. S23 and S24**). We manually analyzed 11 spiny dendrites (**Fig. 4A**, total length 123 µm, 328 dendritic spines) and found 2.9±1.1 synaptic inputs per µm length (**Fig. 4B**, including both Shank2 and Gephyrin positive), with higher density of Shank2-positive (2.6±1.1/µm) than Gephyrin-positive (0.3±0.2/µm) inputs. A large fraction of synapses was found on spine heads (90.2 %), whereas 1.6 % and 8.2 % were located at spine necks and shafts, respectively (**Fig. 4C**). As expected, the vast majority of spine heads were only positive for Shank2 (95.7±2.9 %), while rarely, a spine head had only a Gephyrin-positive connection (0.6%) or had a Gephyrin-positive connection in addition to the usual excitatory input (1.6%). A similarly small number of spine heads (2.1%) had neither a Shank2-nor a Gephyrin-positive connection but featured a PSD in the structural channel, corresponding to cases where either Shank2 was not expressed or not labelled (**Fig. 4D**). Synapses located at shafts were overwhelmingly inhibitory, i.e. positive for Gephyrin (93.5%), whereas the remaining 6.5% were positive for Shank2. As expected, these molecules were mutually exclusive at any given synaptic site (**Fig. 4E**). Also, the overall balance of excitatory vs. inhibitory inputs to dendrites was straightforward to determine based on immunolabeling (90% vs. 10%, respectively, **Fig. 4F**). Results were consistent when repeating the analysis utilizing exclusively structural information, i.e. synapse location (shaft, spine head, neck) of axon outputs for classifying axons into excitatory vs. inhibitory and determining excitation/inhibition balance, similarly as commonly done in EM-based connectomics (**fig. S24**).

**Fig. 4.**
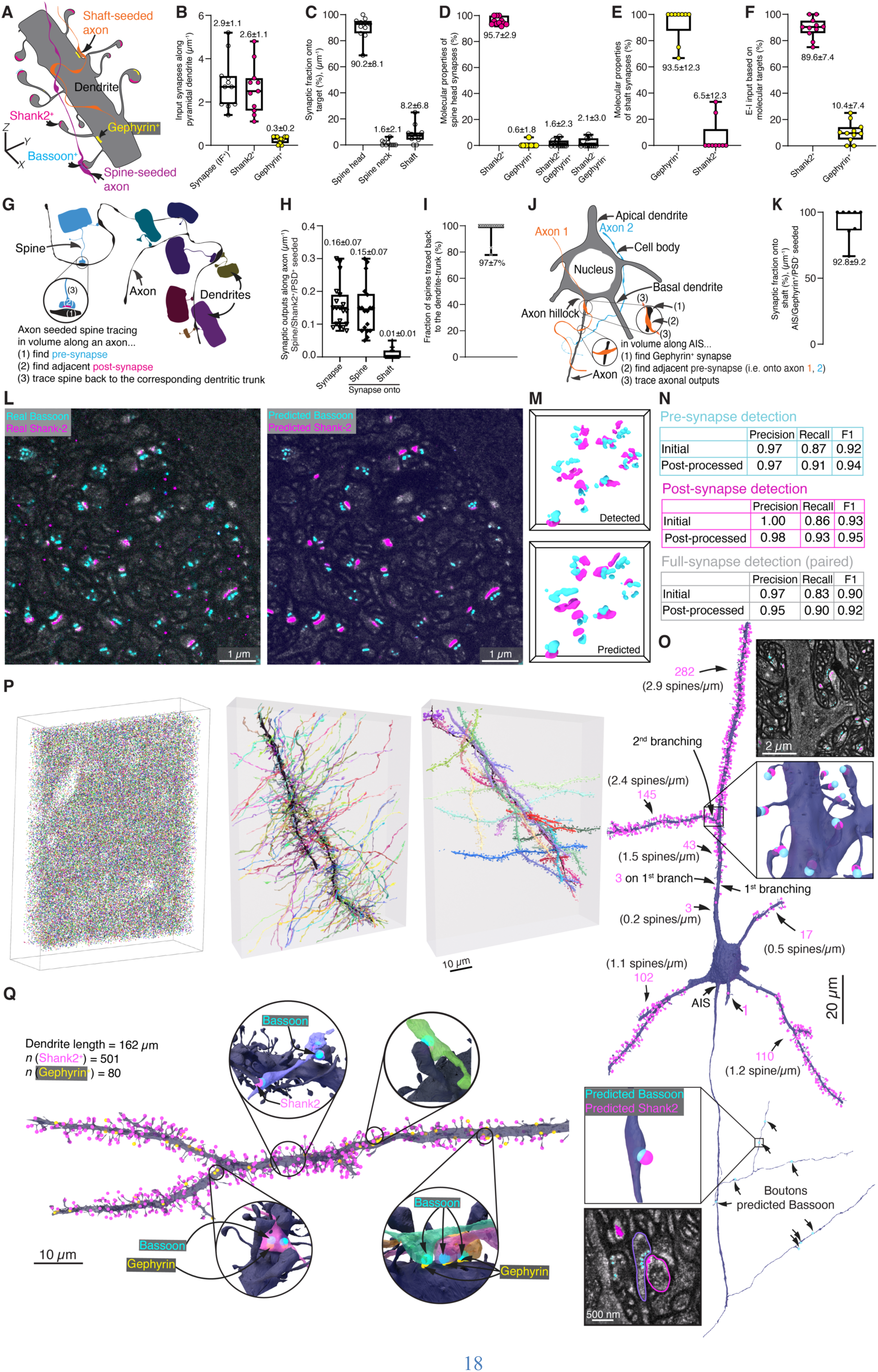
Connectivity analysis and deep-learning based synapse detection. (**A**) Schematic of spiny dendrite with Shank2-expressing excitatory (magenta) and Gephyrin-expressing inhibitory post-synapses (yellow), as well as connected axons (pre-synaptic Bassoon: cyan), with seed locations for axon selection. (**B**) Density of synaptic inputs onto spiny dendrites per unit length, as determined by immunolabeling for excitatory (Shank2 positive) and inhibitory (Gephyrin positive) synapses (IF^+^: immunofluorescence positive). Total density of synapses as well as excitatory and inhibitory synapse density analyzed in two LICONN datasets (total 154,560 µm^3^, 1 animal) from primary somatosensory cortex. Box plot with median, lower and upper quartiles and mean±s.d.. Data points represent individual dendrites throughout (11 dendrites, 123 µm total, 351 synapses). (**C**) Distribution of synaptic input location of spiny dendrite according to spine head, spine neck, or shaft location (all synaptic connections, as defined by the presence of immunolabeling for Shank2 or Gephyrin). (**D**) Molecular properties of synapses located at spine heads, as identified by immunolabeling and structural features (PSD). ∼96% expressed Shank2; in rare cases (0.6%), we observed a spine head that had only a Gephyrin-positive synapse. In 1.6%, both a Shank2-positive synapse (with PSD) and a Gephyrin-positive (without PSD) synapse were present on the same spine head. In 2.1% of spine heads, a synaptic connection was present as judged by morphology and the presence of a PSD, but neither Shank2 nor Gephyrin were expressed or labelled. (**E**) Relative proportion of Gephyrin- and Shank2-positive synapses at dendritic shaft locations. (**F**) Relative abundance of excitatory and inhibitory synaptic inputs onto individual spiny dendrites (excitation-inhibition balance), as defined by immunolabelings for Shank2 and Gephyrin. (**G**) Schematic for selection of spines, seeded by the outputs on an excitatory axon. To select excitatory axons, an arbitrary spine head in the volume (Shank2 and PSD positive) was selected, a connected axon was identified, and the axon traced. All outputs of the axon were evaluated. (**H**) Outputs of spine-seeded axons (density/µm axon length), including synapses onto spines and shafts (21 axons with more than 3 outputs, representing individual data points, total length 990 µm). (**I**) Axon-seeded spine tracing. Fraction of spines that can be traced to the parent dendrite when selecting all spines that are contacted by the same set of axons (21 axons, 128 spines). Datapoints represent individual axons. (**J**) Schematic for selection of inhibitory axons seeded at axon initial segment (AIS) (Gephyrin positive, no PSD). (**K**) Fraction of synaptic outputs onto shaft locations of AIS-seeded, inhibitory axons (8 axons with more than 3 outputs, representing individual data points). (**L**) Deep-learning based prediction of Bassoon and Shank2 location from structural LICONN channel. Single plane from a volumetric LICONN dataset in CA1 stratum radiatum, not included in the training, comparing predicted Bassoon and Shank2 with the respective immunolabeling channels. (**M**) Volumetric renderings of predicted Basson and Shank2 distributions compared to corresponding immunolabeling. (**N**) Precision, recall, and F1 score for deep-learning-based prediction of pre-synapses, post-synapses, and fully assembled synapses, evaluated using manually generated ground truth (180 µm^3^ test dataset, see **fig. S22** for additional test dataset). Metrics are given both for the base prediction and with a post-processing step (see materials and methods). (**O**) Excitatory synaptic input and output fields for a pyramidal neuron from the cortical dataset in **Fig. 1A**. Synapse detections based on prediction of Bassoon and Shank2 locations mapped onto an FFN segmentation. Numbers in magenta indicate detected synapses between indicated dendrite branch points. Magnified views show structural LICONN channel with predictions of Bassoon and Shank2 and corresponding cellular segments with synaptic detections. Partial proofreading applied (eliminating false positive detections without manually adding missed detections). (**P**) Prediction of synaptic connectivity in the hippocampal dataset in **Fig. 2A**. *Left*: Rendering of all synaptic predictions. *Middle*: Single axon with synaptically connected dendrites. *Right*: Single dendrite with synaptically connected axons. (**Q**) Integration of structural, immunolabeling, and deep-learning analysis. Rendering of a dendrite from FFN segmentation of one of the LICONN volumes in panels A-K in primary somatosensory cortex with detection of excitatory (Shank2, 501 synapses) and inhibitory (Gephyrin, 80 synapses) post-synapses based on immunolabeling. Insets show pre-synaptic partners identified from additional deep-learning based prediction of Bassoon.

We then classified axons as excitatory according to structural and molecular characteristics, using the following criteria: we selected a spine in the volume and identified an axon synaptically connected to it (“spine-seeded”), by requiring both (i) a PSD in the structural channel and (ii) Shank2 at the synaptic seed site. When we manually traced 21 such axons (total length 990 µm), they displayed 0.16 ± 0.07 synaptic outputs per µm length (**Fig. 4G,H**), with 0.15±0.07/µm onto spines and 0.01±0.01/µm onto shaft locations, recapitulating the known preference of excitatory outputs for spines (*10*). We used these axons as seeds for a further independent test of spine traceability (“axon-seeded” spine tracing) in LICONN volumes, again allowing us to trace the vast majority of spine necks (96 %, 123 out of 128) to the parent dendrite (**Fig. 4I**).

For inhibitory axons (defined by (i) seeding from a synapse at a dendritic shaft, (ii) lack of a PSD and (iii) presence of Gephyrin), we found an inverse target preference (**fig. S24I,J**). Finally, we selected 3 cells for which the entire axon initial segment (AIS) (*53*) was contained in the imaging volumes (**Fig. 4J**). We observed that AIS were characterized by a pronounced periodic labeling pattern in the structural channel, akin to the actin/spectrin-induced lattice observed in neurites (**Fig. 1D, 5A**), extending over 43.8±1.3 µm. We detected 0.25±0.1 Gephyrin-positive inputs per µm AIS. We selected axons that provided inhibitory input to AIS (Gephyrin positive, no pronounced PSD) and traced their outputs. The 8 axon stretches analyzed with 4 or more outputs formed a total of 46 synapses at dendrites (synapses onto soma or AIS not analyzed), with a strong preference for shaft locations (93%, **Fig. 4K**).

LICONN thus provided a natural means to integrate structural and molecular information to derive fundamental neuronal network parameters related to excitatory and inhibitory connectivity.

### Deep-learning based prediction of synapse locations

To facilitate circuit analysis on a larger scale and to overcome limitations in microscopy hardware and imaging time introduced by adding color channels for further molecular targets, we leveraged the correspondence between molecular features and the local micro- and nano-architecture as represented in the structural channel to predict, rather than measure, synaptic molecule locations. Using deep learning methods, we predicted both Bassoon at pre-synaptic sites and Shank2 at excitatory post-synapses (**Fig. 4L**). Instead of relying on human annotation of synaptic structures, we used pre-processed, super-resolved measurements of Bassoon and Shank2, paired with the structural LICONN channel, as training data (16.699 µm^3^ containing ∼16.250 excitatory synapses). We implemented convolutional neural networks (*26*, *54*) to predict molecule locations from the structural LICONN channel. When evaluated on datasets not included in the training, predicted signals were highly consistent with ground truth immunolabeling data (**Fig. 4L,M**). We computationally converted molecule predictions to pre- and post-synaptic annotations and implemented a similar post-processing step (see materials and methods) as for immunolabeling-based detection to increase accuracy. Comparison of deep-learning based synapse prediction with human ground truth annotations (generated by taking both the immunolabeling and structural LICONN channels into account) resulted in F1 scores >0.9 for detection of synapses (**Fig. 4N**) (pre-synapses: F1=0.94; post-synapses: F1=0.95; full synapses: F1=0.92, using 14.207 µm^3^ (∼13.650 synapses) for training and 180 µm^3^ for testing; see **fig. S22** for application in additional test dataset with different imaging parameters). This demonstrated that excitatory synapse locations can be predicted with high fidelity from structural LICONN data.

We now applied deep-learning based synapse prediction to map the synaptic input field of an identified neuron in a dataset devoid of immunolabeling. Specifically, we chose a pyramidal neuron from the FFN-based segmentation of the cortical dataset in **Fig. 1A** and related both pre-synapse (Bassoon) and excitatory post-synapse (Shank2) predictions to the cellular segmentation (**Fig. 4O, Movie S3**). The imaging volume contained 821 µm of this cell’s neurites. On dendrites (454 µm), we detected 705 excitatory synaptic inputs (1.6/µm dendrite, 475 on apical and 230 on basal dendrites). The volume also contained 367 µm of the neuron’s axonal output, featuring 8 synaptic output predictions. We then sought to analyze overall connectivity patterns in a LICONN volume devoid of immunolabeling. We applied dual prediction of Bassoon and Shank2 to map excitatory synaptic connectivity to the densely proofread neuronal segmentation of the hippocampal volume in **Fig. 2A-F**. We detected 79,291 pre-synapses (density: 0.95/µm^3^) and 71,976 excitatory post-synapses (density: 0.86/µm^3^) and mapped them onto individual neurite segments (**Fig. 4P**). Singling out one axon with its synaptic partners illustrated its rich connectivity, with 31 synapses onto 28 different dendrites. Similarly, when focusing on a single spiny dendrite, we identified 284 axons making a total of 303 synaptic contacts with it, most of which (267) were single connections, whereas 16 axons made 2 synapses and one made 4 synapses (**Fig. 4P**).

We then created a “virtual” 4-color connectomic volume combining the LICONN structural channel, deep-learning based prediction of Bassoon (for detection of pre-synaptic elements), and super-resolved immunolabeling detection of Gephyrin and Shank2 (for molecular differentiation of synapses). With this, we were able to map synaptic inputs onto a dendrite and distinguish excitatory vs. inhibitory synapses (**Fig. 4Q**).

### Identification of cell types and subcellular structures across diverse brain areas

We now sought to leverage both structural and molecular information in LICONN to characterize cells in terms of their types and subcellular specializations. Similar to EM connectomics, analysis of cell shape enabled an expert to classify cells in the cortical dataset in **Fig. 1A** (79 cells analyzed) as pyramidal neurons (65 cells, 82.3%), interneurons (9 cells, 11.4%), and glial (5 cells, 6.3%). Relative proportions were comparable to EM data from mouse visual cortex (*55*). Adding specific immunolabeling molecularly identified cell types, such as inhibitory interneurons expressing parvalbumin or somatostatin (**Fig. 5A**).

**Fig. 5.**
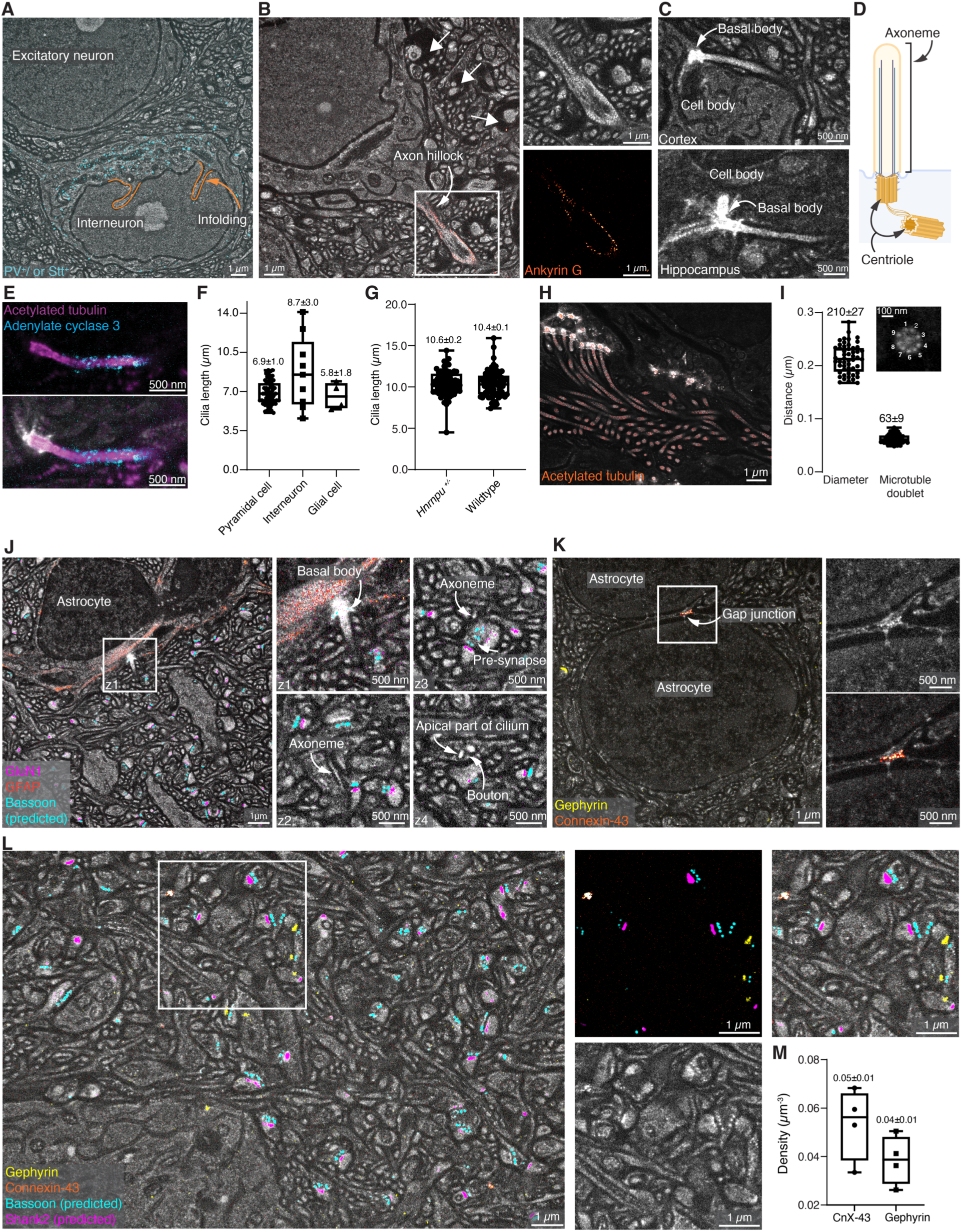
LICONN applications beyond synaptic connectivity. (**A**) Molecular identification of an interneuron via combined immunolabeling (cyan) for parvalbumin and somatostatin. Single plane of LICONN volume in cortex, overlaid with maximum intensity projection of immunolabeling. Prominent infoldings of the interneuron’s nucleus are indicated in orange. Images are representative of *n*=2 immunolabeling replicates in one animal. (**B**) Overlay of Ankyrin G immunolabeling (red) with LICONN structural channel (gray) in cortex, highlighting axon initial segment. Magnified views show immunolabeling and structural channels separately. The periodic Ankyrin G labeling pattern coincides with periodic modulation of protein density in the structural channel. Axons encircled by low-protein density voids (arrows) correspond to myelinated axons (**fig. S25**). Ankyrin G immunolabeling was replicated *n*=3 times. (**C**) Primary cilia in LICONN volumes from cortex and hippocampus (single planes displayed). Centrioles in the basal body are intensely labelled in LICONN. (**D**) Schematic of primary cilia. (**E**) Immunolabeling for acetylated tubulin (magenta) and adenylate cyclase 3 (cyan) at primary cilium in LICONN volume in hippocampal CA1, with the membrane-bound adenylate cyclase signal ensheathing the axoneme. *Top*: immunolabeling. *Bottom*: overlay with structural channel. Single plane displayed. Immunolabelings were replicated *n*=2 times. (**F**) Length of primary cilia according to cell type in the LICONN dataset in **Fig. 1A** (79 cells, data points in box plot (median, lower and upper quartiles, mean±s.d.) represent individual cells). (**G**) Length of primary cilia in the hippocampal CA1 pyramidal layer (**fig. S26**) in wild type (78 cells from 3 technical replicates in each of *n*=3 animals) and *Hnrnpu^+/-^* haploinsufficient mice (80 cells from 3 technical replicates in each of *n*=3 animals). (**H**) Rendering of LICONN volume at the border between the corpus callosum and alveus (**fig. S30**), revealing cells with multiple cilia. Structural channel (gray) overlaid with immunolabeling for acetylated tubulin (red). Representative of replicates in *n*=2 animals. (**I**) Cross section of a cilium with 9-fold symmetry in protein density, likely reflecting the arrangement of microtubule doublets. Box plots indicate the diameter of the microtubule doublet ring and the distance between doublets (median plus upper and lower quartile, 52 cross sections measured across 2 technical replicates in *n*=1 animal (5 imaging volumes)). (**J**) LICONN volume (gray, single plane displayed) in hippocampus with immunolabeling for glial fibrillary acidic protein (GFAP, red) highlighting astrocytes and immunolabeling for the glutamate receptor subunit GluN1 (magenta). Additional deep-learning based prediction of Bassoon location (cyan). Magnified views of the astrocyte’s primary cilium at different z-planes (z1-z4), in close apposition to synaptic boutons. Immunolabelings representative of *n*=2 replicates. (**K**) Molecular identification of gap junctions and inhibitory synapses. Single plane from LICONN volume (gray, cortex) with immunolabeling for astrocytic gap junction protein Connexin-43 (orange) and for the inhibitory synapse marker Gephyrin (yellow), with a magnified view of a gap junction between two astrocytes. Representative of *n*=3 replicates. (**L**) “Virtual” 5-color measurement in cortical neuropil. LICONN structural data (gray) overlaid with immunolabeling for astrocytic gap junction protein Connexin-43 (orange) and inhibitory post-synapses (Gephyrin, yellow), as well as deep-learning based prediction of specific molecules at pre-synapses (Bassoon, cyan) and excitatory post-synapses (Shank2, magenta). Immunolabeling plus prediction and structural channels are shown separately for the boxed region. Immunolabelings representative of *n*=3 replicates. (**M**) Density of Gephyrin-positive inhibitory synapses and Connexin43 (CnX-43)-positive gap junctions (mean, *n*=4 imaging volumes recorded across cortex and hippocampal CA1).

We then used molecular labeling to clarify the identity of subcellular structures and characterize their appearance in the structural channel. Labeling for Ankyrin G molecularly highlighted axon initial segments (AISs). We discovered that at the high resolution employed here, Ankyrin G displayed a lattice-like pattern at AISs, mirrored by a similar pattern in the protein density map (**Fig. 5B**). This differentiated AISs from dendrites which were largely devoid of protein density stripes near the soma. Similarly, labeling for myelin basic protein helped us clarify the identity of halo regions of low protein density around certain axons (**Fig. 5B**), unambiguously identifying them as myelinated axons (**fig. S25**).

Primary cilia (**Fig. 1B, 5C-G**) are important signaling hubs on many neuronal and non-neuronal cells whose connectivity and function in the brain are being intensely explored (*56*, *57*). We corroborated their identity with immunolabeling for acetylated tubulin, a common posttranslational modification of stable microtubules, and adenylate cyclase 3 (**Fig. 5E**). We quantified cilia length according to cell type (**Fig. 5F**) from manually generated skeletons in 79 cells in the cortical dataset in **Fig. 1A** (animal aged 2 months, 65 pyramidal neurons: 6.9 ± 1.0 µm; 9 interneurons: 8.7 ± 3.0 µm; 5 glial cells: 5.8 ± 1.8 µm). Ciliary function is essential for many aspects of brain development, with e.g. primary ciliary dyskinesia being associated with agenesis of the corpus callosum (*58*). In humans, mutations in the Heterogeneous nuclear ribonucleoprotein U (*HNRNPU*) gene cause early-onset epilepsy, autistic features, and intellectual disability. In the developing mouse brain, *Hnrnpu* has been previously localized on primary cilia (*59*). Additionally, changes in the expression of genes associated with cilia organization have been observed in a human model system of induced pluripotent stem-cell derived neurons obtained from patients with mutations in *HNRNPU* (*60*). We hypothesized that mutations in *Hnrnpu* may affect primary cilia length and tested it in a haploinsufficient (*Hnrnpu^+/-^*) mouse model. To showcase how LICONN can provide quantitative data for tissue phenotyping, we skeletonized primary cilia of pyramidal neurons in the hippocampal CA1 region (**fig. S26**) for both *Hnrnpu+/-* (80 cells from *n*=3 animals, with 3 technical replicates in each) and wild type (78 cells from from *n*=3 animals, with 3 technical replicates in each) animals (**Fig. 5G**). This analysis allowed us to conclude that, at least for hippocampal neurons, *Hnrnpu* mutations do not result in obvious primary cilium defects (length wild type: 10.4±0.1 µm, length mutant: 10.6±0.2 µm, no statistically significant difference).

Finally, we applied LICONN more broadly throughout the brain to assess tissue characteristics beyond synaptic connectivity. We confirmed applicability in various regions, including the hypothalamus, piriform area, and cerebellum (**fig. S27**), and imaged the characteristic organization of the hippocampal CA3 stratum lucidum (**fig. S28**). When applying our workflow to white matter, in the region of the corpus callosum, a paramount structure connecting the two hemispheres, and the alveus, we found layers with a high density of myelinated axons (**fig. S29**). We were intrigued by nearby clusters of cells with multiple prominent processes (**Fig. 5H, fig. S30**). Investigating their ultrastructure (**Fig. 5I**), we realized that they corresponded to multiple cilia, originating from basal bodies, again with nine-fold internal symmetry. Cilia were prominently highlighted when immunolabeling for acetylated tubulin (**Fig. 5H**). The microtubule ring had a diameter of 210 ± 27 nm (mean±s.d) and individual doublets were spaced 63 ± 9 nm apart (**Fig. 5I**, 52 cross sections analyzed across 2 technical replicates in *n*=1 animal).

### Molecular identification of gap junctions

The ability to map electrical connectivity via gap junctions would be especially useful, as this information is typically missing from connectomics datasets because their reliable detection requires particularly high resolution in EM (*61*). These intercellular channels contribute to functional network properties in addition to chemical synapses. We immunolabelled Connexin-43, a gap junction protein expressed in astrocytes (*62*) and identified astrocytes by their expression of glial fibrillary acidic protein (GFAP) (**Fig. 5J,K**). LICONN visualized both the electrical connections and cellular partners. Labeling for GFAP also molecularly assigned cellular structures associated with capillaries to astrocytes (**fig. S25**). Immunolabeling of Connexin-43 and Gephyrin allowed us to visualize electrical and inhibitory connections. Complementing this with excitatory synaptic connections via deep-learning based prediction of Bassoon and Shank2 (**Fig. 5L**) showcased simultaneous mapping of excitatory, inhibitory, and electrical connectivity in the same circuit. Similarly, specific immunolabeling allowed us to determine the density of inhibitory (0.038±0.008/µm^3^, Gephyrin), and electrical (0.053±0.013/µm^3^, astrocyte-associated Connexin-43) connections (4 imaging volumes across hippocampus and cortex, 2 each) (**Fig. 5M**).

### Lossless axial extension of imaging volumes

To axially extend LICONN volumes beyond the imaging depth accessible with high-NA objective lenses (0.6 mm working distance in our case), we implemented a block-face imaging method to obtain LICONN volumes built from partially overlapping subvolumes arranged on a 3D grid, which allowed seamless fusion without gaps (**fig. S31**). As before, we first imaged partially overlapping volumetric tiles arranged in 2D to obtain a first slab of LICONN data. We then used a conventional vibratome to slice off most of the imaged hydrogel layer, but placing the cut still within the imaged slab. In a subsequent imaging round, we thus obtained an axially offset multi-tile volume situated more deeply in the tissue while featuring a continuous region of axial overlap with the first multi-tile volume. We then computationally fused the individual imaging volumes in 3D in a voxel-exact manner (see materials and methods), enabling tracing of neuronal structures, including thin axons, across imaging slabs (**fig. S31**; original data for volume fused in 3D is available online in browsable format at: https://neuroglancer-demo.appspot.com/#!gs://liconn-public/ng_states/expid146.json).

Taken together, LICONN is a straightforward technology that directly provides integrated structural and molecular characterization across brain regions, cell types, and spatial scales.

## Discussion

The development of connectomics methods has been driven by the challenge of simultaneously achieving dense 3D reconstruction of neurites, synapse-level connectivity, diverse molecular annotations, and cost-effective scaling to large tissue volumes (e.g. cubic millimeters or more). Successive generations of EM-based technologies have produced enormous progress (*2*), but important limitations remain, particularly in the ability to extract molecular details of tissue. In contrast, current light microscopy techniques for visualizing the cellular architecture of brain tissue (*24–26*, *30*, *37*), including LIONESS (*25*) and CATS (*26*), do not reach the accuracy and traceability required for connectomic reconstruction. Here we introduce LICONN which, like EM, enables reliable manual 3D tracing of dendrites, fine axons, and spines (**Fig. 1**), high accuracy automated 3D reconstruction of those structures (edge accuracy 92.8% and no major morphological merge errors, **Fig. 2**), and reliable detection of chemical synapses (F1>0.9 for either immuno-labeling or deep-learning based detection, **Fig. 3,4**). However, unlike EM, LICONN also enables direct and simultaneous measurement of spatially resolved molecular information, including specific proteins that reveal chemical synapse subtypes (**Fig. 3**), electrical synapses, and important subcellular features (**Fig. 5**).

We applied LICONN to acquire volumes up to roughly one million cubic microns in size (native tissue scale, **Fig. 1A**), which is the same scale as previous EM-based connectomics datasets, sufficiently large for biological analysis of mammalian neural circuits (*1*, *10*, *63*, *64*) and achieved reasonable acquisition rates (6.5h acquisition time for ∼1 million cubic microns native tissue scale, 0.39 Teravoxel, effective voxel rate of 17*10^6^ voxels/s (17 MHz) including overhead due to tile overlap and sample stage positioning) even without dedicated optimizations for imaging speed (*25*, *65*). However, the largest EM datasets span a cubic millimeter, and a major long-term ambition in the field is mapping the hundreds of cubic millimeters encompassing an entire mouse brain (*66*). One promising strategy for scaling LICONN volumes includes hydrogel sectioning to parallelize readout and remove limitations in axial imaging range due to the finite working distance of the objective lens. LICONN hydrogels are mechanically robust, which facilitates handling and sectioning. We typically chose to expand slices of 50 µm thickness for experimental convenience but this does not constitute a fundamental limitation, and other expansion technologies have been scaled to entire mouse brains (*67*). We expect that scaling the LICONN expansion procedure to larger samples will be possible adjusting denaturation, polymerization, and labeling parameters. For example, when slightly adapting hydrogel expansion for 300 µm thick sections (materials and methods, **fig. S31**), we obtained equivalent LICONN imaging results. As one immediate route towards scaling, we further demonstrated that consecutive, overlapping imaging volumes in axial direction can be obtained and computationally fused, removing slabs of the expanded LICONN hydrogel already imaged. Computational analysis of much larger imaging volumes will also be feasible, with the tools for volume fusion (SOFIMA) (*45*) and automated segmentation (FFNs) (*48*) having been applied to petascale EM volumes (*9*), and our synapse detection pipeline implemented using tools (*68*) designed to support processing of large imaging volumes.

We have shown detection of specific proteins based on immunolabeling. By using post-expansion labeling, LICONN benefits from improved epitope access (*27*, *34*) and avoids fluorophore-to-epitope linkage error. Further refinement will expand LICONN’s ability to produce molecularly informed connectomes. For example, an additional important molecular labeling task will be readout of gene expression via combinatorial labeling. Hydrogel-compatible spatial transcriptomics methods have emerged (*69–72*) that measure gene expression directly in tissue. By integrating connectivity with in-situ molecular information from individual cells, LICONN presents a viable path towards multimodal descriptions of mammalian brain cells, including morphology, connectivity (including electrical connections), physiology, and gene expression (*73*, *74*).

We also applied LICONN to quantify cellular properties beyond synaptic connectivity, focusing on cilia. Analyzing primary cilia in a neurodevelopmental disease model exemplified how LICONN can be used to study genotype-to-phenotype relationships and cellular alterations in diseased brains. More generally, LICONN was developed to address arguably the most complex situation in which to reconstruct tissue structure, specifically identification and tracing of the finest neuronal processes such as axons and spines for connectomics; therefore we expect the procedure to be broadly useful in other organs and systems in which high-resolution tissue analysis is desirable and in many cases less demanding.

Finally, we note that LICONN is driven by conventional light microscopy hardware (in our case, spinning disc confocal) that is broadly available in biological laboratories, and, while LICONN sample preparation introduces novel strategies to achieve high-fidelity tissue expansion, the protocol is not fundamentally more complex to perform or reproduce compared to many previous expansion techniques that have been widely adopted. Deep-learning based analysis was critical to the success of the effort. The flood filling networks and deep-learning frameworks we applied for neuronal segmentation and synapse prediction, respectively, have been previously validated in EM connectomics. These and custom code we developed here are available open source. Taken together, LICONN forms the technological basis for routine adoption of connectomic studies in non-specialized neuroscience labs as well as high-resolution studies in further organs.

## Acknowledgments

We thank Sven Dorkenwald and Peter Li for critical reading of the manuscript. We acknowledge expert support by ISTA’s scientific service units: Imaging and Optics, Lab Support, Scientific Computing, Preclinical Facility, Miba Machine Shop, and Library.

## Author contributions

JGD and MRT conceived and initiated the study. MRT conceptualized, developed and applied expansion methodology, performed imaging and analysis, manual annotation and proofreading, visualized data and prepared figures. JL developed and performed computational detection and deep-learning prediction of synapses, analyzed connectivity, processed, analyzed, and visualized data, and prepared figures. VJ and MJ developed FFN based segmentation methods, applied by MJ. MJ analyzed segmentations, developed volume fusion methods and processed data. MRT, NA, JV performed manual tracing. AC performed manual neurite annotations. CK supported experiments. VV performed distortion analysis. CS supported image analysis. GN and BO generated *Hnrnpu*^+/-^ mice and provided tissue from mutant mice. JGD supervised the study, and designed experiments and analyses. JGD wrote the paper with MRT, VJ, JL, and MJ. All authors read and approved the manuscript.

