## Supplementary Materials for "Light-microscopy based dense connectomic reconstruction of mammalian brain tissue"

### Materials and Methods

Animals

Animal procedures were performed in accordance with national law (BGBLA 114 and Directive 522), European Directive 2010/63/EU and institutional guidelines for animal experimentation and were approved by the Austrian Federal Ministry for Education, Science and Research (authorizations BMBWF-V/Sb: 2020-0.363.126, 2021-0.547.215, 2021-0.842.237, 2022-0.121.445 and 2023-0.930.355).

Animals were housed in groups of 3-4 animals under controlled laboratory conditions (12:12 h light/dark cycle with lights on at 07:00 a.m.; 21 ± 1 °C; 55 ± 10 % humidity) with food (pellets, 10 mm) and autoclaved water *ad libitum*. Animals were housed in commercially available individually ventilated cages (IVCs) made from Polysulfone with a solid cage floor, dust free bedding (woodchips) and nesting material.

For all experiments, male and female mice were used interchangeably to demonstrate the technology. Adult mice (aged typically 2-3 months, unless otherwise noted) were used as indicated with the following genotypes: C57BL/6J wild type mice, *Thy1-eGFP* (STOCK Tg(Thy1-eGFP)MJrs/J mice, #007788, RRID:IMSR_JAX:007788 hemizygous), and haploinsufficient *Hnrnpu*$^{+/-}$ mice (deletion of one allele of *Hnrnpu* spanning axons 4 to 14, generated by crossing the *HnrnpU*$^{wt/flox}$ line (Hnrnpu<tm1.1Tman>/J, Strain:#032187, RRID:IMSR_JAX:032187) with the *CMV-Cre*$^{Cre/Cre}$ line (B6.C-Tg(CMV-cre)1Cgn/J, Strain #:006054, RRID:IMSR_JAX:006054).

Reagents

| Compound | Abbreviation | Vendor | Identifier |
|---|---|---|---|
| Acrylamide | AA | Sigma-Aldrich/Merck | A9099 |
| Sodium acrylate | SA | AK Scientific | R624 |
| N,N′-Methylenebisacrylamide | BIS | Sigma-Aldrich/Merck | M7279 |
| Ammonium persulfate | APS | Sigma-Aldrich/Merck | A3678 |
| N,N,N',N'- tetramethyl-ethylenediamine | TEMED | Sigma-Aldrich/Merck | T9281 |
| 4-Hydroxy-2,2,6,6-tetramethylpiperidin-1 | TEMPO | Sigma-Aldrich/Merck | 176141 |
| Sodium dodecyl sulfate | SDS | Sigma-Aldrich/Merck | 436143 |
| Sodium chloride | ---- | Sigma-Aldrich/Merck | 71383 |
| Tris 1 M, pH 8.0, RNase-free | ---- | ThermoFisher Scientific/ Invitrogen | AM9856 |
| Paraformaldehyde | PFA | Sigma-Aldrich/Merck | 158127 |
| Glycine | ---- | Sigma-Aldrich/Merck | 50046 |
| Sodium azide | ---- | Sigma-Aldrich/Merck | 71289 |
| Ketamine 100 mg/ml | ---- | MSD Tiergesundheit | NA |
| Xylazine 20 mg/ml | Xylasol | Livisto | NA |
| Novalgine 0.5 mg/ml | Metamizol | Sanofi | NA |
| Isoflurane | ---- | Virbac/ Vetflurane | NA |
| Glycerol triglycidyl ether (Glycidyl Glycerol-Ether, Polyfunctional) | TGE | Polysciences Europe GmbH | 09221 |

| | | | |
|---|---|---|---|
| Glycidyl Methacrylate (Glycidyl Acrylate) | GMA | TCI | G0497 |
| Sodium bicarbonate | $NaHCO_3$ | Sigma-Aldrich/Merck | 792519 |
| Tween-20 | Tween | Fisher BioReagents | BP337 |
| Sodium hydroxide | NaOH | Sigma Aldrich/Merck | S5881 |
| Hydrochloride | HCl | --- | NA |
| Poly-L-Lysine Hydrochloride | PLL-HCl | Sigma-Aldrich/Merck | P2658 |
| Acrylic acid N-hydroxysuccinimide ester | NAS | Sigma-Aldrich/Merck | A8060 |
| 6-((acryloyl)amino)hexanoic acid, succinimidyl ester | AcX | ThermoFisher Scientific | A20770 |
| 4’6-diamidino-2- phenylindole | DAPI | Sigma-Aldrich/Merck | D9542 |

**Table 1: List of chemicals**

| Compound | Abbr. | Vendor | Identifier | Working concentration |
|---|---|---|---|---|
| N-hydroxysuccinimidyl-ester Alexa Fluor 488 | NHS-AF 488 | Jena Bioscience (Click Chemistry Tools, Vectorlabs) | APC-002-1 (Click Chemistry Tools: 1338) | 40 µM |
| N-hydroxysuccinimidyl-ester Alexa Fluor 546 | NHS-AF 546 | Thermo Fisher Scientific | A20002 | |
| N-hydroxysuccinimidyl-ester Alexa Fluor 594 | NHS-AF 594 | Jena Bioscience | APC-004-1 | |
| N-hydroxysuccinimidyl-ester Atto 488 | NHS-Atto 488 | Atto-Tec | AD 488-31 | |

**Table 2: List of NHS-coupled fluorophores**

| Target, host species | Abbr. | Vendor | Identifier | Clonality | Working dilution |
|---|---|---|---|---|---|
| Anti-Bassoon antibody, Mouse | Anti-Bsn | Synaptic Systems | 141 011 | Monoclonal | 1:300 |
| Anti-Bassoon antibody, Rabbit | Anti-Bsn | Synaptic Systems | 141 003 | Polyclonal | 1:300 |
| Anti-RIM-1/2 antibody, Guinea pig | Anti-RIM-1/2 | Synaptic Systems | 140 205 | Polyclonal | 1:300 |
| Anti-Munc13-1 antibody, Rabbit | Anti-Munc13-1 | Synaptic Systems | 126 103 | Polyclonal | 1:300 |
| Anti-vesicular glutamate transporter 1 antibody, Rabbit | Anti-vGlut1 | Synaptic Systems | 135 302 | Polyclonal | 1:300 |
| Anti-vesicular gamma-aminobutyric acid transporter antibody, Rabbit | Anti-vGAT | Synaptic Systems | 131 003 | Polyclonal | 1:200 |
| Anti-$Ca^{2+}$ P/Q antibody, Guinea pig | Anti- $Ca^{2+}$ P/Q | Synaptic Systems | 152 205 | Polyclonal | 1:300 |
| Anti-NMDA type glutamate receptor antibody, Mouse | Anti-GluN1 | Synaptic Systems | 114 011 | Monoclonal | 1:300 |
| Anti-Postsynaptic density-95 antibody, Mouse | Anti-PSD95 | ThermoFisher/ Invitrogen | MA1-046 | Monoclonal | 1:400 |
| Anti-Shank2 antibody, Guinea pig | Anti-Shank2 | Synaptic Systems | 162 204 | Polyclonal | 1:300 |
| Anti-Shank3 antibody, Guinea pig | Anti-Shank3 | Synaptic Systems | 162 304 | Polyclonal | 1:300 |
| Anti-Shank1/2/3 antibody, Mouse | Anti-Shank1/2/3 | Santa Cruz Biotechnology | Sc-393963 AC | Monoclonal | 1:100 |
| Anti-Gephyrin antibody, Mouse | Anti-Gephyrin | Synaptic Systems | 147 111 | Monoclonal | 1:300 |
| Anti-Gephyrin antibody, Mouse | Anti-Gephyrin | Santa Cruz Biotechnology | sc-25311 | Monoclonal | 1:200 |
| Anti-glial fibrillary acidic protein antibody, Mouse | Anti-GFAP | Synaptic Systems | 173 011 | Monoclonal | 1:300 |
| Anti-glial fibrillary acidic | Anti-GFAP | ThermoFisher/ Invitrogen | PA1-10019 | Polyclonal | 1:400 |

| | | | | | |
|---|---|---|---|---|---|
| protein antibody, Rabbit | | | | | |
| Anti-Vimentin antibody, Guinea pig | Anti-Vimentin | Synaptic Systems | 172 004 | Polyclonal | 1:300 |
| Anti-Acetylated tubulin antibody, Mouse | Anti-a-Tubulin | Sigma-Aldrich/Merck | T7451 | Monoclonal | 1:200 |
| Anti-Adenylate cyclase-3 antibody, Rabbit | Anti-AC3 | NOVUS Biologicals | NBP1-92683 | Polyclonal | 1:200 |
| Anti-Ankyrin G antibody, Guinea pig | Anti-Ankyrin G | Synaptic Systems | 386 005 | Polyclonal | 1:300 |
| Anti-Nucleoporin-98 antibody, Rabbit | Anti-NUP-98 | Cell Signaling | 2598 | Monoclonal | 1:200 |
| Anti-Green fluorescent protein antibody, Rabbit | Anti-GFP | ThermoFisher/ Invitrogen | A-11122 | Polyclonal | 1:300 |
| Anti-Myelin basic protein antibody, Mouse | Anti-MBP | BioLegend | 808403 | Monoclonal | 1:200 |
| Anti-Connexin43 antibody, Rabbit | Anti-CnX-43 | Sigma-Aldrich/Merck | C6219 | Polyclonal | 1:200 |
| Anti-Pmp70 antibody, Rabbit | Anti-Pmp70 | Abcam | ab85550 | Polyclonal | 1:400 |

**Table 3: List of primary antibodies**

| Target | Species | fluorophore | Vendor | Identifier | Working dilution |
|---|---|---|---|---|---|
| Mouse IgG (H+L) | Goat | Alexa Fluor 488 | Thermo Fisher/ Invitrogen | A-11001 | 1:400 |
| Mouse IgG (H+L) | Goat | Alexa Fluor 546 | Thermo Fisher/ Invitrogen | A-11030 | 1:400 |
| Mouse IgG (H+L) | Goat | STAR RED | Abberior | STRED-1001 | 1:200 |
| Rabbit IgG (H+L) | Goat | Alexa Fluor Plus 488 | Thermo Fisher/ Invitrogen | A-32731 | 1:400 |
| Rabbit IgG (H+L) | Goat | Alexa Fluor 546 | Thermo Fisher/ Invitrogen | A-11035 | 1:400 |
| Rabbit IgG | Goat | STAR RED | Abberior | STRED-1002 | 1:200 |
| Guinea pig IgG | Goat | STAR RED | Abberior | STRED-1006 | 1:200 |

**Table 4: List of secondary antibodies**

| Solution | Abbr. | Composition | Concentration |
| --- | --- | --- | --- |
| 1X Phosphate Buffered Saline, pH 7.4 | 1X PBS | Sodium Phosphate dibasic ($Na_2HPO_4$) | 10.00 mM |
| | | Potassium Phosphate Monobasic ($KH_2PO_4$) | 1.98 mM |
| | | Potassium chloride (KCl) | 2.68 mM |
| | | Sodium chloride (NaCl) | 136.89 mM |
| Double deionized water, pH 7.4 | $ddH_2O$ | Desalted water | --- |
| Sodium bicarbonate, pH 8.5 | 100 mM $NaHCO_3$ | Sodium bicarbonate ($NaHCO_3$) | 100 mM |
| | | Desalted water | --- |
| Denaturation buffer, pH 9.0 | DB | Sodium dodecyl sulfate (SDS) | 200 mM |
| | | Sodium chloride (NaCl) | 200 mM |
| | | TRIS-HCl | 50 mM |
| | | Desalted water | --- |
| Perfusion solution, pH 7.4 | PS | 4% Paraformaldehyde (PFA) | 4 g/ 100 g |
| | | 10% Acrylamide (AA) | 10 g/ 100g |
| | | 1X PBS | --- |

**Table 5: List of solutions**

Structures of chemicals

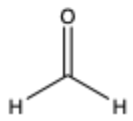

Formaldehyde (FA) Molecular weight (MW) 30.03 (g/mol)

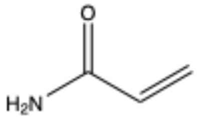
Acrylamide (AA), MW 71.08 (g/mol)

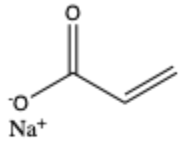
Sodium acrylate (SA), MW 94.04 (g/mol)

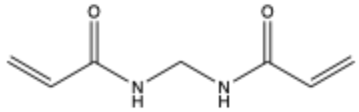
N,N‘-Methylenebisacrylamide (BIS), MW 154.17 (g/mol)

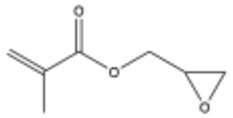
Glycidyl Methacrylate (GMA), MW 142.15 (g/mol)

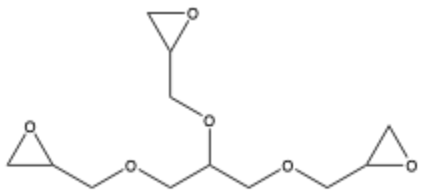
Glycerol triglycidyl ether (TGE), MW 260.29 (g/mol)

<u>Epoxide ring opening under basic condition</u>

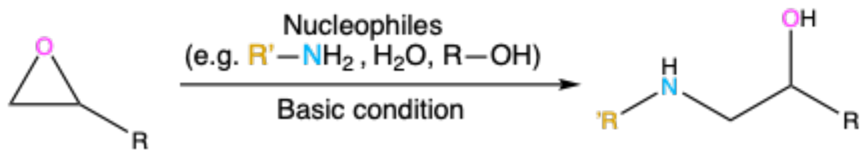


**Experimental procedures**

<u>Transcardial fixative perfusion</u>

Solutions were prepared on the same day and kept at room temperature (RT). Animals were first anesthetized with isoflurane (1-2 % (volume/volume, v/v)) and then with Ketamine (80-100 mg $kg^{-1}$ of body weight) and Xylazine (10 mg $kg^{-1}$) intraperitoneally, combined with Metamizol (200 mg $kg^{-1}$) subcutaneously for analgesia. After checking for deep anaesthesia by toe pinch, they were transcardially

perfused at a flow rate of 7 ml min$^{-1}$ using a peristaltic pump, first with RT 1X PBS for 2 min and then with a solution (RT) containing 4 % (weight/volume, w/v) PFA (pH 7.4) and 10% acrylamide (AA) in 1X PBS for 6 min.
Brains were harvested and postfixed for 8-12 h at 4 °C with gentle agitation, using the same fixative solution.

Brain tissue processing
Brains were washed 3x in cold (4° C) 1X PBS for ~1 min each with gentle agitation. They were sectioned coronally at 50 µm thickness using a Vibratome (Leica VT 1200 S). Where indicated, sections were prepared at 300 µm thickness. Sections were placed in ice-cold 1X PBS supplemented with 100 mM glycine to quench PFA reactive groups for 6-8 h at 4° C and washed 3x for ~1 min in 4 °C 1X PBS. They were stored at 4 °C in 1X PBS supplemented with 0.015% $NaN_3$ for typically up to 3 months.

Epoxide treatment
Brain sections were washed 2x for 15 min each in 1X PBS at RT with gentle agitation. They were then washed 2x for 15 min each in 100 mM sodium bicarbonate in dd$H_2O$, pH 8.0, at RT followed by incubation with 0.1% (w/v) TGE and 0.1% (w/v) GMA in 100 mM sodium bicarbonate in dd$H_2O$ (pH 8.0) for 3 h at 37 °C with gentle agitation in a chemical hood. For 300-µm thick slices, TGE and GMA concentration was increased to 1% (w/v). Samples were then washed with 1X PBS for 1 h with gentle agitation at RT.

Pre-expansion immunolabeling for distortion analysis
Pre-expansion immunolabeling was only performed for distortion analysis measurements. All other immunolabelings were performed post-expansion. Brain slices were permeabilized for 60-80 min with 0.2% Tween-20 in 1X PBS at RT with gentle agitation. Labeling with primary anti-GFP antibody was performed in 5% BSA and 0.2% Tween-20 in 1X PBS ON at 4 °C with gentle agitation. Samples were then washed 3-5x for ~30 min each in 1X PBS at RT with gentle agitation, followed by staining with secondary antibody ON at 4 °C, in the same buffer as for the primary antibody labeling, with gentle agitation, followed by 3-5 washes for ~30 min each in 1X PBS with gentle agitation. Pre-expansion images were acquired on a spinning disc confocal microscope after the first gelation step.

Hydrogel embedding and expansion

| | Acrylamide (% w/v) | Sodium acrylate (% w/v) | N,N′-Methylenebisacrylamide (%w/v) | APS (% w/v) | TEMED (% w/v) | TEMPO |
|---|---|---|---|---|---|---|
| 1st expandable hydrogel | 10 | 12.5 | 0.075 | 0.15 | 0.15 | 0.001 |
| Stabilizing hydrogel | 10 | ---- | 0.025 | 0.05 | 0.05 | ---- |
| 2nd expandable hydrogel | 10 | 19 | 0.025 | 0.05 | 0.05 | ---- |

**Table 6: Composition of hydrogels.**

For preparation of the 1st hydrogel solution, AA and sodium acrylate (SA) were mixed in $ddH_2O$, vortexed and centrifuged at 4500 G for 5 min. Supernatant was transferred to a fresh 50 ml tube and supplemented with N,N′-Methylenebisacrylamide (BIS) stock solution (prepared in $ddH_2O$) to a final BIS concentration of 0.075% w/v. After adjustment to the final volume, solutions were aliquoted in 2 ml tubes and stored at -20 °C, typically for up to 1 month. Solutions for stabilizing and 2nd expandable hydrogels were prepared similarly, usually freshly before experiments.

Polymerization of the first expandable hydrogel

In an ice-water bath, the 1st hydrogel monomer solution (see **Table 6**) was first supplemented with 0.001% TEMPO and then with 0.15% APS and 0.15% TEMED and vortexed. The following sample handling steps were performed in an ice-water bath. Brain sections were pre-incubated with the hydrogel monomer solution for 30-45 min (75-90 min for 300 μm thick slices) with gentle agitation in a 24 well plate. A gelation chamber was assembled from two cover glasses on bottom and top and #1 cover glass strips on either side as spacers (for 300 μm sections: one #1 and one #1.5 cover glass strip). Gelation was then carried out at 37 °C in a humidified chamber for 2h. For optionally acquiring pre-expansion images, the sample was imaged on a spinning disc confocal microscope (Oxford Instruments, Andor Dragonfly) after the first hour of gelation for ~15 min and then further incubated at 37 °C for another ~45 minutes.

Disruption of mechanical cohesiveness, expansion, hydrogel neutralization

After gelation, the top coverslip was removed and the hydrogel-tissue hybrid trimmed to the region of interest. Together with the bottom coverslip, it was then transferred to a 5 ml beaker with 2 ml of denaturation buffer (200 mM SDS, 200 mM NaCl, 50 mM TRIS-HCl in $ddH_2O$, pH adjusted to 9.0 with NaOH). The beaker was placed in a larger vessel containing pre-heated water and transferred to a water bath at 95 °C for 100 min. For 300 μm thick samples, denaturation time was extended to 4 h, starting at 85 °C and ramping up to 95 °C within ~15 min.

For expansion, samples were placed in $ddH_2O$, with water being changed every ~20 min until no further increase in gel size was observed.

We neutralized unreacted double bonds of the divinyl-cross linker (BIS) remaining after polymerization of the 1st hydrogel to ensure that it would not react with the following stabilizing and 2nd expandable hydrogels. For this, we incubated with 0.2% APS plus 0.2% TEMED in $ddH_2O$ for 2.5 h at 37 °C with gentle agitation, followed by one washing step with agitation in $ddH_2O$ for 30 min at 37 °C and a similar washing step at RT.

Immunolabeling

Expanded hydrogels were again trimmed to the region of interest and incubated in 1X PBS for 10 min at RT, shrinking the gel in a 12-well plate. It was then incubated with primary antibodies in 1X PBS at 4°C ON with gentle agitation in a 24-well plate, followed by ≥3 washes for 1 h each in 1X PBS with gentle agitation at RT. Secondary antibody incubation was also performed in 1X PBS at 4°C ON with gentle agitation, followed by ≥3 washes for 1 h each in 1X PBS with gentle agitation at RT. Post-fixation was conducted for 15 min at RT with 4% PFA in 1X PBS, followed by quenching with 100 mM glycine in 1X PBS for 10 min at RT with gentle agitation and subsequent 3x washing in 1X PBS for ~3 min each. Hydrogels were re-expanded in $ddH_2O$ in a 12-well plate before further processing and labeling was optionally evaluated by imaging.

Polymerization of stabilizing hydrogel and neutralization

Expanded hydrogels were pre-incubated on ice-water for 3-3.5 h or 5 h for 50 µm and 300 µm thick slices, respectively, with the monomer solution of the stabilizing hydrogel (10% (w/v) AA, 0.025% (w/v) BIS, 0.05% (v/v) TEMED and 0.05% (w/v) APS in $ddH_2O$) with gentle agitation in a 12-well plate. After removing excess monomer solution, the hydrogel was sandwiched between a 22 x 22 $mm^2$ coverslip placed on a microscopy slide and an 18 mm round coverslip on top of the hydrogel without spacers, and surrounded by monomer solution. Gelation was then carried out for 2 h at 37 °C in a pre-warmed (37 °C) humidified chamber. Gels were washed with $ddH_2O$ for ~30-60 min at RT.

To neutralize remaining unreacted double bonds of the BIS crosslinker in the stabilizing hydrogel network and ensure that the hydrogel network did not react with the following 2$^{nd}$ expandable hydrogel network, the hydrogel-tissue hybrid was incubated with 0.2% APS and 0.2% TEMED in $ddH_2O$ for 2.5 h at 37 °C with gentle agitation, followed by washing with agitation in $ddH_2O$ for 30 min at 37 °C and a second washing step at RT.

Polymerization of second expandable hydrogel

Stabilized and neutralized hydrogels were pre-incubated on ice-water with the monomer solution for the second expandable hydrogel (19% SA, 10% AA, 0.025% (w/v) BIS, 0.05% TEMED, 0.05% APS in $ddH_2O$) in a 12-well plate for 3-3.5 h (50 µm slices), or 5 h (300 µm slices). After removing excess monomer solution, the hydrogel was again sandwiched between a 22 x 22 $mm^2$ coverslip placed on a microscopy slide and an 18 x 18 $mm^2$ coverslip on top of the hydrogel without spacers, and surrounded by monomer solution. Gelation was then carried out for 2 h at 37 °C in a pre-warmed (37 °C) humidified chamber. Hydrogels were washed with 1X PBS for ~30 min at RT.

Protein labeling

Pan-protein staining was performed with either 40 µM ATTO 488 NHS-ester or 40 µM Alexa Fluor 488 NHS-ester in 1X PBS ON at 4 °C with gentle agitation. Hydrogels were optionally washed 3x for 1 h total with 1X PBS, and expanded for up to 4 h before imaging in $ddH_2O$ with fluid exchange approximately every hour.

Mounting of expanded hydrogels for imaging

Before mounting for imaging, the region of interest was located with the spinning disc confocal microscope in the expanded hydrogel after placing it onto a plastic culture dish with the bottom replaced with a 50 mm #1 cover glass. For final imaging, the hydrogel was trimmed to the region of interest. 40 mm round coverslips (Bioptechs, Butler, PA, USA) were rinsed with $ddH_2O$, covered with Poly-L-lysine and incubated at 37 °C for up to 3 h. Coated coverslips were stored at 4 °C in 1X PBS for up to 1 week. Right before mounting, coverslips were washed with $ddH_2O$ and placed in a home-built imaging chamber made from aluminium. The hydrogel was placed in the centre of the coverslip and further stabilized with two-component dental silicone (twinsil extrahart, picodent), and immersed in $ddH_2O$. We typically imaged for up to 8 h.

Imaging: Spinning disc confocal microscopy

Imaging was performed on an Andor Dragonfly microscope based on a Nikon Ti2E inverted stand with motorized stage and an Andor Zyla 4.2 Megapixel sCMOS camera (2048 x 2048 pixels). Data was acquired using Andor Fusion software version 2.2. Two pinhole disc patterns (25 µm and 40 µm hole diameter) and four continuous-wave excitation lasers (405 nm, 488 nm, 561 nm and

637 nm) were available. Imaging of non-expanded and expanded samples was performed with a 40x water immersion objective (Nikon Apochromat LWD 40x lambda S/NA 1.15/Water/working distance (WD) 0.6 mm), using the 40 µm disc pattern. For overview imaging, also a Nikon CFI P-Apochromat 20x/NA 0.95/WD 0.95 mm objective lens was used. The structural channel was imaged with 488 nm excitation wavelength with a 521/38 nm detection bandpass filter. Typical exposure times were 110-150 ms and laser power was set to 10-15% of available laser power. Voxels measured 150 x 150 nm$^2$ laterally and axial ($z$-direction) step size was chosen 200 nm, 300 nm or 400 nm. The lateral field of view (FOV) was typically 307 x 307 µm$^2$, corresponding to ~20 x 20 µm$^2$ biological tissue scale. Axial imaging extent was typically chosen 20-25 µm in biological tissue scale, limited by the 600 µm working distance of the objective lens. For tiled measurements, the "field" mode was used with 10% lateral overlap between individual subvolumes. For additional imaging of immunolabeling channels, color channels (with excitation wavelengths of 488 nm (structural channel), 637 nm, 561 nm) were recorded sequentially in a frame-wise manner with exposure times of 120-200 ms per immunolabeling channel and laser power settings of 10-20%. For detection, 594/43 nm and 685/47 nm bandpass filters were used for the respective color channels. Images for distortion analysis were acquired with 150 x 150 x 200 nm$^3$ voxel size with the 40x/NA1.15 water immersion objective lens.

Extension of imaging volumes in *z*-direction by block-face imaging and sectioning of expanded hydrogels

To seamlessly stitch imaging volumes not only laterally but also in axial direction, we performed the following "lossless" sectioning procedure, ensuring overlap of imaging volumes both laterally and axially. We expanded 300 µm thick brain tissue sections and mounted them on PLL-coated coverslips for imaging as described above. We first imaged a multi-tile volume arranged on a 2D grid. We then removed the hydrogel and twinseal from the imaging chamber and glued the back surface of the hydrogel onto the sample holder of a vibratome (Leica VT1200S, vibrating blade microtome) with superglue (Loctite 401, Henkel), gently flattening out the hydrogel by passing a soft plastic sheet over it. The vibratome chamber was partially filled with water, surrounding but not covering the hydrogel. We zeroed the vertical blade position on the hydrogel surface and lowered it to the desired cutting position chosen to fall within the already imaged region (e.g. cutting 320 µm below hydrogel surface after imaging an axial range of 466 µm physical imaging range in fig. S31). Cutting was performed at 0.2 mm/s blade advancement. We then removed the hydrogel from the vibratome with the plastic sheet, mounted it for imaging with the cut surface facing the PLL-coated coverslip and imaged a second layer of tiled, overlapping imaging volumes similar as before.

### Data analysis

Data handling and format conversion

Raw imaging data was obtained in ims format, as generated by the spinning disc confocal microscope (Andor Dragonfly), which is a hierarchical data format with 6 resolution levels, from which we extracted the highest resolution level (level 0). Data conversion for downstream analysis was performed with custom python scripts implemented in Python v3.8 or higher, including the Imaris-ims-file-reader, zarr, webKnossos and tifffile packages. At various processing steps, file format conversion was necessary and was performed on ISTA's high-performance computing cluster, using either a single node or SLURM for task allocation, according to the resource requirement based on file size. We transferred files to cluster storage and converted from the respective source format to tiff, zarr, n5 or webKnossos (wk) format according to processing needs.

Image processing

Imaging data were processed using Fiji v.1.54f, including the CLAHE, BigWarp and BigData viewer plugins. For visualization, display ranges were adjusted, also accounting for a camera background signal of ~100. Intensity lookup tables were linear unless otherwise noted.

Overlay of immunolabelings with LICONN data was performed in Gimp version 2.10.34 after saving individual channels separately in RGB format in Fiji. Background in immunolabelings was set to transparent (alpha = black) and immunolabelings were overlaid with the structural channel. The output was saved in png format.

CLAHE (contrast limited adaptive histogram equalization) was applied as indicated for individual figure panels, using either Fiji or a Python based workflow as described below. In Fiji, first the display range was adjusted and in the CLAHE plugin the block size and histogram bin were set to the minimum and maximum values of the display range. The maximum slope was set to 3 and the standard (not fast) processing option was chosen. 3D-renderings of cilia were done with Imaris version 9.3.

Distortion analysis and calculation of expansion factor

Analysis of distortions in the expansion procedure was adapted from previously published methods (*1*, *2*) and code was adapted from (https://github.com/Yujie-S/Click-ExM_data_process_and_example). We analyzed a total of 14 imaging volumes of ~20 x 20 µm$^2$ laterally and 9-15 µm axially, recorded across 4 technical replicates in *n*=3 *Thy1-eGFP* mice, featuring cytosolic expression of eGFP in a sparse subset of neurons. Imaging volumes corresponded to single tile measurements (no stitching/fusion of subvolumes). For distortion analysis, immunolabeling against eGFP was performed before subjecting the samples to the LICONN expansion procedure and the eGFP channel was used for analysis. Pre-expansion images were acquired after ~1 h of polymerization of the first hydrogel. Imaging volumes containing the same structures were acquired after expansion of the first hydrogel (before application of the stabilizing hydrogel) and after expansion of the second expandable hydrogel, with a spinning-disc confocal microscope and a 40x/NA 1.15 water immersion objective lens, as in LICONN imaging. This allowed assessing distortions beyond the ~200 nm spatial scale. Using the BigWarp (*3*) plugin in Fiji, we first manually placed ~25 landmarks at corresponding neuronal features in maximum intensity projections of the respective pre- and post-expansion imaging volumes. Using BigWarp, we then applied either a similarity transformation (including isotropic scaling, translation and rotation) or an affine transformation (including scaling, translation, rotation, and shearing) to the pre-expansion images to best match landmarks in pre- and post-expansion images. Expansion factors were extracted as the linear scale factor in the similarity transformations. We smoothed pre- and post-expansion projection images with Gaussian filters of different widths to account for the different resolution in the respective images. For individual expansion steps, σ was set to 1 and 6 pixels for pre- and post-expansion images, respectively. For total (iterative) expansion, σ was set to 1 and 16 for pre- and post-expansion images, respectively, resulting in comparable appearance as judged by visual inspection. For one of the datasets, the intensity range was slightly saturated at the main dendrite branch in the pre-expansion image. We now used the imregdemons.m function in MATLAB (version R2022b, MathWorks) to compute distortion vector fields (*1*, *4*) from the aligned pre-/post-expansion images. For this, we used the images before the respective expansion step as a reference and evaluated distortions arising from that expansion step. For display purposes, we used the quiver function in Matlab with vector scaling set to 1.5. We then computed and plotted distortions (measurement error) for different measurement lengths, as previously described (*5*). For this, we generated binary masks of neuronal structures of interest by thresholding pre- and post-expansion images, and then combining, dilating and eroding the resulting masks using parameters as judged by visual inspection. We then randomly sampled $2*10^5$ pairs of points restricted to the masked area of interest. These pairs of points defined a set of vectors in the pre-expansion image. We then found a corresponding set of transformed vectors by applying the distortion field generated with imregdemons.m to the respective pairs of points. For each measurement length, we then computed the measurement errors as lengths of the difference vectors between the pre-expansion and transformed vectors combined from 3-4 different datasets within one experiment. From these measurement errors, the root mean square (RMS) error was calculated across different measurement scales. Finally, the mean and standard deviation of RMS errors across the *n*=4 technical replicates were plotted for individual expansion cycles and transformations. For display purposes, we smoothed the measurement error curve with a 1D median filter.

Stitching and fusion of imaging tiles
We extended SOFIMA (https://github.com/google-research/sofima) to support seamless stitching of 3D tiles. Briefly, we processed every volumetric tile with CLAHE, applied independently to every in-plane (*xy*) section of the tile. The tiles had an in-plane size of 2048x2048 $voxel^2$ and axial extent varying depending on experiment. We laid out the tiles on a 2D grid according to imaging metadata, and used a coarse positioning step to find their initial location in the coordinate system of the output volume. This was followed by fine alignment to elastically deform them to obtain seamless matches between adjacent tiles and form 3D sections, ~20-30 um thick. The tiles were then rendered with linear distance-weighted blending of overlapping image content. For the coarse positioning step, we extracted a small $[50\ voxel]^3$ cube of image content from the 'preceding' tile of every adjacent tile pair, and used cross-correlation to identify the corresponding image content in the 'following' tile, forming an offset vector between the tiles. The tiles were then modelled as unit masses connected by springs of resting lengths equal to the computed offsets, and the system was allowed to relax, with the relaxed state determining the tile positions at the beginning of the fine alignment step. For fine alignment, we modelled every tile as an elastic mesh of unit masses placed on a regular grid with a 3D-stride of 20 voxels and nearest neighbor masses connected with Hookean springs. Coarse positioning resulted in the neighboring tiles partially overlapping each other, and for any mass located in the overlapping image region, we again used cross-correlation to identify the position within the neighboring tile of a $[80\ voxel]^3$ image patch centred on the mass (PATCH_SIZE=80). These formed point correspondences between the tiles, which we modelled as 0-length Hookean springs, and allowed the whole system to relax. The relaxed set of tile meshes was used to render the aligned volume.
For datasets comprising more than one thick section (fig. S31), we first stitched tiles into thick sections as described above, and then aligned them to each other. To do so, we manually identified the amount of overlap in the axial dimension (typically 100-200 voxels), used the overlapping subvolume to compute a 3D flow field (with a stride of 40 voxels and a patch size of 80 voxels), modelled the following section as a 3D spring mesh, and allowed it to relax according to the flow field. The relaxed mesh was then used to render the following section in alignment with the preceding one. This is similar to the "fine alignment" of individual tiles, and the overall strategy is a direct 3D analogue of how alignment of 2D sections is done with SOFIMA for volume electron microscopy datasets.

Details of processing were performed as follows. In all cases, we based stitching and fusion on the structural imaging channel. We first transferred individual raw ims files (uint16 format) to cluster storage and extracted the highest resolution level (level0). We then used a BigStitcher-based FIJI macro to concatenate the data and convert to n5 format. We then applied CLAHE tile-by-tile with Python. Data handling was done with the tensorstore library (google.github.io/tensorstore, version 0.1.33). We used the numpy.clip function to clip the intensity range plane-by-plane typically between 120 and 350, determined by visual inspection. For CLAHE, we used skimage.exposure.equalize_adapthist to apply histogram equalization, with a clip limit of 0.03. We multiplied the output (ranging from 0 to 1) with 255 and saved as uint8 format.

In the rigid, coarse alignment step, we determined the relative arrangement of tile pairs inputting the known tile layout (arrangement of tiles in the image acquisition). We set the QUERY_R_ORTHO and QUERY_R_OVERLAP parameters to 25. We set the minimum tile overlap (QUERY_OVERLAP_OFFSET) to 60 (100 for dataset in Fig. 2A) and the size of the search area within tiles (SEARCH_OVERLAP, SEARCH_R_ORTHO) to 300 (400 and 600, respectively, for dataset in Fig. 2A). The second, fine stitching, step was used to warp tiles and achieve a smooth transition at tile borders, using a stride of (20, 20, 20) voxels. This step outputs a mesh that ensures a globally consistent warping and relative arrangement of individual tiles. In the last step, we applied the obtained mesh to either the CLAHE-modified structural imaging data for downstream (automated) segmentation/skeletonization, or the raw structural imaging data, as well as to immunolabeling channels, resulting in the respective fused imaging volumes in n5 format. We performed these processing steps on ISTA's high performance computing cluster or, in the case of the measurement in Fig. 2A and fig. S31, on Google's high-performance computing infrastructure, requesting resources according to the processing task. Typically, for CLAHE, we requested 16 CPUs (Intel Xeon CPU E5–2680 v4 @ 2.40-GHz or similar) and 150 GB of RAM. For coarse stitching, we typically requested 3 GPUs (GeForce RTX 2080 Ti or similar), 32 CPUs and 100 GB or RAM. Fine stitching was done with 1 GPU, 16 CPUs and up to 970 GB of RAM. Rendering was typically done requesting 32 CPUs and up to 700 GB of RAM.

Manual segmentation and proofreading

For manual segmentation, we used VAST version 1.4.0 (downloaded from https://lichtman.rc.fas.harvard.edu/vast/) with the data in 8-bit format. Segmentation of the data in Fig. 1G was originally performed by one person, and the resulting segmentation was used for training. For visualization in Fig. 1G, segmentation was partially proofread by another person using VAST v1.4.1.

Skeletonization

We used webKnossos v.22.05.1 for manual skeletonization, running on a local computer with 2× AMD EPIC MILAN 75F3 processor, 32-core 32C/64T, 2.95 GHz with 512 GB of memory, 2 TB NVMe-SSD and an additional 27 TB NVMe-SSD, as well as 4 NVIDIA A6000 GPUs. To reduce required memory resources, we converted data to uint8 format when converting to the webKnossos (wk) format. Before conversion, we clipped the intensity range according to visual inspection, to account for intensity outliers and avoid signal overflow. For convenient exploration of the datasets across spatial scales, we made use of 10 precomputed resolution levels, using the webKnossos downsample function. Typically, raw data rather than CLAHE-processed data was used for tracing. When comparing automated segmentation of the volume in Fig. 2A with manually generated skeletons, we checked sites of discrepancy and corrected skeletons as appropriate.

Generation of GFP-based ground truth

For testing traceability by comparison with an independently generated ground truth, we compared manual tracings of neurites in the structural LICONN channel with their structure revealed by cytosolic expression of eGFP in *Thy1-EGFP* mice. The LICONN structural channel and the eGFP channel were recorded at similar spatial resolution after tissue expansion with a spinning disc confocal microscope. The structural channel was recorded in the 488 nm excitation channel, whereas eGFP was detected by post-expansion immunolabeling as described above with imaging at 561 nm excitation. Voxel sizes were either 150 x 150 x 200 $nm^3$ or 150 x 150 x 300 $nm^3$, with the largest step size referring to the *z*-direction. Data were converted to wk format for tracing. Using the webKnossos access permissions feature, we assigned permissions to ensure blinding as required. For generating "ground truth" skeletons of eGFP expressing neurons, both the structural and the eGFP channels were loaded and jointly considered. In addition, one seed point was placed in each eGFP-expressing neuronal structure. Together with the LICONN channel, these seed points were provided to two human annotators blinded to the eGFP channel who independently traced the marked structures based on the structural channel.

For dendritic spines, annotators were asked to indicate these as short branches diverging from the main dendrite trunk. The annotators were then asked to compare the individual skeletons and generate a "consensus" skeleton for each of the traced structures. Human consensus skeletons were then overlaid with and compared to the eGFP-based ground truth using webKnossos. Before analysing the test datasets, tracers were allowed to train on 2 eGFP-expressing single-tile training datasets for dendrites and 2 single-tile datasets for axons in a non-blinded fashion. These datasets were not included in further analysis.

Neuronal instance segmentation

We used flood-filling networks (FFNs, available at https://github.com/google/ffn) (*6*) to automatically segment the datasets. We used the original convolutional stack architecture with a 33 x 33 x 33 voxel field of view (FOV), step size of 8 voxels, and network depth extended to 20 residual modules. The models were applied to data with 19.4 x 19.4 x 25.9 $nm^3$ voxel size (in original tissue scale). For this, CLAHE-processed volumetric imaging data was 2x downsampled laterally relative to the imaging voxel size using volume averaging in Python. Axially, datasets with 200 nm axial imaging step size (effective voxel size 9.7 x 9.8 x 13.0 $nm^3$, obtained by scaling with exF of 15.44, e.g. Fig. 2A) were also downsampled 2-fold in *z*-direction, whereas datasets with 400 nm axial step size (effective voxel size 9.7 x 9.8 x 25.9 $nm^3$, e.g. Fig. 1A) were not downsampled in *z*-direction. Training was performed with a batch size of 128 and learning rate of $10^{-4}$ using the AdamW (*7*) optimizer with 32 NVIDIA V100 GPUs, for up to 30 days, which corresponded to about 8.8 Million training steps.

We used an iterative approach to collect the volumetric ground truth annotations for training the FFNs. First, 162 neurite fragments covering 110 Mvx (Megavoxels, at full imaging resolution) were manually annotated as described above, using VAST in the dataset displayed in Fig. 1G (single tile). The model #1 trained with this data on full resolution imaging data was used to segment the multi-tile volume in Fig. 2A. We then manually agglomerated 103 neurite fragments covering 447 Mvx (voxel number referring to full imaging resolution) within that segmentation (for description of manual agglomeration see below). We used this to train another FFN model (#2), using both the newly collected data and the preexisting manually painted annotations for training with voxels 2x downsampled in every direction (19.4 x 19.4 x 25.9 $nm^3$ effective voxel size). Within the segmentation results of model 2, we manually proofread a larger set of axons (540 axons, 920 Mvx at full imaging resolution,115 Mvx downsampled, 17.8 mm cumulative path length) and dendrites (314 dendrites, 3,344 Mvx at full imaging resolution, 418 Mvx downsampled, 23.8 mm cumulative path length), and used this data to train the final (#3) FFN model which was used to produce all the automated neurite segmentations displayed in the manuscript.

We followed a segmentation strategy optimized for first reducing the number of segments affected by merge errors ("base segmentation"), and then agglomerating these largely merge-free supervoxels into larger neurites, as described in the following sections. For the base segmentation, we applied the seed-order oversegmentation consensus (OC) procedure (*6*), followed by OC with two more segmentations generated with alternative FFN checkpoints (snapshots of weights saved at regular time intervals during training), which were manually selected for having a small number of wrong merges. This reduced the number of falsely merged supervoxels in the segments evaluated with manually generated skeletons in the volume in Fig. 2A to 0 whereas the overall volume contained at least 5 mergers.

We automatically skeletonized the base segmentation for downstream use in agglomeration and proofreading using the TEASAR algorithm (*8*) and heuristically identified all nodes of degree 1 in the resulting skeleton graphs as "endpoints".

Semantic segmentation

To automatically classify segments into distinct subclasses, in particular axons and dendrites, we developed a semantic segmentation pipeline.

For this, an experimenter classified manually agglomerated objects in a segmentation of a 512 x 512 x 334 voxel subvolume (at the downsampled 19.4 x 19.4 x 25.9 $nm^3$ effective voxel size) of the dataset in Fig. 2A into one of five classes: axons (331 segments, 15 Mvx), dendrites (36 segments, 11.1 Mvx), myelinated axons (6 segments, 1.1 Mvx), oligodendrocytes (2 segments, 0.15 Mvx), and (astro)glia (47 fragments, 1.3 Mvx). We used these annotations to train a neural network model to perform semantic segmentation (i.e. voxel-wise multi-class prediction) of the structural LICONN data. The neural network used the same residual 3D convolution stack architecture as the FFN, with a depth of 8 residual modules and a FOV of 33 x 33 x 33 voxels, but with the convolutions operating in 'valid' mode, and the residual summation discarding context at the boundary of the larger-sized argument. The model was trained with asynchronous stochastic gradient descent (SGD) at a learning rate of $10^{-4}$, using 16 NVIDIA V100 GPUs and a batch size of 16. Training lasted approximately 38h, during which 40.7 Million steps were executed. Training points from all five classes were sampled at equal frequencies.

We used the trained model to generate a semantic segmentation of the entire volume in Fig. 2A and associate every voxel with the class predicted with the highest probability. These voxelwise classifications were then aggregated over all voxels of every instance segment, and the class predicted most frequently was associated with that segment.

Segment agglomeration

The instance segmentation pipeline described above is optimized to make it unlikely that an individual super-voxel incorrectly covers more than one neurite. This happens at the cost of an increased number of split errors, i.e. assigning multiple smaller segments to the same neurite.

To mitigate this, we applied a multistep agglomeration procedure in which the FFN model was used to evaluate whether a pair of geometrically proximal segments should be merged together (*6*) and how a segment could be further extended from a heuristically detected endpoint in the skeletons generated from the base segmentation (*9*). In the first step, we used the "relaxed acceptance criterion" (*9*). In the second step, we merged two segments when the endpoint-seeded FFN inference recovered 90% of the voxels of both segments within a $[200\ \text{voxel}]^3$ subvolume centred at the endpoint.

We further trained two FFN models with a 3D U-net architecture and a $128^3$ voxel FOV and without a FOV movement policy. We used the same training data as for the main instance segmentation model, but split it into two non-overlapping sets, so that one model was trained exclusively on axons, and the other exclusively on dendrites. We applied the axon model to endpoints of all axon segments. Within the prediction results for every endpoint we determined the segment *S* with the highest recovery fraction (other than the segment *E* containing the endpoint) and agglomerated *S* and *E* if that fraction was at least 50%, and if *S* had an endpoint within 100 voxels of the endpoint of *E.* We applied the dendrite model to all skeleton nodes of dendritic trunks (skeletons generated from the base segmentation), taking any dendritic segment of at least 100,000 voxels to be a trunk. For every evaluated point, we collected any putative spine (dendritic segment of less than 100,000 voxels) associated with at least a 50% recovery fraction. We then agglomerated all spines to the trunk for which it had the highest recovery fraction.
The agglomeration procedure resulted in a graph with 516,437 edges, where each edge represents one pair of automatically joined base segments.

Agglomeration graph filtering

The semantic segmentation model allowed us to associate per-class voxel counts with all base and agglomerated segments. We used these classifications to filter the agglomeration graph similarly to prior work (*9*). Specifically, we sorted the edge list in decreasing order of the associated scores, and discarded any edge that would cause a merge between inconsistent classes. The class combinations we considered were: (astro)glia vs. axons vs. dendrites vs. oligodendrocytes; (astro)glia and oligodendrocytes vs. others; (astro)glia and oligodendrocytes vs. axons vs. dendrites. An (agglomerated) segment was considered to belong to one of these composite classes when it had a total size of at least 100,000 voxels, and at least 75% of its voxels were classified as the target class. This criterion was relaxed to 10,000 voxels and 50%, respectively, for the endpoint agglomeration step, in which we also disallowed merging together objects larger than 100,000 voxels.

Manual proofreading of neuron segmentations

The automated segmentation of the dataset obtained as a result of the agglomeration procedure was expected to have remaining split and merge errors, which we decided to correct in a structured manual proofreading workflow, leading to a marked increase in proofreading speed. The workflow was implemented as a Python script driving the Neuroglancer viewer, within which an expert annotator interactively inspected segments and the structural channel images in both 2D cross-sections and as 3D mesh rendering. The tool had features for automatically moving the 3D-cursor to predetermined locations (such as end points of segments), marking segments for future view, and directly editing the agglomeration graph by adding or removing edges.

Using the results of the semantic segmentation to classify all agglomerated segments, we organized the proofreading process as follows:

1. All axon segments (n=2,674) of at least 10,000 voxels and touching a single face of the bounding box of the volume were inspected for splits and manually connected to appropriate partner segments where necessary. The tool was configured to automatically navigate to heuristically detected endpoints of the axon segments.
2. All axons within the volume (n=18,667) were reviewed for shape implausibilities resulting from putative merge errors, which were then inspected and corrected.
3. Dendritic spines not connected to a trunk (n=2,782) were reviewed and connected to a trunk when one could be unambiguously identified.
4. All dendritic branches (n=1,643) were reviewed for incorrectly associated spines, which were separated, and subsequently manually connected to the correct trunk.

Synapse detection via immunolabeling

To automatically detect pre- and postsynaptic sites in volumetric datasets comprising the LICONN structural channel and immunolabeling channels for Bassoon and Shank2, respectively, we loaded the multi-color image stacks in tiff format. We converted image stacks for molecular signals to 32-bit floating numbers and adjusted contrast to a minimum of 101 to account for camera background and the maximum to the 99.5$^{th}$ and 99$^{th}$ percentile for the dataset in Fig. 4L (300 nm *z*-step size) and 99.95$^{th}$ and 99$^{th}$ percentile for the datasets in Fig. 3G,I and 4Q (200 nm or 400 nm *z*-step size) for Bassoon and Shank2, respectively. We then applied background subtraction by applying Gaussian filters of two different widths using scipy.ndimage.gaussian_filter, with σ=5 voxels and σ=11 voxels, corresponding to signal and background, respectively for Bassoon (σ=6 voxels and σ=10 voxels for datasets with 200 nm *z*-step size) and σ=4 voxels and σ=11 voxels (σ=6 voxels and σ=12 voxels for datasets with 200 nm and 400 nm *z*-step size) for Shank2. Background was subtracted from signal and negative values were set to zero. We then applied Otsu thresholding and used the skimage.measure.label function for converting the resulting binary mask into instance segmentations. We then applied the regionprops function to extract the maximum intensity value for the underlying LICONN channel for each of the 3D-regions defined by the individual segments. The list of maximum intensity values had a bimodal distribution, which we used to distinguish between synaptic regions (high intensity in structural channel) and unspecific immunolabeling (low intensity in structural channel). To separate the two populations, we again used Otsu thresholding. To remove further non-specific detections, we performed additional size filtering. Segments that occupied <60 voxels were removed, as well as objects that spanned less than a certain number of planes in the *z*-direction (300 nm *z*-step size: <13; 200 nm *z*-step size: <14 for Bassoon and <12 for Shank2; 400 nm *z*-step size: <12 for Shank2). Listed parameters were determined from a grid search on a separate validation dataset or a subvolume of the analyzed dataset for each of the *z*-step sizes, choosing the parameter set that resulted in the highest F1 score. We finally converted the individual segments into point annotations, using the regionprops function to find the weighted centroid for each segment based on the underlying LICONN data.

For detection of Gephyrin positive post-synapses, we applied a simplified algorithm involving background removal, Otsu-thresholding, morphological opening and size filtering and manually proofread the output.

Validation of immunostaining-based pre- and post-synapse detection

For validation of automated synapse detection, a human annotator created ground-truth synapse annotation in a dataset comprising the LICONN structural data, Bassoon immunolabeling and Shank2 immunolabeling. For each synapse, they generated a "skeleton" with the source point in the pre-synapse and the target point in the post-synapse, near the respective centres of the molecular signals using webKnossos v.22.05.1, after conversion of the datasets to wk format. Bassoon signals without corresponding Shank2 signals, likely in large part corresponding to inhibitory synapses, were annotated as single source points. Similarly, at volume edges, instances of Shank2 signals without corresponding Bassoon signals occurred, which were annotated as single target points. The human annotator also indicated synaptic connections which lacked Shank2 signal but displayed a clear post-synaptic density in the structural channel as excitatory post-synapse. The special case of a single post-synapse connected to more than one pre-synapse was handled by text comments. In total, two datasets were annotated. One test dataset had 300 nm z-step size and comprised 700 x 700 x 200 voxel of 9.7 x 9.7 x 19.4 $nm^3$ size. This dataset comprised 261 pre-synapses, 247 post-synapses, forming 242 full synapses (with one-to-many connections contributing a count of 1 with each edge), with 25 occurrences of unpaired pre-synapses and 7 unpaired post-synapses. The second test dataset had 200 nm z-step size and comprised 1000 x 1000 x 600 voxel of 9.7 x 9.7 x 13.0 $nm^3$ size. This dataset comprised 1116 pre-synapses, 1084 post-synapses, forming 1059 full synapses (with one-to-many connections contributing a count of 1 with each edge), with 57 occurrences of unpaired pre-synapses and 25 unpaired post-synapses. We generated further validation datasets for parameter tuning, which were not included in the test data.

To compare the automatically generated point annotations of Bassoon or Shank2 signals with the corresponding human ground-truth point annotations, we performed spatial matching of the respective point annotations. For this, we defined a cost matrix with entry $C_{ij}$ corresponding to the Euclidean distance between automatically generated point annotation i and manually generated point annotation j. We defined a matching limit $r_{matching}$, i.e. an upper distance beyond which matching was not allowed and set it to $r_{matching}$=30 voxels after stepping this parameter in a test dataset. For $Cij>r_{matching}$, we set this matrix entry to an arbitrary large number. We now used the scipy.optimize.linear_sum_assignment function to solve the linear assignment problem of finding corresponding points, i.e. the single-point to single-point matches with minimum Euclidean distances. Pairs matching between automatically and manually generated annotations were classified as true positive (TP) automated detections. Automated detections without corresponding ground truth element within the matching radius $r_{matching}$ were classified as false positive (FP) detections. Ground truth detections without corresponding match in the automated detection were classified as false negative (FN) detections.

Association to full synapses

To connect corresponding pre-synaptic sites automatically identified based on Bassoon immunolabeling with post-synaptic sites automatically identified based on Shank2 immunolabeling, we first defined a cost matrix with entries $S_{ij}$ corresponding to the Euclidean distances between pre-synaptic annotations j and post-synaptic annotations i. We then defined a binary matching matrix $M_{ij}$ with $M_{ij}$=1 indicating a connection between the respective pre- and postsynaptic sites (else 0).

Solving this as a linear assignment problem as above would result exclusively in 1:1 matches, which would not capture cases where one pre-synapse is connected to multiple post-synapses or vice versa. We therefore developed a pipeline that gave priority to the spatially closest occurrences of Bassoon and Shank2 during the pre-/post-synaptic pairing procedure but additionally allowed for one-to-many connections.

In a first pass, we paired every post-synaptic site with the closest pre-synaptic sites, i.e. we identified the smallest $S_{ij}$ within each row of the cost matrix and set the corresponding $M_{ij}$ to 1 in case $S_{ij}$ was below a maximum matching distance of $d_{matching}$=30 voxels. This value was taken from the distance distribution between manually generated pre- and post-synaptic annotations, choosing a value close to the observed maximum distance of 31 voxels.

Hence, each post-synapse was at most associated with one pre-synapse whereas each pre-synapse could be associated to 0, 1 or more post-synapses. In a second pass, we now exclusively considered those Bassoon signals that were left unpaired in the previous run ("floating" pre-synapses), i.e. Bassoon occurrences j that did not have a $M_{ij}$=1 after the first pass. For these, we found the post-synaptic entry i with the lowest distance and set the corresponding $M_{ij}$ to 1 in case this distance was below the matching limit $d_{matching}$. This identified the previously unpaired pre-synapses that had Shank2 signal in close spatial proximity, i.e. within the matching limit.

First associating the closest pre- and post-synaptic sites in this way did not allow additional connections between pre- and post-synaptic detections that were already assembled into a synaptic connection in a more closely matching pair, which would occur if simply matching all pre- and post-synaptic occurrences within the maximum matching distance $d_{matching}$.

Validation of synapse assembly

We classified automated detections of synaptic connections (i.e. a connection between an individual pre-synaptic annotation and an individual post-synaptic annotation) as true positive if the following criteria were fulfilled: i) true positive detection of a Bassoon occurrence, ii) true positive detection of a Shank2 occurrence and iii) an automatically detected connection between these two that was also present in the manually annotated connections. We classified automated detections of synaptic connections as false positive in case i) a pre-synaptic and a post-synaptic detection were automatically connected but were not connected in the manual annotation or ii) a false-positive molecule detection (i.e. Bassoon or Shank2 prediction) were assigned a synaptic connection. We classified detections of synaptic connections as false negative in case a manually annotated synaptic connection was not present in the automated detection of synaptic connections. A typical scenario for this was a missed automated detection of pre- or post-synaptic sites, which can also happen if two neighboring Bassoon occurrences (or Shank2 occurrences) were blurred into each other during the smoothing step and thus undersegmented.

For the determination of F1 scores, we revised false negative detections that were close to the border of the analyzed imaging volume in *z*-direction, since potential true positive detections near the border may have been removed during the z-size filtering step. For this, we automatically generated 2 lists of detections: one that included objects spanning less than the specified number of planes in the *z*-direction and a second in which these structures were removed. Additional true positive detections were then defined based on the following rules: i) the detection was true positive in the first list and false negative in the second list, ii) the detection was located within the first 10 or last 10 imaging planes of the volume.

Automated post-processing of synapse detections

To account for those cases where a particular synapse failed to be retained during one of the processing steps for automated synapse detection, we implemented an automated postprocessing step. Such cases included failed detection in case the synaptic immunolabeling was either weak or absent at a particular synapse and therefore did not meet the cutoff threshold (e.g. due to lack of Shank2 expression), or the sampling of the structural LICONN feature did not pass the global threshold, or size filtering. Here, we identified unpaired pre- and post-synaptic detections and defined a local search area around those and checked whether bright structural features were present in the LICONN channel around those unpaired detections. In case a prominent post-synaptic density or bright pre-synaptic feature were present, we classified these as additional pre- or post-synaptic detections, and added them to the respective lists of pre- and post-synapses, including them in the pre-/post-synaptic matching algorithm.

In brief, we checked individual rows and columns of the matching matrix $M_{ij}$ that only contained zeros, corresponding to unmatched post- and pre-synapses, respectively. We then defined a 100 x 100 x 100 voxel$^3$ bounding box around those locations (or smaller at imaging volume edges). We then applied intensity rescaling to the structural LICONN channel between p1 and p99.95, where p1 refers to the first percentile and p99.95 to the 99.95th percentile. We then removed background using Gaussian filters of two different widths at σ=2 voxels and σ=10 voxels, subtraction, and clipping of negative values to zero. We then performed thresholding at thr = p95 + 0.4 × (p100 − p95), with p95 and p100 being the 95th and 100th percentile, respectively. We then removed previously detected pre- and post-synapses by multiplying the resulting mask with a mask corresponding to an inverted semantic segmentation of the previously detected pre- and post-synapses. We then did a binary opening (eroding and then dilating) with a ball of 2-voxel radius. We then performed instance segmentation with skimage.measure.label, removed structures smaller than 20 voxels and determined the weighted centroid based on the intensity distribution in the underlying structural LICONN channel. We classified the resulting objects as a new pre- or post-synaptic detection when their Euclidian distance from the originally unpaired pre- or post-synaptic site was smaller than the matching limit $r_{matching}$. We used these to update the list of pre- and post-synaptic detections used for the association to full synapses.

Deep-learning based synapse detection

We devised a strategy for identifying locations of pre-synapses and post-synapses from the structural imaging channel alone. We employed deep learning prediction of synapse location with an approach that did not require human input for the generation of training data. For this, we implemented a U-net architecture for image translation to predict synaptic molecule location from structural data, using measured molecule locations as input to the training. We then converted the predicted molecule locations to automatically generated synapse detections, for this step using a pipeline that was similar to the immunolabeling based pipeline above. We generated two independent models for predicting the location of the presynaptic marker Bassoon and for predicting the excitatory-synapse specific postsynaptic marker Shank2.

*Generation of training data*

We converted 3D super-resolved measurements of synaptic molecule locations (based on specific immunolabeling), paired with the structural imaging channel, to training data with the following workflow. We first converted tiff images to 32-bit floating format and split them into individual datasets for the structural and each of the immunolabeling channels, maintaining the original voxel size. We then implemented a preprocessing pipeline to automatically remove unspecific immunolabeling signals, similar to the approach taken for the immunolabeling based synapse detection described above. However, unlike in that pipeline (where we performed a strong

smoothing step to avoid oversegmentation of the spatially structured Bassoon and Shank2 distributions), we here preserved the spatial distribution of the Bassoon and Shank2 signals more closely, allowing the deep-learning network to recapitulate the molecule arrangement within the respective synaptic compartments, including the lattice-like arrangement of Bassoon, paralleled by high-intensity (protein-rich) structural features.
We rescaled intensity using the skimage.exposure.rescale_intensity function, setting the minimum to 101 and the maximum to the 99.995$^{th}$ percentile. We then removed background by applying isotropic Gaussian filters of two different widths using scipy.ndimage.gaussian_filter, with σ=0.5 voxels and σ=10 voxels, corresponding to signal and background, respectively, subtracting background from signal and clipping negative values to 0. We applied Otsu thresholding to generate a binary mask and used the skimage.measure.label function to derive an instance segmentation. We then sampled the maximum intensity value of the underlying structural data using the skimage.measure.regionprops function. We applied Otsu thresholding to distinguish segments with underlying high intensity LICONN features, which we classified as specific immunolabelings, and without high-intensity features, which we classified as non-specific immunolabelings. We next removed the latter and converted the remaining segments back to an overall binary mask. To approximate the intensity distribution of the immunolabelings, we blurred the resulting mask with a 3D-Gaussian of σ=1 voxel.

For the structural channel, we applied simple intensity normalization to image pixel values $im$ ($im' = \frac{im-min}{max-min}$), where $min$ is the 0$^{th}$ percentile and $max$ is the 100$^{th}$ percentile of the intensity distribution. We converted to 8-bit format by multiplying the normalized data with 255 and clipping the intensity range between 0 and 255, followed by type casting.
We saved the paired immunolabeling and structural data in zarr format, allowing resource-efficient handling of large datasets.

*Network architecture and training*
We implemented deep-learning pipelines with Pytorch v.1.12.1 (https://pytorch.org) and used the Gunpowder framework v 1.2.2 (https://github.com/funkelab/gunpowder) to implement our data loading, augmentation, training, and prediction pipeline that conveniently allowed processing of big datasets. For prediction of molecule location, we used the 3D U-net architecture from reference[54]. We used Adam's optimizer (torch.optim.Adam) with $10^{-4}$ learning rate. We used the torch.nn.MSEloss function to implement mean squared error as loss function. The batch size was set to 8 and the training input size was 64x128x128 voxels, with the smallest number referring to the $z$-direction.

We performed network training on ISTA's high-performance computing cluster, using SLURM for task allocation. We typically requested 8 CPUs, graphics acceleration by 1 GPU (Nvidia GeForce) and 700 GB of RAM.

We trained on datasets with 300 nm $z$-step size (6 individual tiles, 16.699 μm$^3$ total volume, containing ~16.250 excitatory synapses) and applied the resulting model to datasets with 200 nm, 300 nm or 400 nm $z$-step size. We performed training with 100.000 iterations and saved intermediate checkpoints every 5.000 iterations. For selecting a model for the different $z$-step sizes, we applied models from various checkpoints and took the best-performing checkpoint when evaluating F1 on the respective test datasets. For 200 nm $z$-step size we took the 70.000 and 50.000 iteration checkpoints for Bassoon and Shank2 prediction, respectively, and applied the same model

also to 400 nm *z*-step size. For 300 nm *z*-step size datasets, we used the 20.000 iterations checkpoint both for Bassoon and Shank2 prediction.
During the validation phase for the 300 nm *z*-step size prediction, training data comprised 5 of the 6 tiles (total volume 14.207 $\mu m^3$, with ~13.650 excitatory synapses) while one dataset was used for testing. For final training, all 6 datasets were included. During training, patches of training data (normalized between 0 and 1 with gp.normalize) were randomly sampled from the training datasets with equal probability and used in training if at least 0.1% of voxels were included in the binary mask generated during pre-processing of molecular signals (i.e. 1048.6 voxels of the 64x128x128 voxel patches). Training data was augmented by mirroring and transposing in *xy*-direction on structural and molecular channels. In addition, we applied intensity augmentation on the structural channel with (0.9, 1.1, -0.1, 0.1) for the two scaling and the two intensity shifting parameters, respectively, and enabled range clipping.

*Prediction*
Prediction was performed on ISTA's high-performance computing cluster using SLURM for task allocation. We requested 8 CPUs, graphics acceleration by 1 GPU (Nvidia GeForce RTX 2080Ti or similar) and 100 GB of RAM. Prediction was performed in a piece-wise, "scanning" mode, again with 64x128x128 voxel patch size after converting LICONN structural data to zarr format while applying the same normalization procedure as for the training data.
To convert predicted molecule locations to point annotations, we first rescaled intensity using the skimage.exposure.rescale_intensity, taking p1 and p99.95 as limits, with p1 referring to the first percentile and p99.95 to the 99.95$^{th}$ percentile. We then applied 3D-Gaussian filtering with σ=5 for Bassoon and Shank2. We applied Otsu thresholding to derive a binary mask and skimage.measure.label to extract instance segmentations. We erased volume segments smaller than 60 voxels and those that covered less than 13 imaging planes. We applied the skimage.measure.regionprops function to determine the weighted centroid for each segment based on the intensity of the underlying LICONN structural channel and placed a point annotation at that location.
Association of pre- and post-synaptic sites and validation of deep-learning based synapse prediction were

Visualization
Neurite and synaptic segmentations, skeletons, and imaging volumes were visualized in 3D with Blender versions 2.92, 3.2, or 3.51 (https:// www.blender.org/). Video files were also generated with Blender. For visualization of neurites on an overview scale (e.g. Fig. 1A, C,D), precomputed meshes from Neuroglancer were used. Detailed views of 3D-renderings (e.g. Fig. 1A zoomed dendrite, Fig. 1E,F, 3C) were based on meshes generated with the marching cubes implementation in scikit-image. For 3D rendering of molecular signals in the context of neuronal segmentations, immunolabeling channels and predicted molecule distributions were intensity scaled according to visual inspection, background was removed with σ≈0.5 and σ≈5 to preserve shapes and then Otsu thresholded. To associate these signals with a particular neuronal segment, the segment was expanded with a ball of radius 6 and any molecular signals with at least 1 voxel overlap were retained. Meshes were again generated with the marching cubes algorithm.

Associating synapse detections with FFN derived neuronal segments
To map detected synapses onto selected neuronal segments, we first expanded the segment with a ball of radius of 5 or 6 voxels. To map post-synaptic sites onto dendrites, we selected those

detections whose centre coordinates were located within the expanded neuronal segment. We recalled the corresponding pre-synaptic partners using the matching matrix $M_{ij}$ to generate the list of full synapses associated with that segment. For axons, we first identified the pre-synapses located within the expanded segment, and then retrieved the corresponding post-synapses from the matching matrix. Due to the large size of the imaging volume, computations were performed on manually defined patches, and results were combined for visualization and further analysis.

Automated analysis of connectivity based on deep-learning based synapse prediction and neurite segmentation

To analyze connectivity in Fig. 4P, we first performed deep-learning based synapse prediction on the imaging volume. For this, we predicted Bassoon and Shank2 molecule locations in smaller overlapping patches of the volume, and combined them into the final full-size prediction volume. Next, we converted the predicted molecule locations into synaptic point annotations using the algorithm described above. Due to the large size of the volume, pre-synapse and post-synapse annotations were computed on manually defined non-overlapping patches. Association of pre- and post-synapses, i.e. computation of the matching matrix $M_{ij}$, was performed on the entire volume to avoid errors in full synapse assembly due to separation by a patch boundary. We then associated pre- and post-synaptic annotations with the neuronal segmentation.

For each pre-synapse (post-synapse) location, we identify the underlying segment ID from the FFN-based neurite segmentation. If no segment was present at the coordinates of a specific pre- or postsynaptic detection, we performed an automated search in a single plane for the nearest non-zero segment in a 20x20 voxel neighborhood. If i) the squared distance between the respective pre-synapse (post-synapse) and the nearest non-zero segment was less than 30 voxel$^2$ and ii) the segment was classified as an axon (dendrite) by the neurite classifier described above, the respective pre-synapse (post-synapse) was mapped to the segment.

Finally, we combined the information from the matching matrix $M_{ij}$ and the synapse-segment associations obtained in the previous step to compute a matrix of connectivity between neuronal segments. For this, we iterated through the matching matrix $M_{ij}$ and added a connection between the segment associated with post-synapse i and the segment associated with pre-synapse j.

Statistics

GraphPad Prism version 10.1.2 was used for statistical analysis and for plotting. Bootstrap analysis in fig. S24 was performed according to (*11*) based on code available at https://gitlab.mpcdf.mpg.de/connectomics/human_primate.

Creation of figures

Figures were created in Adobe Illustrator version 27.7 The schematic in Fig. 5D was generated with Biorender (biorender.com).

### Data and materials availability

All relevant code will be made publicly available through an appropriate open-source platform (e.g. GitHub) upon publication of the manuscript.

Original data and segmentations for two example datasets are available in browsable format at the following locations:

**Fig. 1A-B**: https://neuroglancer-demo.appspot.com/#!gs://liconn-public/ng_states/fig1a.json
**Fig. 2A-F**: https://neuroglancer-demo.appspot.com/#!gs://liconn-public/ng_states/expid82.json

**Fig. 2C**: https://neuroglancer-demo.appspot.com/#!gs://liconn-public/ng_states/expid82_fig2_axons.json
**Fig. 2D**: https://neuroglancer-demo.appspot.com/#!gs://liconn-public/ng_states/expid82_fig2_dends.json

Original data for a LICONN volume fused from consecutive overlapping imaging slabs obtained via hydrogel sectioning is available here:
**Fig. S31:** https://neuroglancer-demo.appspot.com/#!gs://liconn-public/ng_states/expid146.json

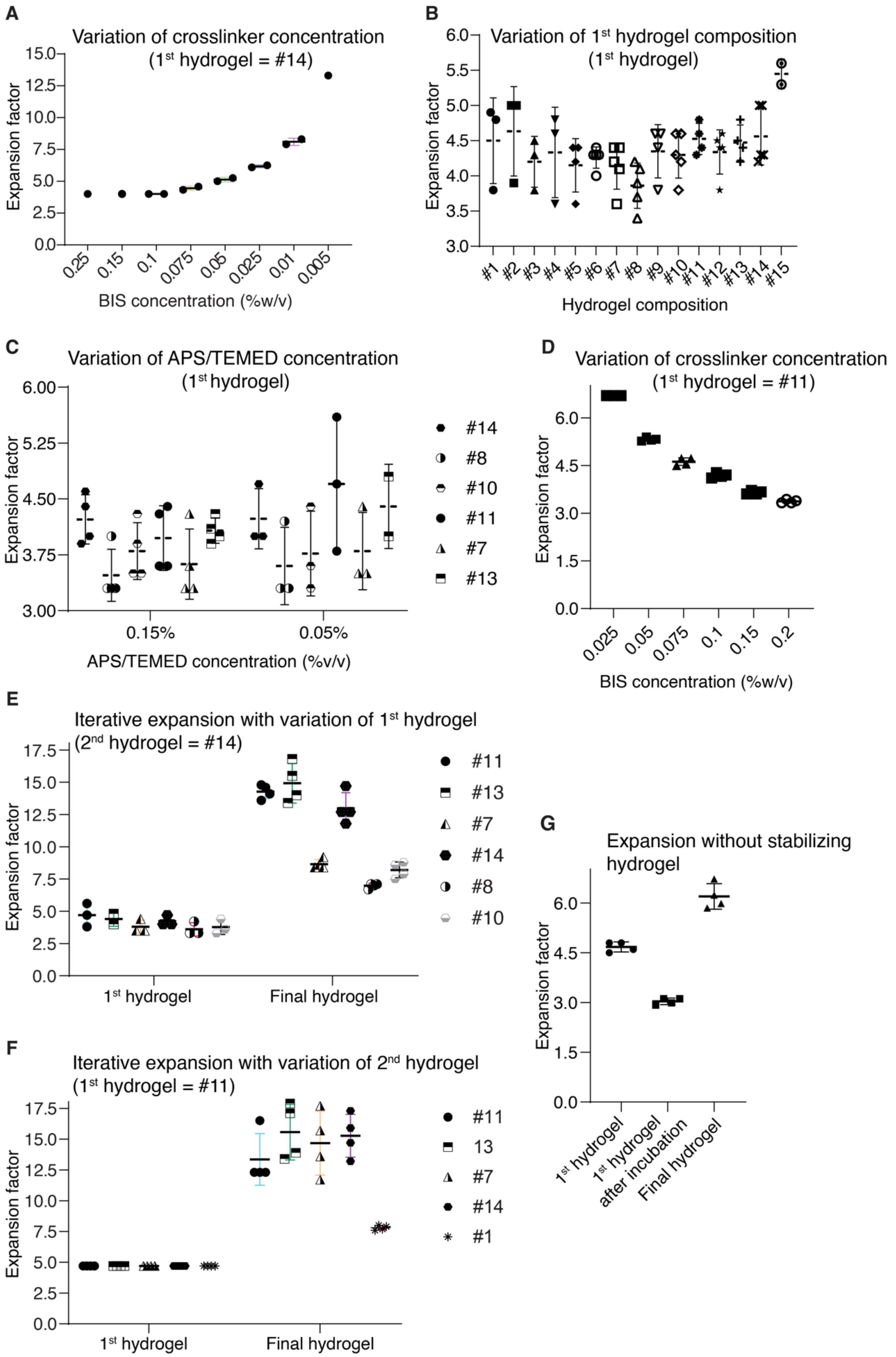
A
Variation of crosslinker concentration
(1st hydrogel = #14)
Expansion factor
BIS concentration (%w/v)
0.25 0.15 0.1 0.075 0.05 0.025 0.01 0.005
B
Variation of 1st hydrogel composition
(1st hydrogel)
Expansion factor
Hydrogel composition
#1 #2 #3 #4 #5 #6 #7 #8 #9 #10 #11 #12 #13 #14 #15
C
Variation of APS/TEMED concentration
(1st hydrogel)
Expansion factor
APS/TEMED concentration (%v/v)
0.15% 0.05%
#14 #8 #10 #11 #7 #13
D
Variation of crosslinker concentration
(1st hydrogel = #11)
Expansion factor
BIS concentration (%w/v)
0.025 0.05 0.075 0.1 0.15 0.2
E
Iterative expansion with variation of 1st hydrogel
(2nd hydrogel = #14)
Expansion factor
1st hydrogel Final hydrogel
#11 #13 #7 #14 #8 #10
F
Iterative expansion with variation of 2nd hydrogel
(1st hydrogel = #11)
Expansion factor
1st hydrogel Final hydrogel
#11 13 #7 #14 #1
G
Expansion without stabilizing hydrogel
Expansion factor
1st hydrogel
1st hydrogel after incubation
Final hydrogel

**Fig. S1. Optimization of hydrogel compositions. (A)** Expansion factor in single-step expansion as a function of crosslinker (BIS) concentration. Hydrogel composition as in hydrogel #14 (see Table S7) but with variation of BIS concentration. Reduction of crosslinker concentration increased exF but resulted in mechanically unstable hydrogels. **(B)** ExF of the first expandable hydrogel for different hydrogel compositions according to Table S7. Hydrogel composition #11 was chosen for the first expandable hydrogel in LICONN, taking additional factors like expansion fidelity and mechanical stability into account. For example, hydrogel #15 resulted in a mechanically unstable hydrogel. **(C)** Expansion factor of the first expandable hydrogel for different concentrations of polymerization initiator (APS) and accelerator (TEMED), used at equal concentrations, for different hydrogel compositions according to Table S7. **(D)** Evaluation of expansion factor for the hydrogel composition of the first expandable hydrogel (composition #11) in LICONN, as a function of crosslinker concentration. **(E)** Expansion factor after the first and second expansion steps in iterative expansion as a function of composition of the first expandable hydrogel. The composition of the second expandable hydrogel was kept constant, using the one chosen for LICONN (#14). **(F)** Expansion factor after the first and second expansion steps in iterative expansion as a function of composition of the second expandable hydrogel. The composition of the first expandable hydrogel was kept constant, using the one chosen for LICONN (#11). **(G)** Effect of omitting the stabilizing hydrogel. Expansion factor after the first expansion step, after incubation with the monomer solution for the second expandable hydrogel (when omitting the stabilizing hydrogel), and after polymerizing and expanding the second swellable hydrogel. Expansion factor was drastically reduced with respect to the LICONN procedure. Compositions of first and second expandable hydrogels were identical to the LICONN procedure. Expansion factors were evaluated by measuring hydrogel size with a caliper in technical samples without biological specimens. Graphs represent mean ± s.d., data points represent individual technical replicates.

| ID | Acrylamide (%w/v) | Sodium acrylate (%w/v) | N,N´-Methylenebisacrylamide (%w/v) |
|---|---|---|---|
| | AA | SA | BIS |
| #1 | 4 | 5.3 | 0.075 |
| #2 | 4 | 7 | |
| #3 | 6 | 7 | |
| #4 | 7 | 7 | |
| #5 | 10 | 7 | |
| #6 | 12.5 | 7 | |
| #7 | 15 | 7 | |
| #8 | 20 | 7 | |
| #9 | 10 | 10 | |
| #10 | 14 | 10 | |
| #11 | 10 | 12.5 | |
| #12 | 12.5 | 12.5 | |
| #13 | 10 | 15 | |
| #14 | 10 | 19 | |
| #15 | 2.5 | 8.6 | |
| #16 | 12.5 | 10 | |

**Table S7. Composition of hydrogels.** Hydrogels used in various optimization steps as indicated in the respective Supplementary Figures. For LICONN, compositions #11 and #14 were chosen for the first and second expandable hydrogels, respectively.

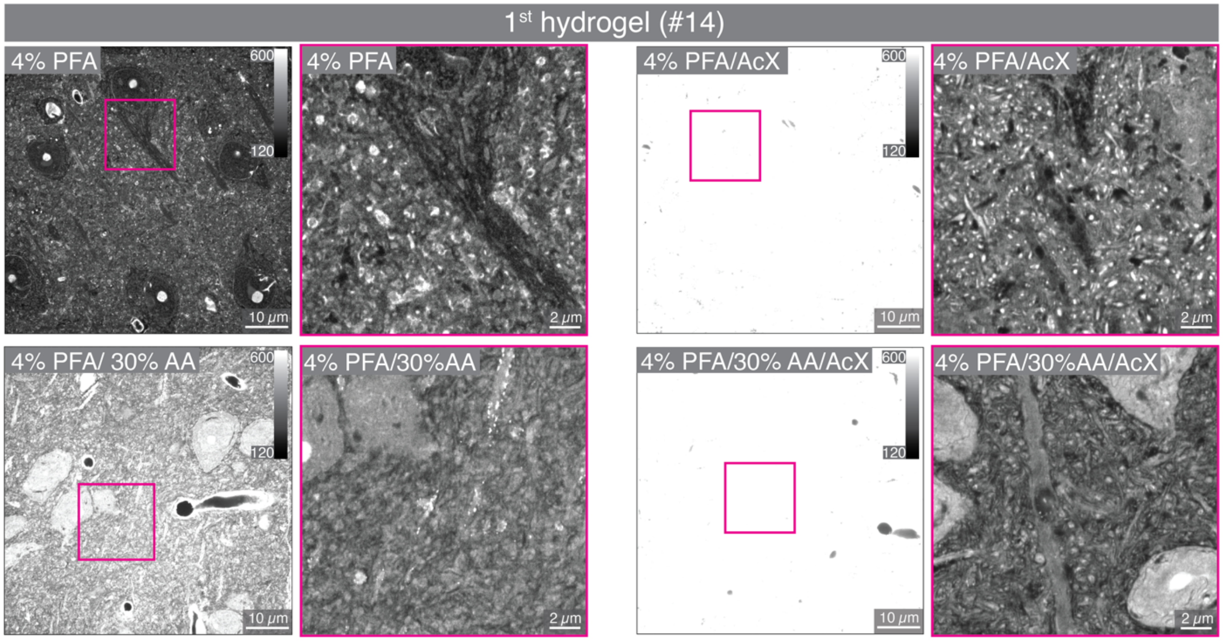


**Fig. S2. Initial evaluation of protein anchoring.** Confocal images with magnified views after first hydrogel expansion using hydrogel composition #14 (Table S7). Imaging followed transcardial fixative perfusion, postfixation, quenching with glycine, slicing, optional additional anchoring with acryloyl-X (AcX, 100 µg/ml), hydrogel embedding, denaturation, protein density (pan) labeling, and expansion. *Left, top*: transcardial fixative perfusion with 4% PFA in 1X PBS. *Left, bottom*: Transcardial fixative perfusion with 2% AA in 1X PBS followed by 30% AA and 4% PFA in 1X PBS. Addition of AA increased signal level (same imaging parameters and intensity lookup table in all overview images; adjusted intensity ranges in magnified views). *Right*: Additional anchoring with AcX strongly increased signal level (intensity range saturated in overview images) both for PFA and PFA+AA perfusion. However, we decided against AcX as anchoring agent, as the strong anchoring was paralleled with compromises in preservation of cellular structure. Data representative of *n*=3 technical replicates.

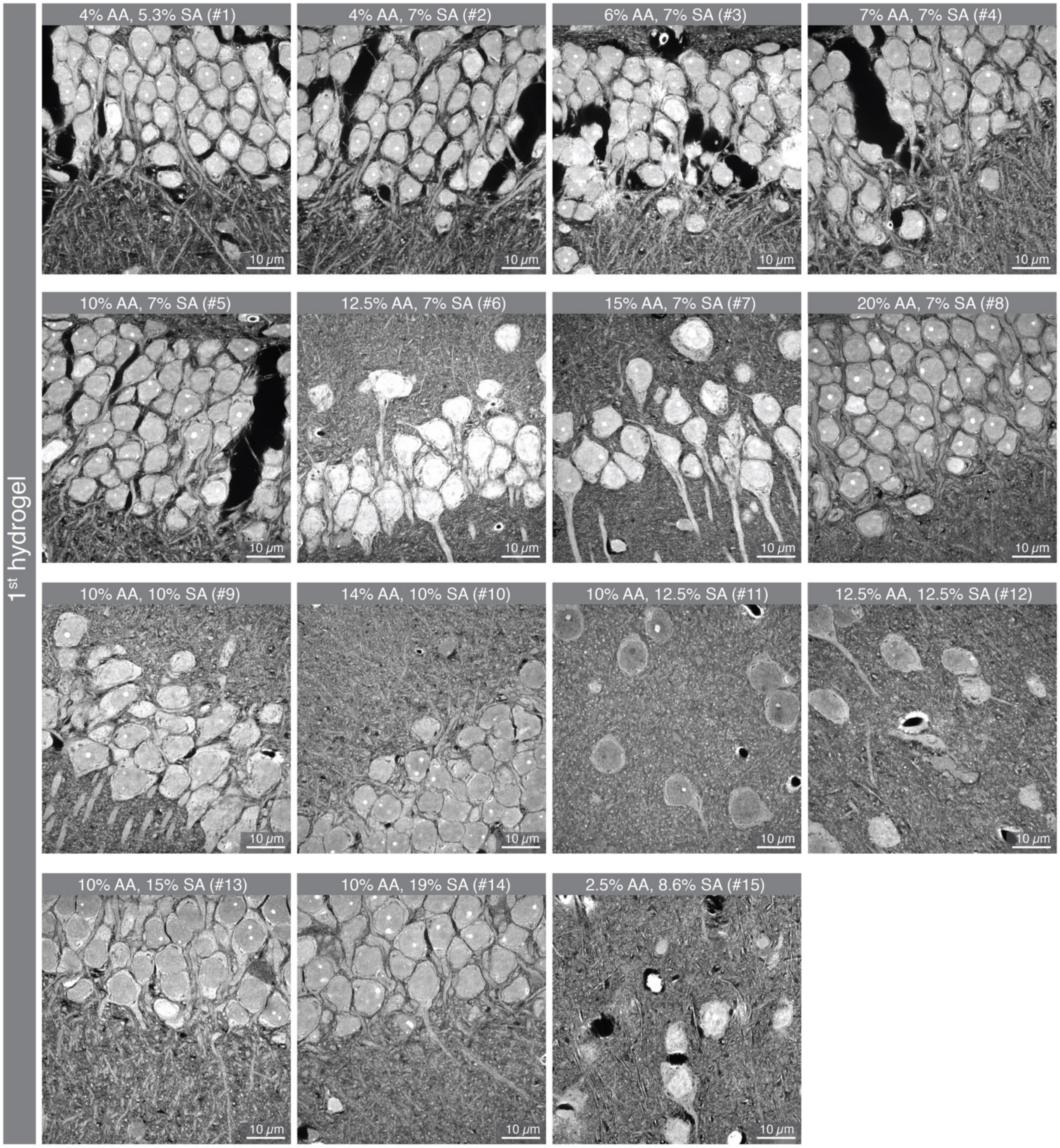


**Fig. S3. Tissue preservation as a function of the composition of the first expandable hydrogel.** Confocal images after the first expansion step for different hydrogel compositions according to Table S7 in hippocampus (#1, #2, #3, #4, #5, #6, #7, #8, #9, #10, #13) or cortex (#11, #12, #15). For transcardial fixative perfusion, previously used conditions were employed in this measurement (perfusion with 2% AA in PBS, followed by 30% AA and 4 % PFA in PBS) (*12*). Brains were post-fixed overnight in the same solution at 4 °C. After slicing, samples were quenched with glycine and incubated with acryloyl-X (100 µg/ml in PBS, overnight) for anchoring of proteins to the hydrogel. Denaturation was performed for 100 min at 95 °C in the same denaturation buffer as used for LICONN experiments. Protein density (pan) labeling and expansion were performed before imaging. Hydrogel compositions with low AA concentrations resulted in fragile hydrogels and ruptures of the tissue-hydrogel hybrid after expansion (black areas). Each condition was replicated at least $n$=2 times. Hydrogels #14 and #15 were replicated $n$=5 times.

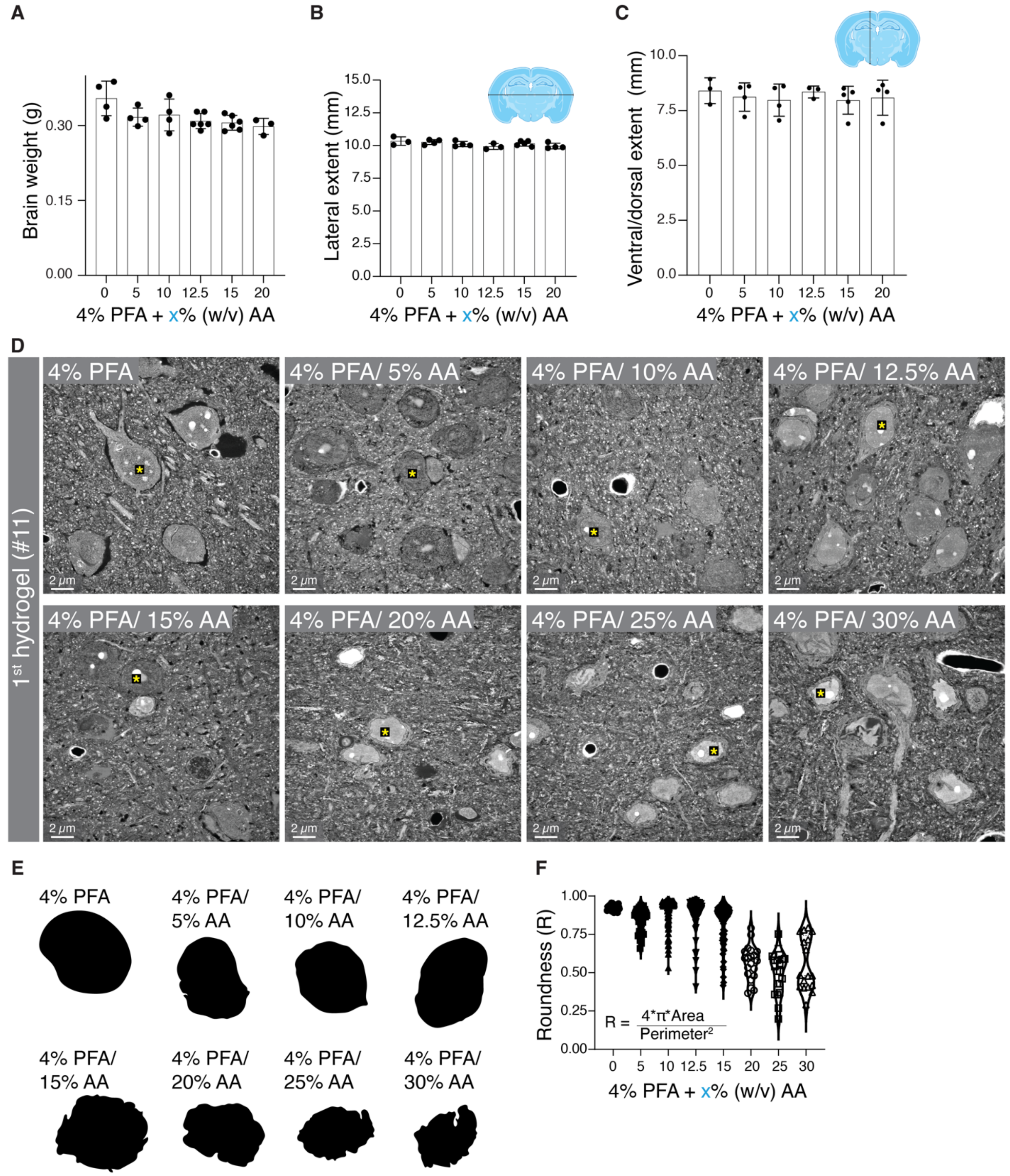


**Fig. S4. Optimization of transcardial fixative perfusion conditions. (A-C)** Weight, lateral, and ventral/dorsal extent of brains harvested after transcardial fixative perfusion, first with 1X PBS and then with 4% PFA and varying concentrations of AA. Weight shows a tendency towards lower values with increasing AA concentration. Brain weight was determined after overnight postfixation in the same solution as used in perfusion and washing in 1X PBS. For quantification of dimensions (see schematic), brains were sliced, stained with DAPI and mounted in Mowiol on a microscopy slide before confocal imaging of corresponding coronal slices across the various conditions. Individual data points represent individual animals (*n*=3 to *n*=6). **(D)** Confocal images after overnight postfixation in the same solution, slicing, quenching with glycine, hydrogel embedding (composition #11), denaturation at 95 °C for 100 min, protein density (pan) labeling, and expansion. No additional protein anchoring to the hydrogel was applied. Omission of AA in the transcardial perfusion solution led to voids around cells whereas the highest AA concentrations led to corrugated cell and nuclear outlines. **(E)** Shapes of the cell nuclei indicated by asterisks in (D). **(F)** Quantification of the roundness R of nuclei according to $R=4\pi*area/perimeter^2$. Data points represent individual nuclei assessed across *n*=3 or more biological replicates.

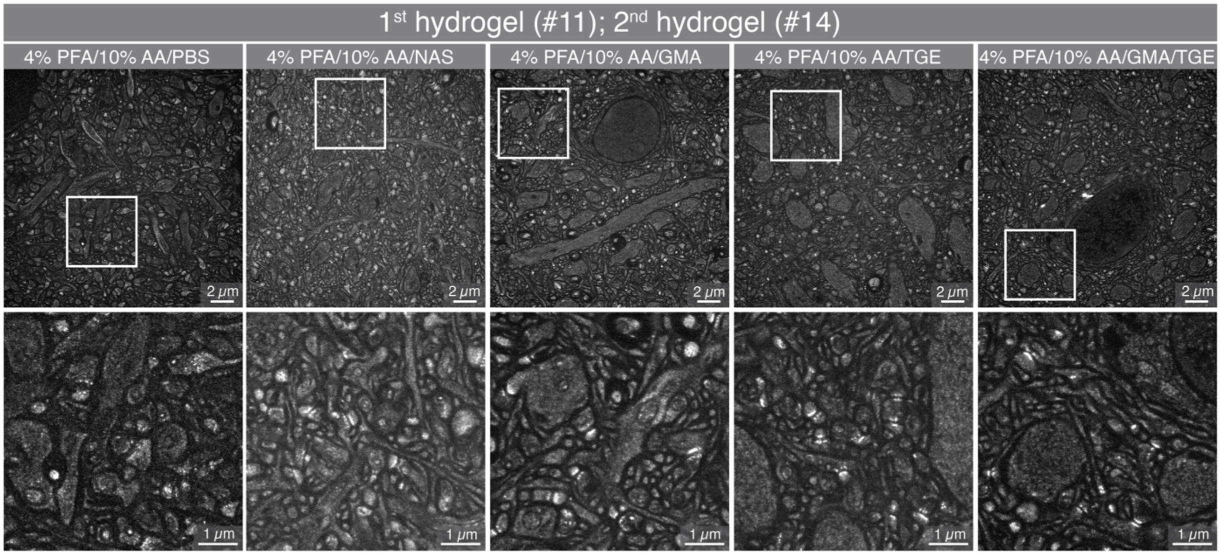


**Fig. S5. Effect of NAS anchoring and epoxides evaluated after the second expansion step.** Overview confocal images and magnified regions as indicated by white boxes in hippocampal CA1 stratum radiatum. The transcardial fixative perfusion procedure (4% PFA and 10 % AA in 1X PBS) and all subsequent steps were performed according to the final LICONN parameters (1$^{st}$ expandable hydrogel: composition #11, 2$^{nd}$ expandable hydrogel: composition #14). After postfixation, slicing and quenching, one of the following steps were performed from left to right: (i) no additional anchoring, (ii) anchoring with NAS (360 μM) for 3h at room temperature, (iii) application of epoxide GMA (3h, 37 °C), (iv) application of epoxide TGE (3h, 37 °C), (v) application of GMA and TGE (3h, 37 °C). Intensity lookup tables were adjusted to account for different overall signal intensities in the images. Both NAS anchoring and epoxide treatment produced high quality datasets whereas omission of these reagents led to reduced signal levels and poorer structural representation. We opted for epoxide treatment as this led to improved signal-to-noise ratio and more pronounced delineation of high protein-density features at synapses. Representative of *n*=3 technical replicates.

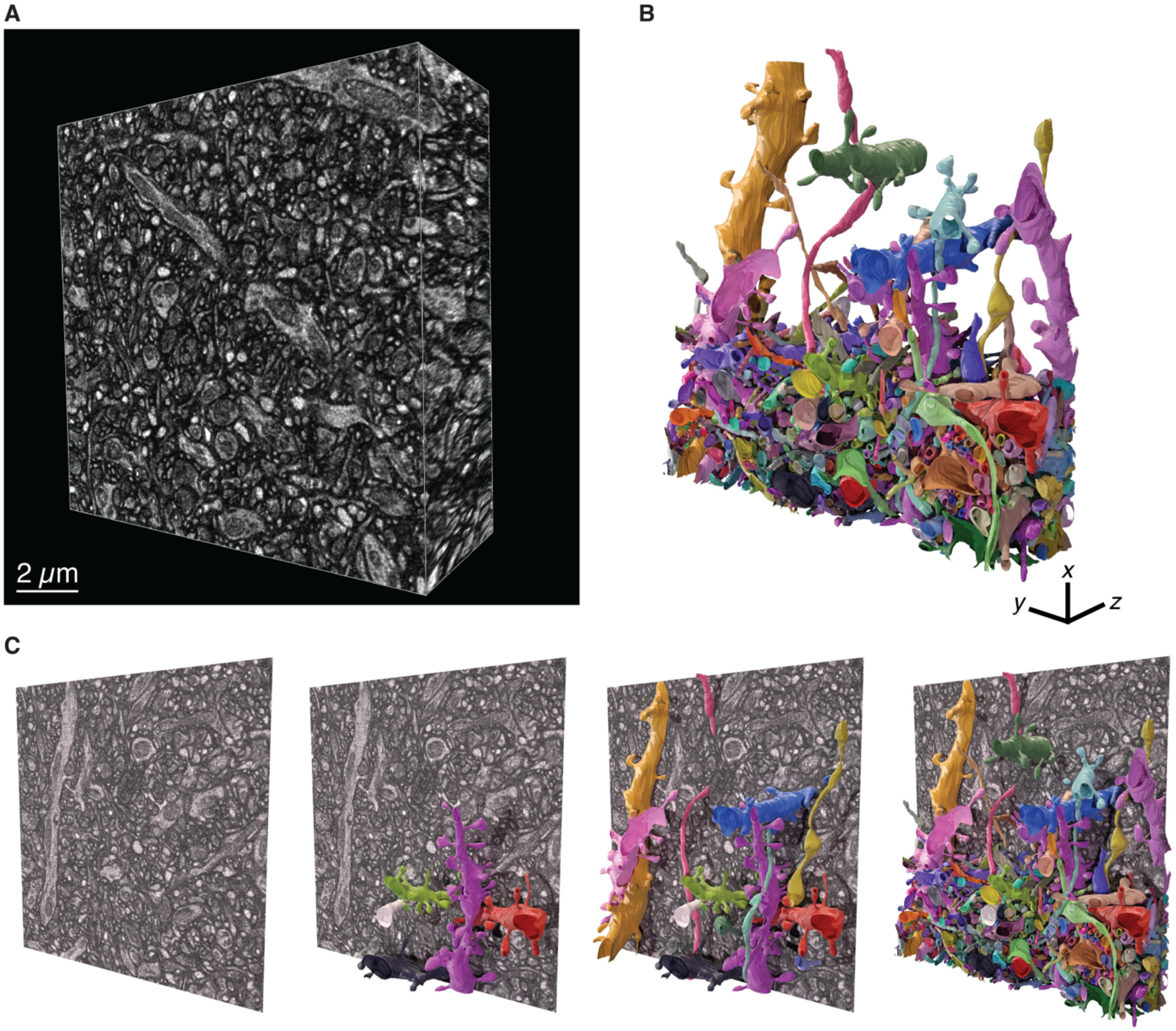


**Fig. S6. Manual annotation of LICONN volume with NAS anchoring. (A)** 3D rendering of a 13 x 13 x 5 μm$^3$ LICONN volume with NAS anchoring (different from dataset in Fig. 1G) in the hippocampal CA1 stratum oriens. **(B)** Rendering of manual annotations of a subset of neuronal structures. The bottom third was densely annotated. **(C)** Single confocal imaging plane with progressively increasing number of rendered segments. Transcardial fixative perfusion was performed with 4% PFA, 10 % AA in 1X PBS, followed by postfixation, slicing, glycine quenching, NAS anchoring and hydrogel embedding. The first and second expandable hydrogels had compositions #13 and #14, respectively, according to Table S7.

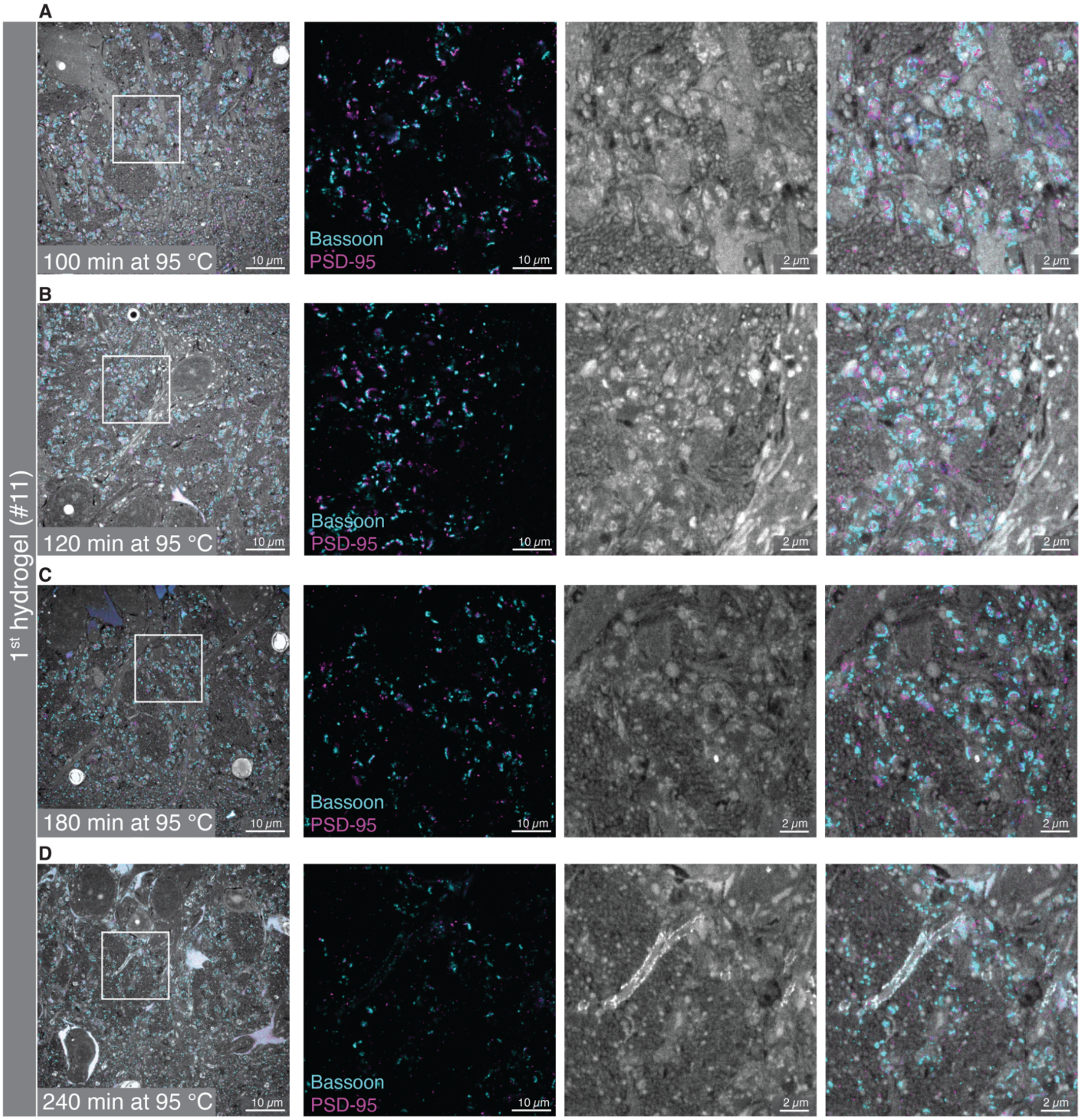


**Fig. S7. Optimization of denaturation duration.** (**A-D**) Confocal images in the hippocampal CA3 stratum lucidum after the first hydrogel expansion step in the LICONN procedure, with protein density (pan) labeling (gray) and immunolabeling for Bassoon (cyan) and PSD-95 (magenta). Denaturation time was varied. Increasing denaturation time led to signal reduction of the protein-density labeling and immunolabeling and compromised structural integrity. Representative of *n*=3 technical replicates.

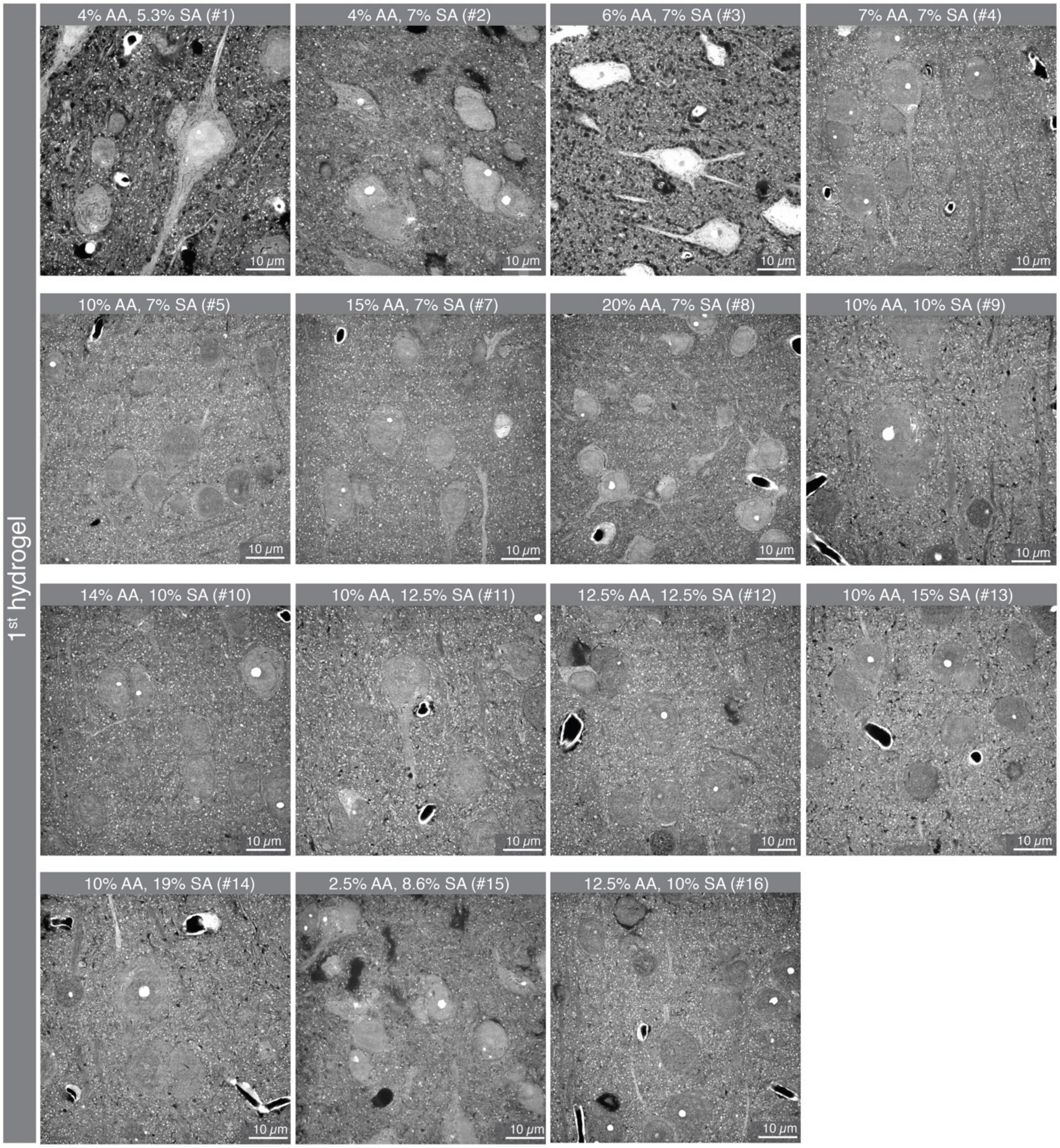


**Fig. S8. Evaluation of the composition of the first expandable hydrogel, using optimized perfusion and epoxide treatment.** Representative confocal images in cortex after the first expansion step and protein density (pan) labeling (gray). Composition of the first expandable hydrogel was varied according to Table S7 while other parameters were identical as in the LICONN procedure. Hydrogels with low AA concentration (e.g. #1, #2, #3, #15) showed compromised integrity and voids throughout the tissue hydrogel hybrid. Representative of *n*=2 technical replicates.

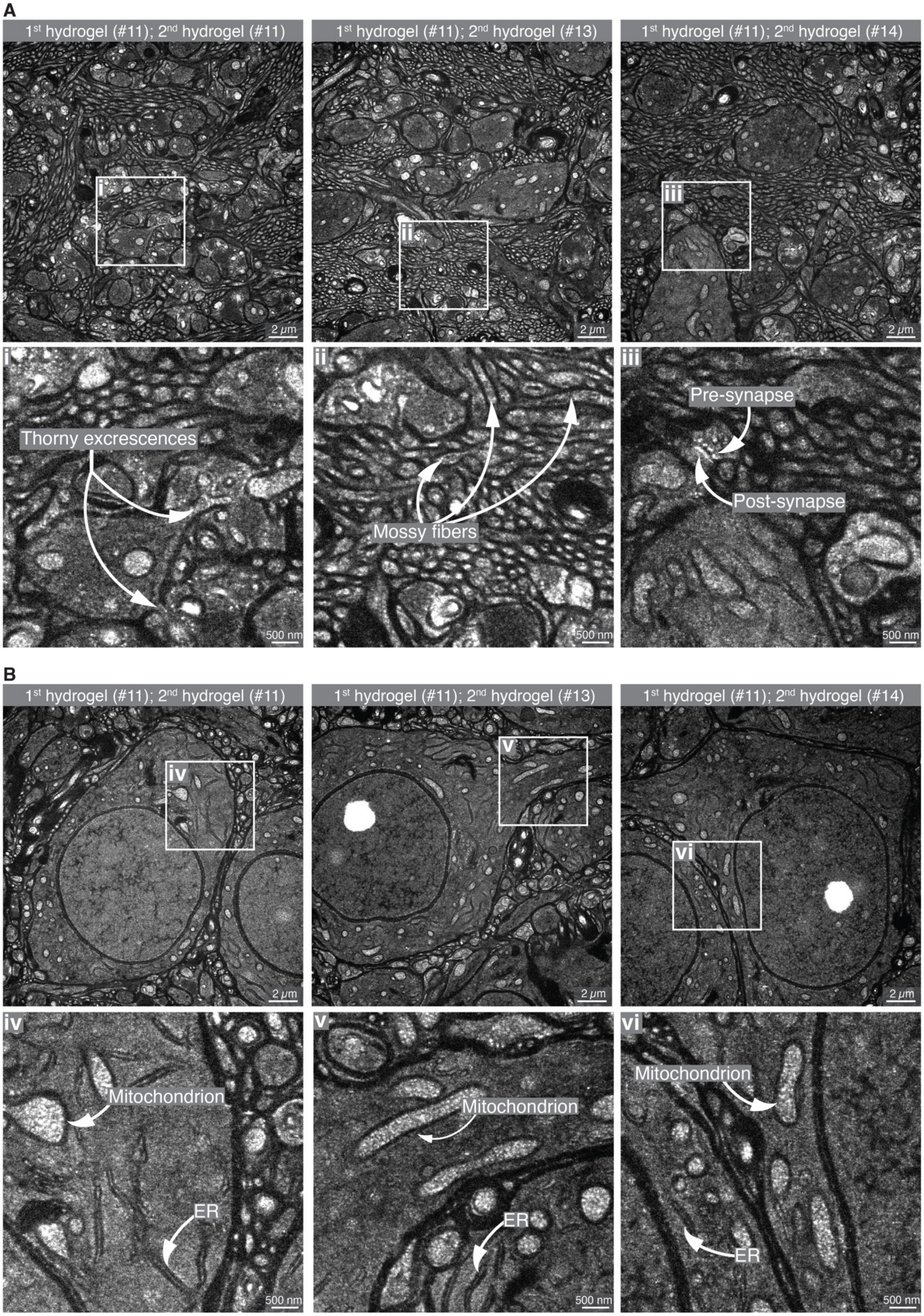
A
1st hydrogel (#11); 2nd hydrogel (#11)
1st hydrogel (#11); 2nd hydrogel (#13)
1st hydrogel (#11); 2nd hydrogel (#14)
i
ii
iii
2 μm
Thorny excrescences
Mossy fibers
Pre-synapse
Post-synapse
500 nm
B
1st hydrogel (#11); 2nd hydrogel (#11)
1st hydrogel (#11); 2nd hydrogel (#13)
1st hydrogel (#11); 2nd hydrogel (#14)
iv
v
vi
2 μm
Mitochondrion
ER
500 nm

**Fig. S9. Robustness against variation of parameters.** Confocal images after full expansion with optimized denaturation duration, first expandable and stabilizing hydrogels. Perfusion was performed with 4% PFA and 5 % AA in 1X PBS. For protein anchoring, NAS was used. The second expandable hydrogel was varied with compositions according to Table S7. **(A)** Overview and magnified views in the hippocampal CA3 stratum lucidum. Here, mossy fibers, the axons of dentate gyrus (DG) granule cells, are organized in bundles and form excitatory synapses with complex dendritic spines dubbed "thorny excrescences" on proximal dendrites of CA3 pyramidal neurons. **(B)** Similar measurements at cell somata in the neighboring CA3 stratum pyramidale. All displayed conditions produced high-quality datasets. Representative of $n$=3 technical replicates for each condition.

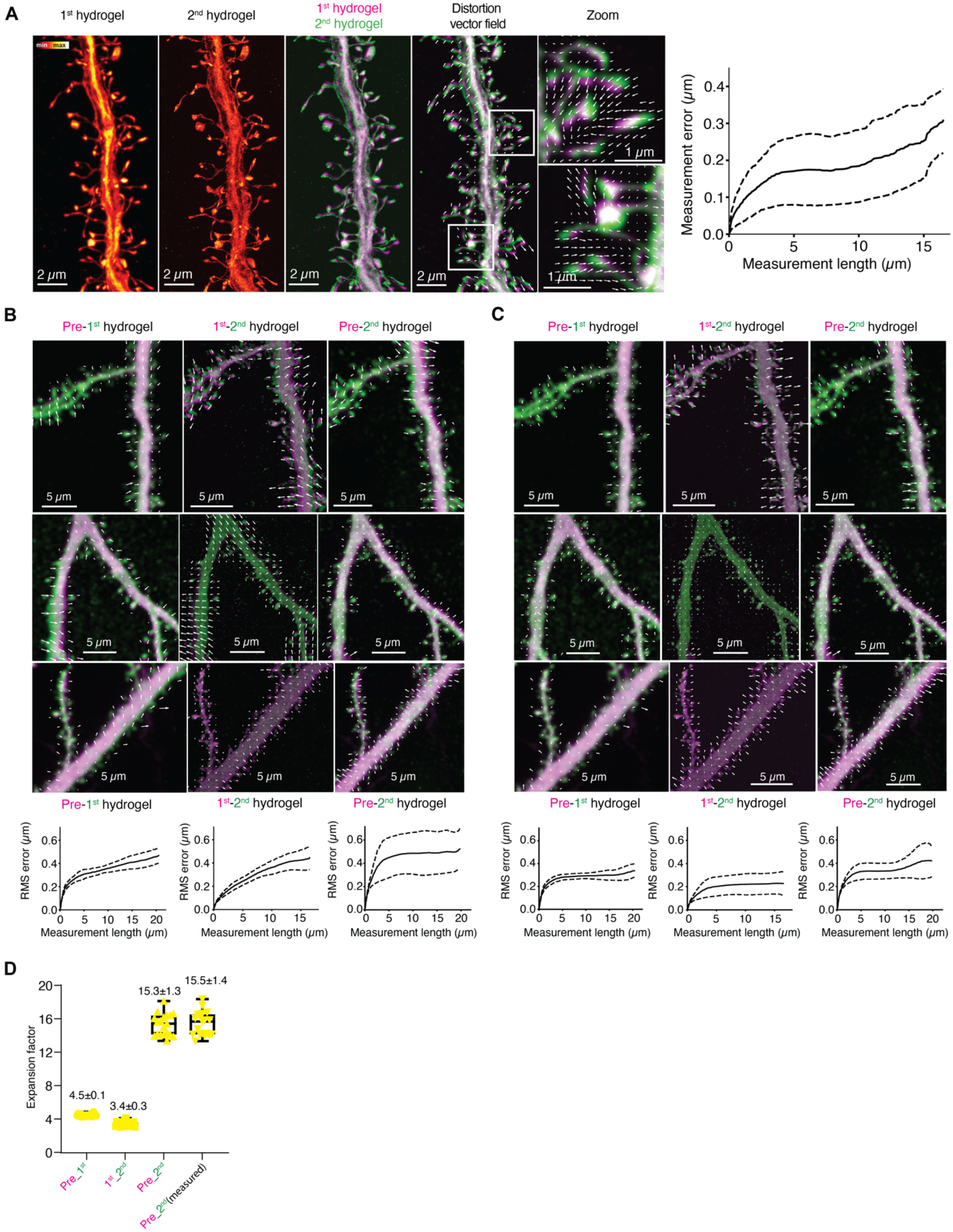


**Fig. S10. Analysis of expansion-induced distortions and expansion factor. (A)** Example of distortion analysis, illustrated here for the second LICONN expansion step. *Left to right*: (i)

Maximum intensity projection of a confocal imaging volume after the first LICONN expansion step in cortex of a Thy1-eGFP mouse, with an eGFP expressing dendrite stretch. The cytosolically expressed eGFP was visualized by immunolabeling. (ii) Maximum intensity projection of the same region after the second LICONN expansion step. Maximum intensity projections were aligned with a similarity transformation (including isotropic scaling, translation and rotation). (iii) Overlay. (iv) Vector field of distortions during the second expansion step with two magnified views, overlaid with the respective maximum intensity projections. Beyond distortions introduced by the expansion step itself, they may also include additional distortions from manual handling and mounting for imaging. Gaussian smoothing was applied to achieve similar appearance of images before and after expansion for distortion analysis (see materials and methods). (v) Distortions as a function of measurement length for this specific measurement. Scale bars: 2 µm, magnified views: 1 µm. **(B)** Distortion vector fields for the first (*left*: pre-expansion to1$^{st}$ hydrogel) and second (*middle*: 1$^{st}$ to 2$^{nd}$ hydrogel) individual expansion steps and for the overall LICONN procedure (right: pre-expansion to 2$^{nd}$ hydrogel). Images represent three examples of measurements used in the analysis. Color coding in the maximum intensity projections is according to the respective figure headings. Similarity transformation and Gaussian smoothing were applied as in (A). *Bottom*: Root mean square (RMS) measurement error over different measurement lengths for the first and second expansion steps and for the overall LICONN procedure, evaluated in 14 fields of view in 4 technical replicates, recorded in cortex across *n*=3 animals. Scale bars: 5 µm. **(C)** Similar analysis using an affine transformation (including scaling, translation, rotation and shearing) for initial alignment of maximum intensity projections. **(D)** Expansion factor determined as the linear scaling factor in the similarity transformation as in panel (A) in the 14 fields of view analyzed across 4 technical replicates in *n*=3 animals). ExF was evaluated for the first (pre_1$^{st}$) and second (1$^{st}$_2$^{nd}$) expansion steps individually. The value obtained as product (pre_2$^{nd}$) of the exF in the first and second steps was consistent with the value obtained when directly aligning images of pre-expansion and fully expanded hydrogels (pre_2$^{nd}$ (measured)). Scale bars, measurement errors and measurement lengths are scaled to the pre-expansion tissue size. For display purposes, the length of arrows in the distortion vector fields was multiplied by 1.5, while for the magnified views in panel A, the original arrow length is shown.

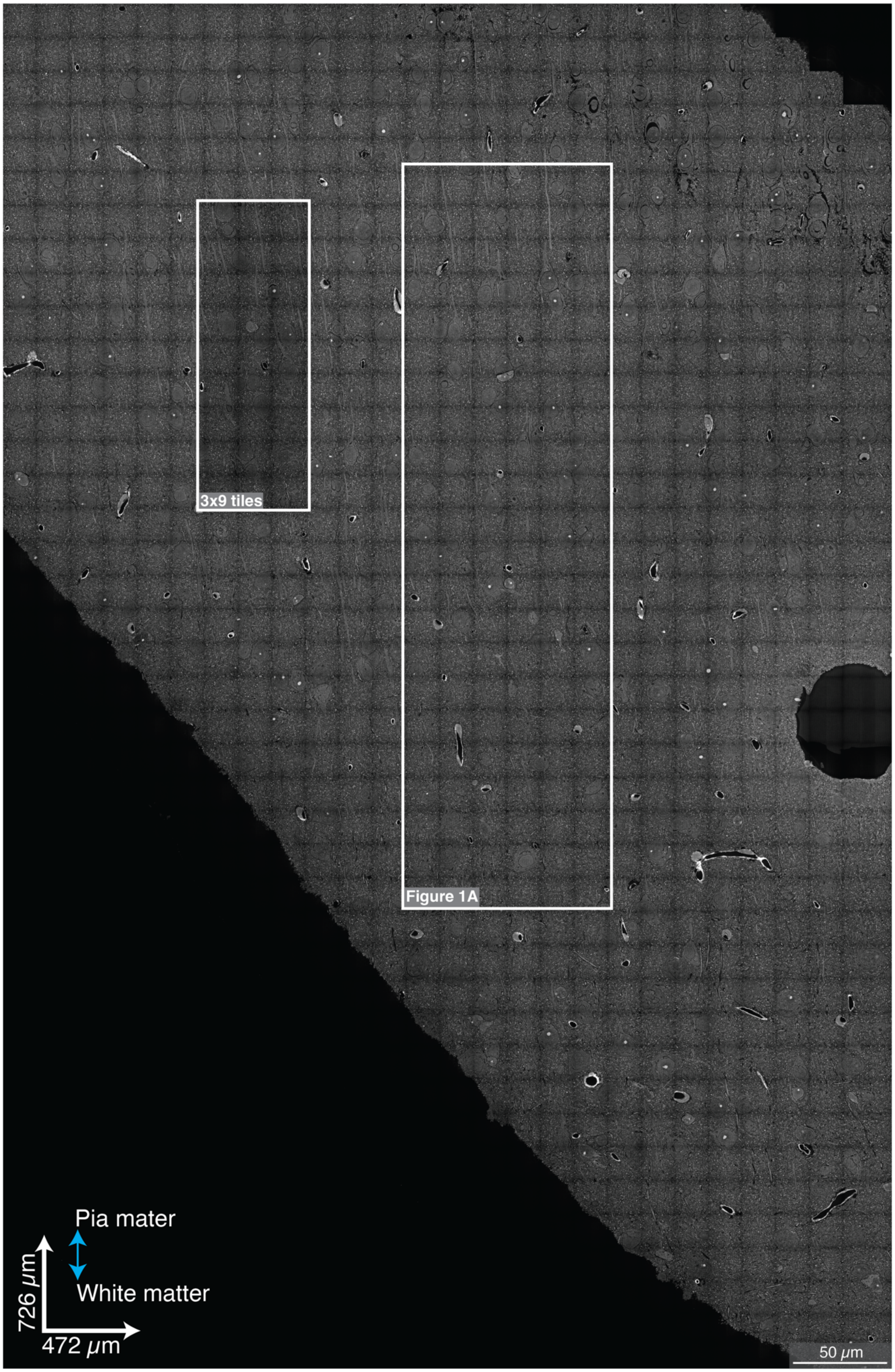
3x9 tiles
Figure 1A
Pia mater
White matter
726 $\mu$m
472 $\mu$m
50 $\mu$m

**Fig. S11. Overview imaging in cortex.** Single plane overview image in the region of the somatosensory cortex comprising the imaging volume in Fig. 1A as indicated by the boxed region. The lower signal in the region indicated on the left reflects photobleaching during previous recording of an imaging volume. The dark circular region in the right margin of the image represents a defect in the tissue-hydrogel hybrid, which can occasionally be observed in histological sample processing. Such defects are not specific to LICONN and did not impede our measurements, such that we did not screen samples for defects before imaging. If required for a particular application, such as scaling to much larger volumes, detecting defects would be possible before LICONN imaging by coarse overview scanning of the samples.

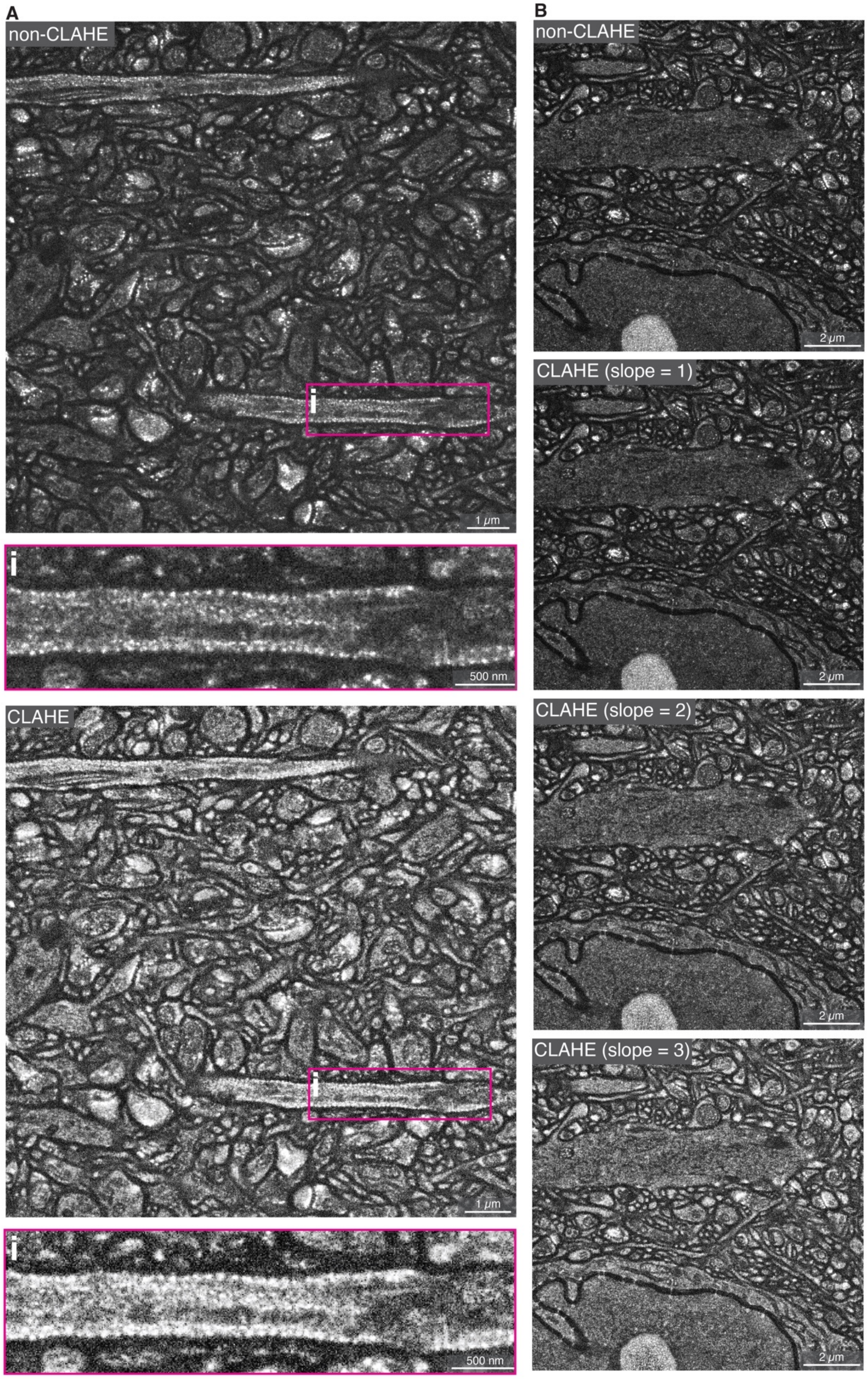
A
non-CLAHE
i
1 μm
i
500 nm
CLAHE
i
1 μm
i
500 nm
B
non-CLAHE
2 μm
CLAHE (slope = 1)
2 μm
CLAHE (slope = 2)
2 μm
CLAHE (slope = 3)
2 μm

**Fig. S12. Comparison of CLAHE-processed data with raw imaging data. (A)** *Top:* Raw (non-CLAHE) LICONN imaging data from somatosensory cortex (subregion of a single imaging plane from the dataset in Fig. 1A) with magnified view of boxed region, showing the periodic protein-density modulation induced by the actin-spectrin cytoskeletal rings present in a subset of neurites. *Bottom*: Same data after processing with contrast limited adaptive histogram equalization (CLAHE). High protein-density features at synapses and the periodic lattice stand out more clearly in the raw data. For automated neurite segmentation and display purposes, we used the CLAHE processed data as indicated. **(B)** Effect of CLAHE as a function of increasing slope parameter, as implemented in FIJI/ImageJ.

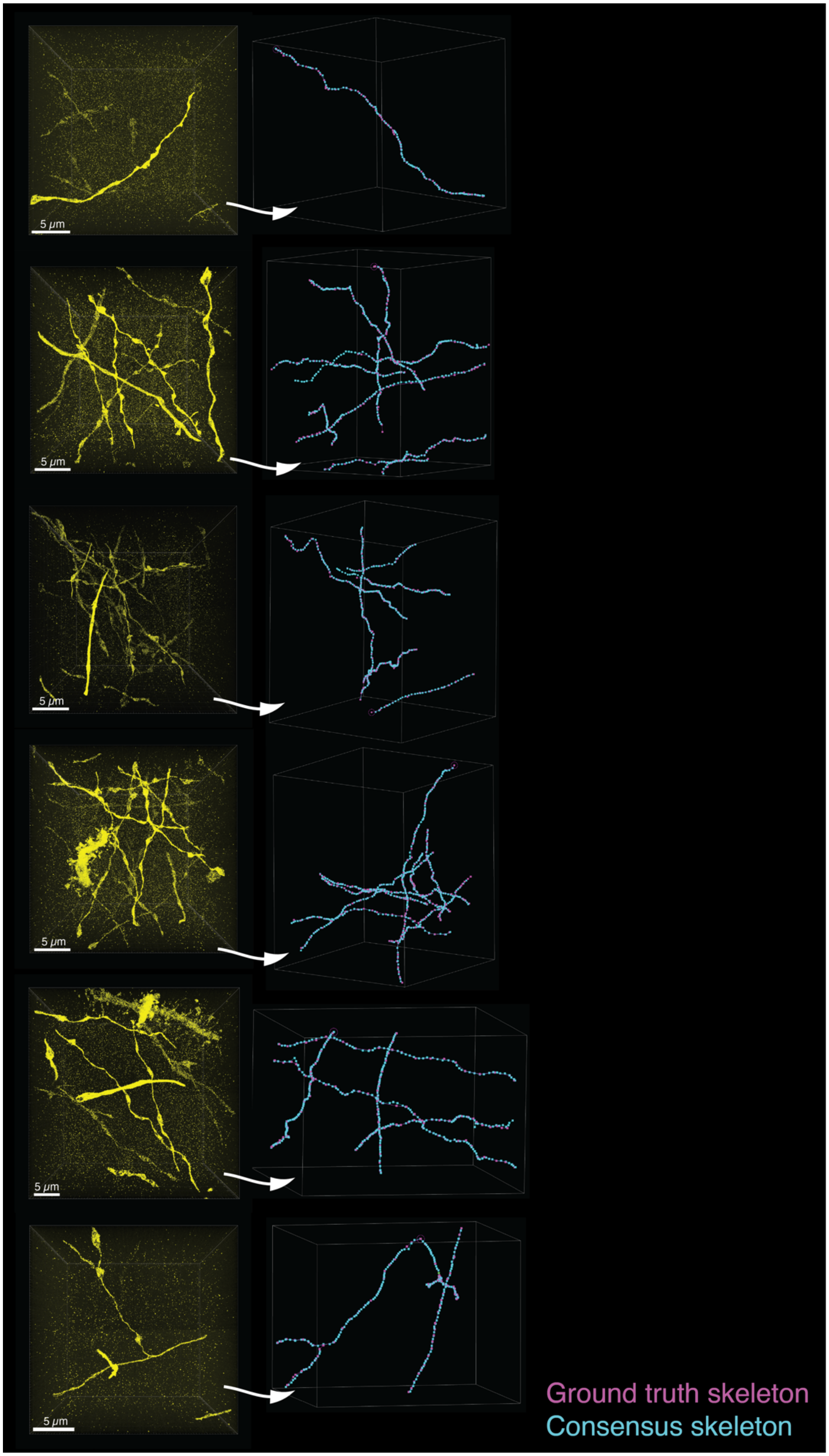
5 μm
5 μm
5 μm
5 μm
5 μm
5 μm
Ground truth skeleton
Consensus skeleton

**Fig. S13. Traceability of axons evaluated against ground truth from sparse eGFP expression (part 1).** *Left:* Volumetric renderings of eGFP signal (yellow) in LICONN volumes recorded in cortex of *Thy1-eGFP* animals with cytosolic expression of eGFP in a sparse subset of neurons. *Right*: Renderings of skeletons of the same structures generated from the LICONN structural channel (consensus skeletons, cyan) by two human annotators blinded to the eGFP channel. These are overlaid with skeletons that were generated by an independent annotator, taking both the LICONN structural channel and the eGFP signal into account (ground truth skeletons, magenta). Note that volume renderings for the eGFP signal and skeletons are from different camera positions. eGFP data was visualized with Imaris software. In total, 37 axon stretches (880 µm cumulative length of eGFP-expressing axons, recorded across *n*=3 technical replicates across *n*=2 animals in cortex) were analysed.

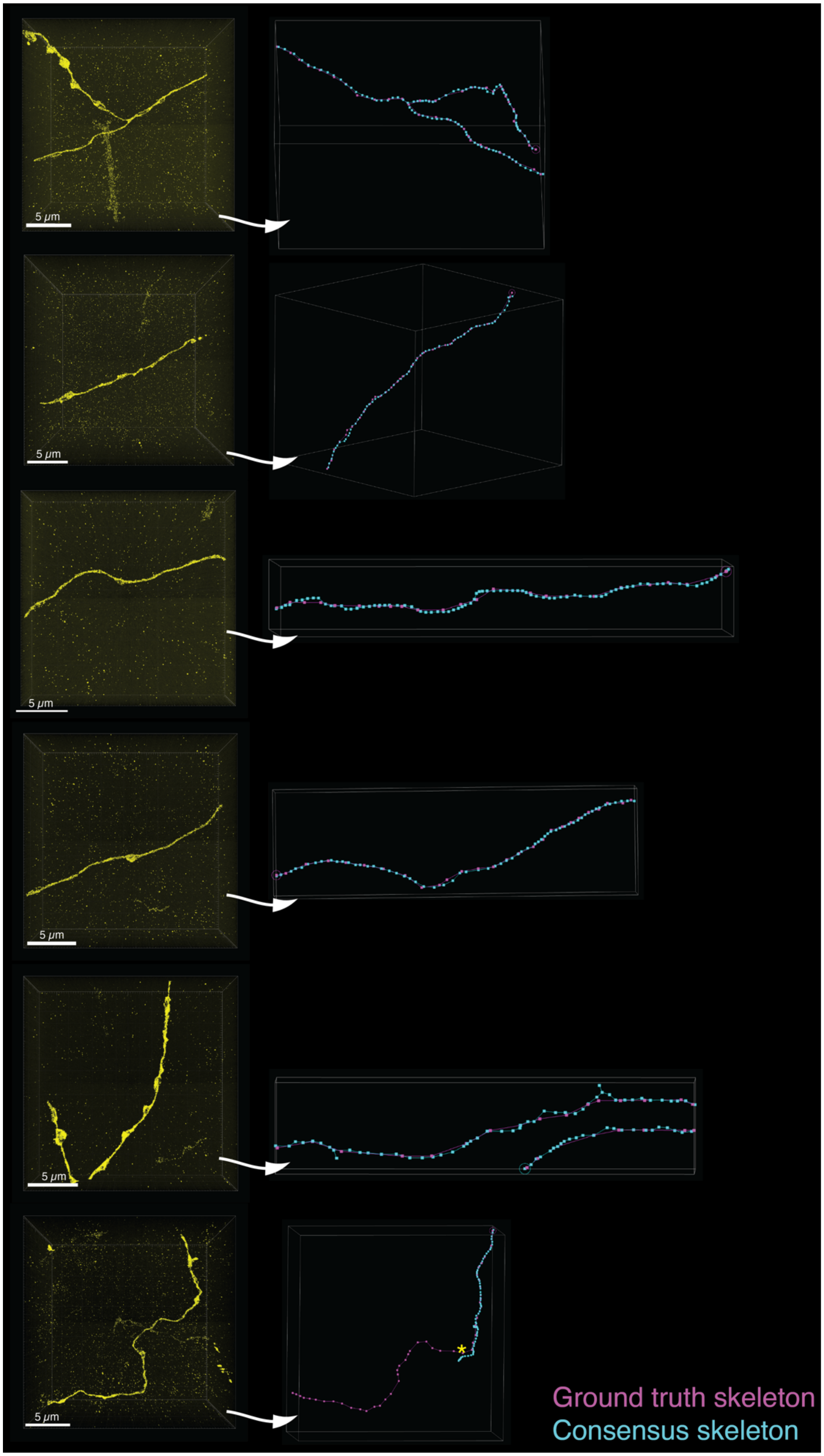
5 μm
5 μm
5 μm
5 μm
5 μm
5 μm
Ground truth skeleton
Consensus skeleton

**Fig. S14. Traceability of axons evaluated against ground truth from sparse eGFP expression (part 2).** *Left:* Volumetric renderings of eGFP signal (yellow) in LICONN volumes recorded in cortex of *Thy1-eGFP* animals with cytosolic expression of eGFP in a sparse subset of neurons. *Right*: Renderings of skeletons of the same structures generated from the LICONN structural channel (consensus skeletons, cyan) by two human annotators blinded to the eGFP channel. These are overlaid with skeletons that were generated by an independent annotator, taking both the LICONN structural channel and the eGFP signal into account (ground truth skeletons, magenta). In the lowest panel, a tracing error (asterisk) occurred in the consensus skeletons generated by the blinded human annotators. Note that volume renderings for the eGFP signal and skeletons are from different camera positions. eGFP data was visualized with Imaris software. In total, 37 axon stretches (880 µm cumulative length of eGFP-expressing axons, recorded across *n*=3 technical replicates across *n*=2 animals in cortex) were analysed.

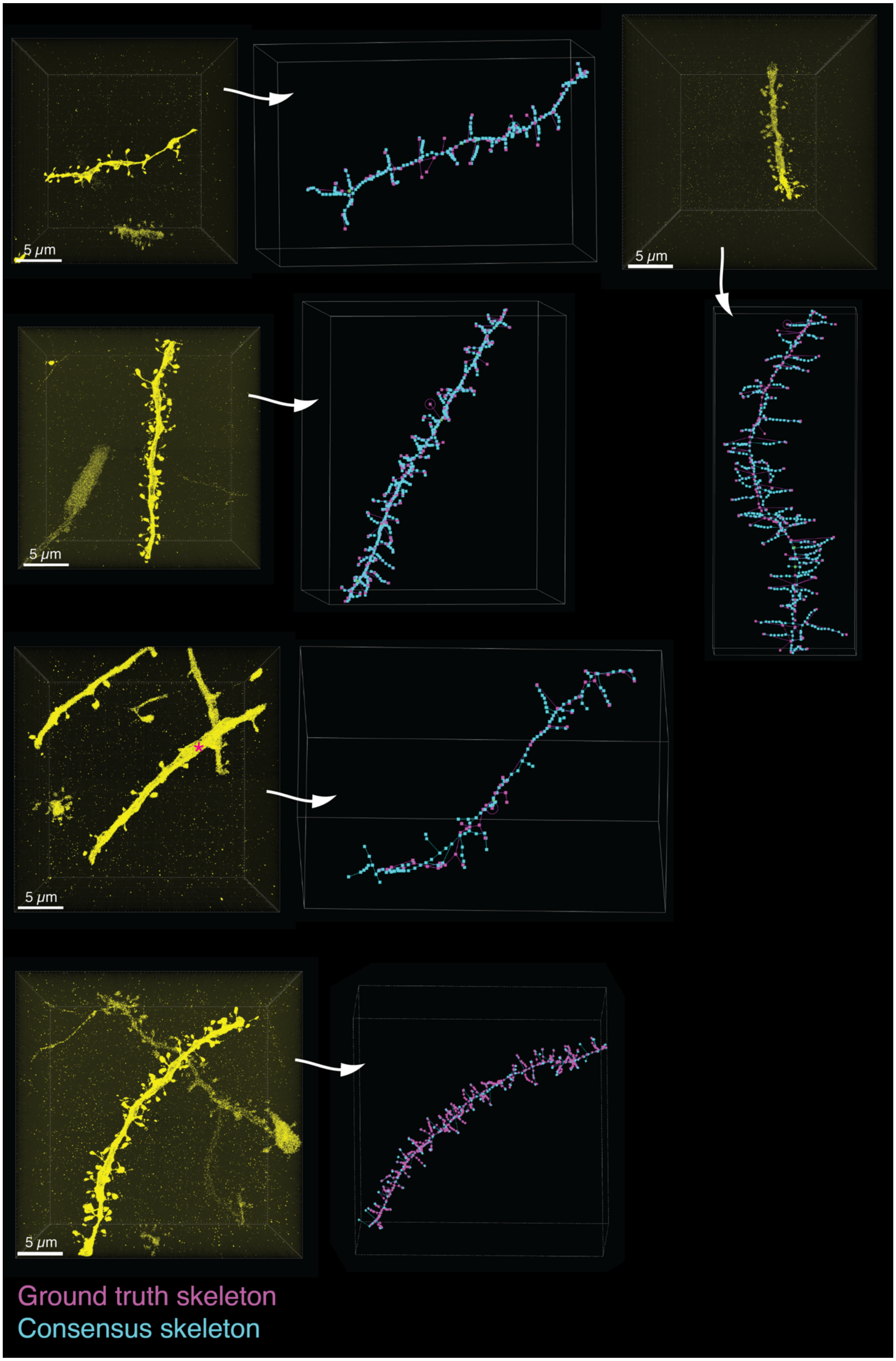
5 μm
5 μm
5 μm
5 μm
5 μm
Ground truth skeleton
Consensus skeleton

**Fig. S15. Traceability of dendritic evaluated against ground truth from sparse eGFP expression.** Volumetric renderings of eGFP signal (yellow) in LICONN volumes recorded in the hippocampal CA1 region (stratum oriens or stratum radiatum) of *Thy1-eGFP* animals with cytosolic expression of eGFP in a sparse subset of neurons. Renderings of skeletons of the same structures generated from the LICONN structural channel (consensus skeletons, cyan) by two human annotators blinded to the eGFP channel. These are overlaid with skeletons that were generated by an independent annotator, taking both the LICONN structural channel and the eGFP signal into account (ground truth skeletons, magenta). In the second to last volume, the dendrite stretch used for analysis is indicated by an asterisk. Note that camera positions are different in the volume renderings of the eGFP signal and skeletons. eGFP data was visualized with Imaris software. Analysis across 5 datasets from *n*=3 technical replicates across *n*=2 animals.

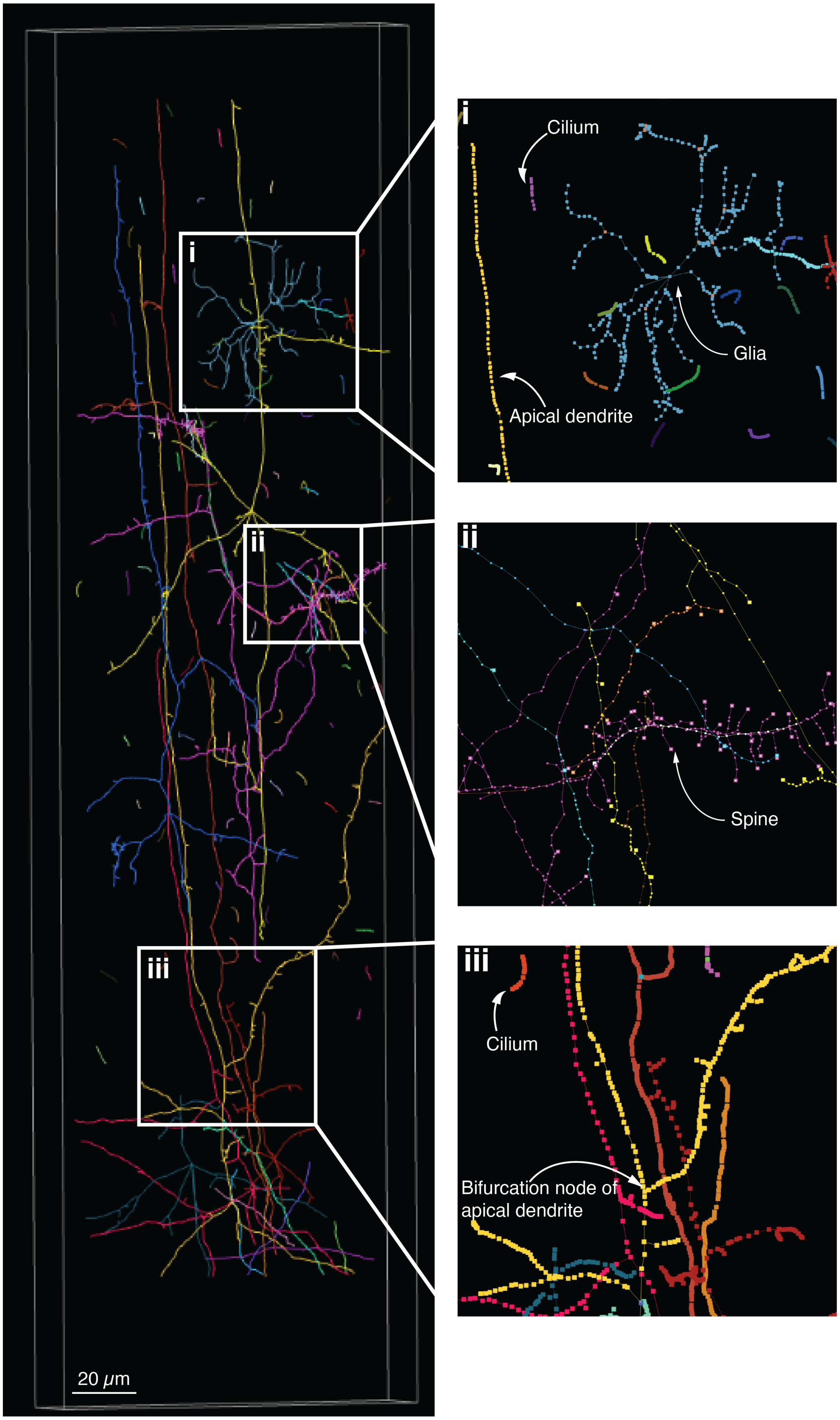

i
ii
iii
20 μm
Cilium
Glia
Apical dendrite
Spine
Bifurcation node of
apical dendrite

**Fig. S16. Tracing of neuronal structures in cortex.** Manually generated skeletons in the dataset in Fig. 1A, showing traceability of axons and dendrites across borders of fused imaging tiles. Magnified views indicate examples of glial cells, primary cilia used in further analysis (Fig. 5) and dendrites. Spines were only comprehensively traced in the right part of the dendrite indicated in the magnified panel ii.

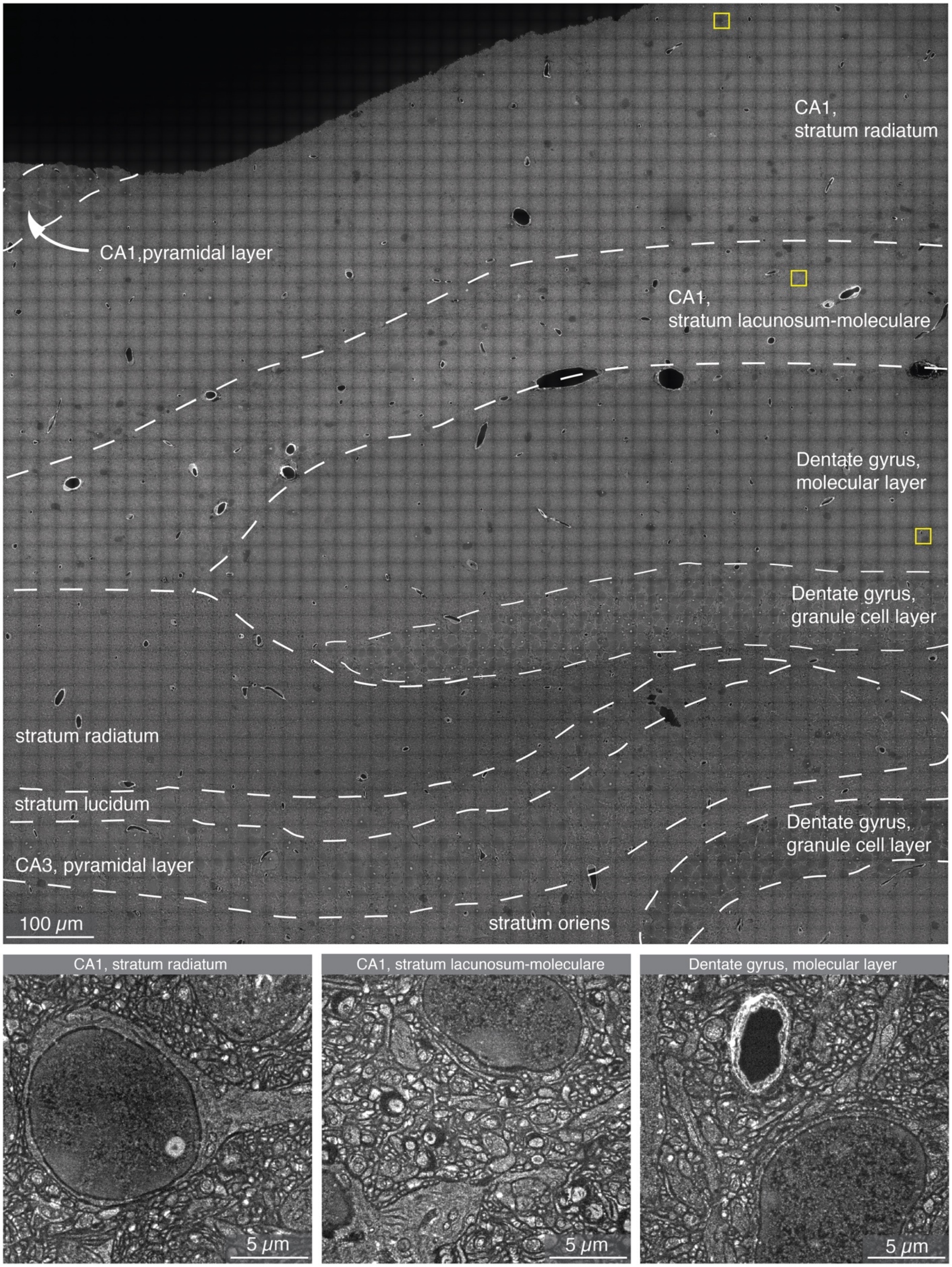
CA1,
stratum radiatum
CA1,pyramidal layer
CA1,
stratum lacunosum-moleculare
Dentate gyrus,
molecular layer
Dentate gyrus,
granule cell layer
stratum radiatum
stratum lucidum
Dentate gyrus,
granule cell layer
CA3, pyramidal layer
100 μm
stratum oriens
CA1, stratum radiatum
CA1, stratum lacunosum-moleculare
Dentate gyrus, molecular layer
5 μm
5 μm
5 μm

**Fig. S17. Hippocampal architecture analyzed with LICONN.** Hippocampal overview composed of 2,969 high-resolution tiles from spinning disc confocal microscopy with delineation of subregions and layers. Yellow boxes indicate the regions magnified in the bottom panels.

**A**

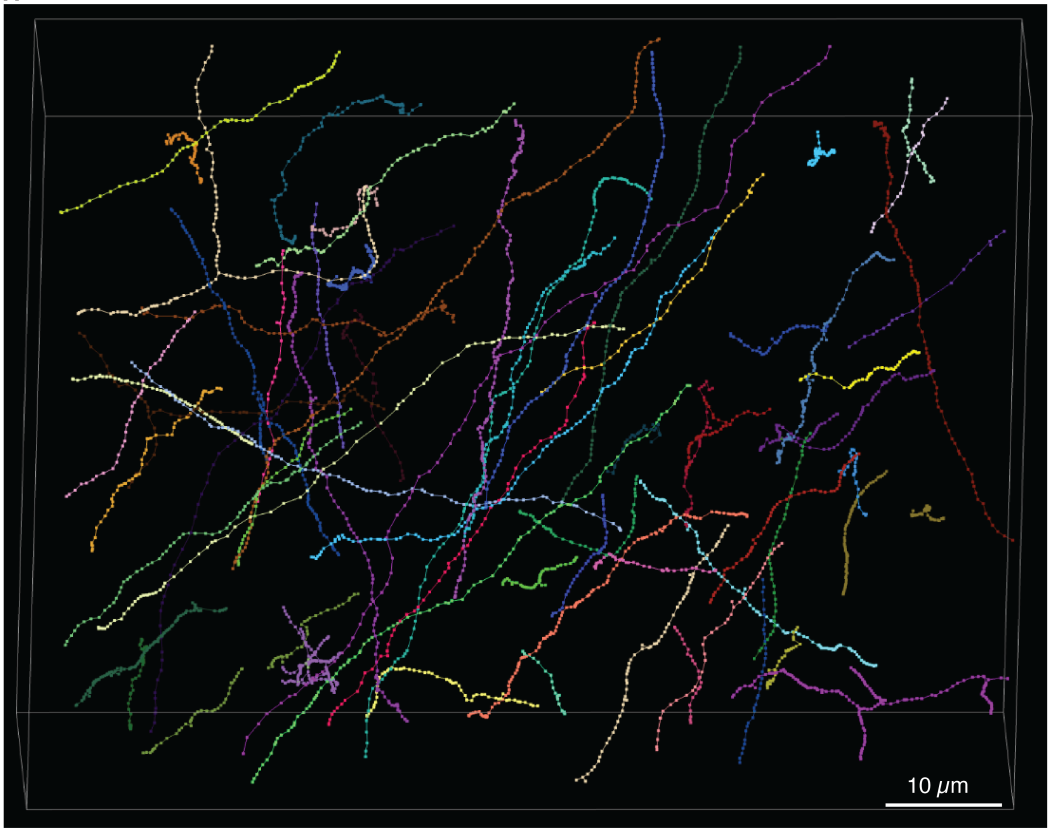


**B**

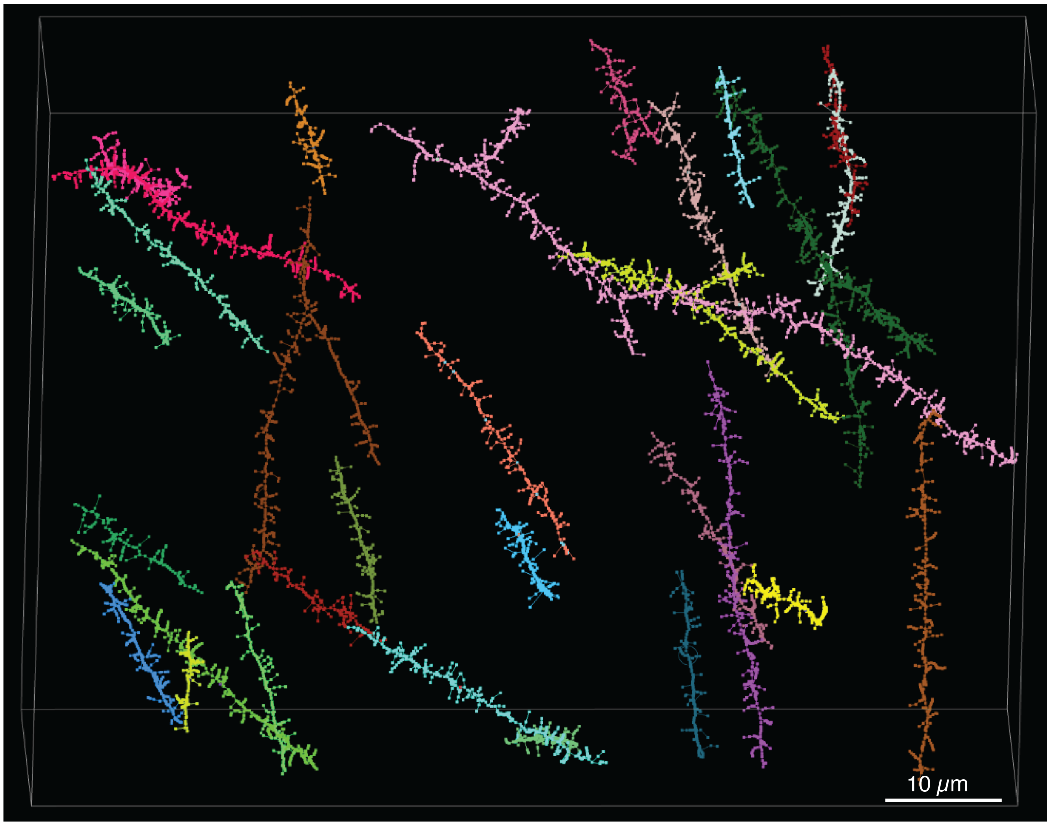

**Fig. S18. Manual tracing of axons and dendrites in hippocampus.** Manually generated skeletons in the dataset in Fig. 2A, used for evaluation of automated segmentation accuracy. Axons and dendrites were automatically selected at random from the output of a semantic classifier (see Methods). **(A)** 3D-rendering of the 69 traced axons. **(B)** 3D-rendering of the 30 traced dendrite stretches with 1,041 dendritic spines. Dendritic spines are indicated as branches to dendritic shafts.

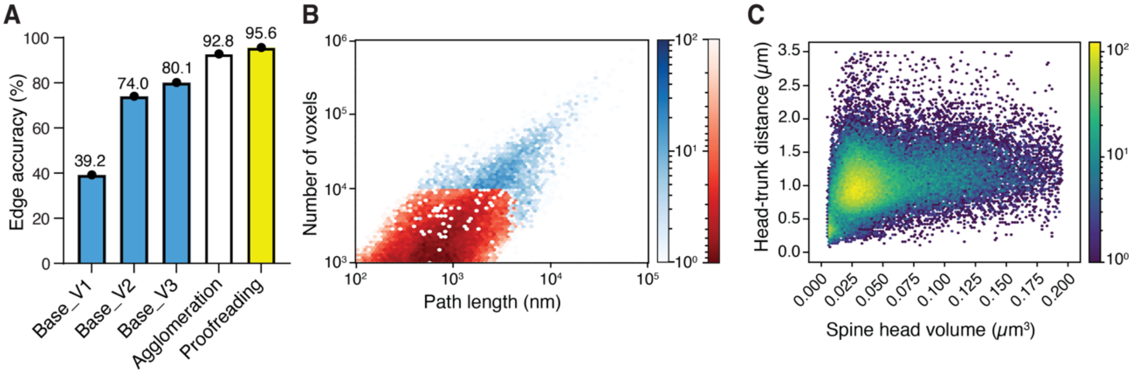


**Fig. S19. Evaluation of the performance of automated segmentation. (A)** Edge accuracy for the various stages of FFN segmentation. Blue bars: Base segmentations with models V1-V3 using increasing amounts of training data. White bar: Edge accuracy after automated agglomeration of the base agglomeration using the final model (V3). Yellow bar: Edge accuracy after additional manual proofreading. **(B)** Segment size (number of voxels) versus path length of individual segments for the segmentation in Fig. 2. Segments that entered the final agglomeration are indicated in blue. Segments below 10,000 voxels volume were disregarded in the agglomeration (red), except for a small number of longer segments with volume below 10,000 voxels. (**C**) Scatter plot of head-to-trunk distance vs. spine head volume for dendritic spines in the segmentation volume in Fig. 2A.

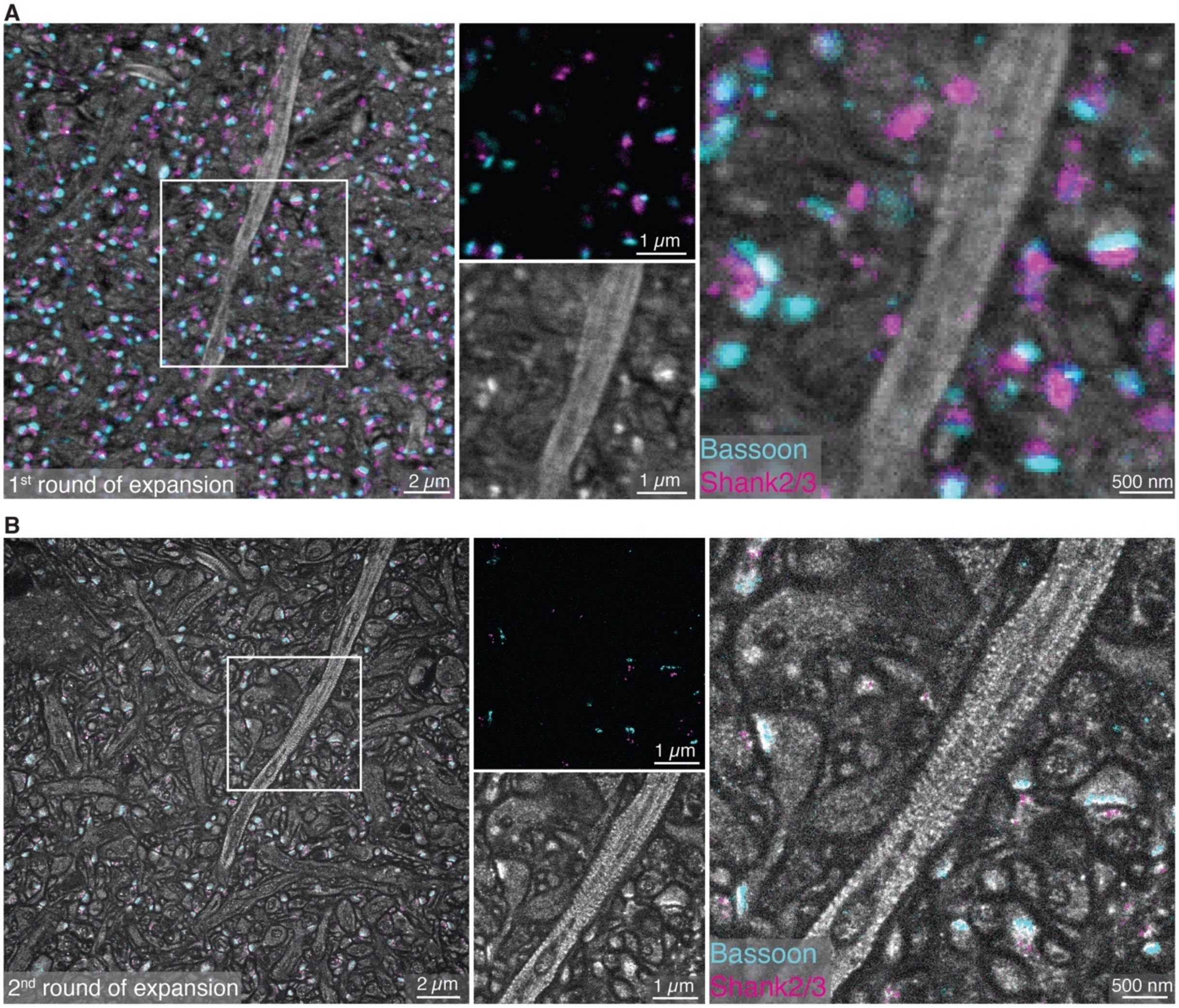


**Fig. S20. Improved resolution in structural and molecular LICONN channels. (A)** Confocal image in the hippocampal CA1 region (stratum oriens) after the first LICONN expansion step with immunolabeling for Bassoon (cyan) and Shank2/3 (magenta) overlaid with the structural channel (gray). *Left*: overview. *Middle, right*: Immunolabeling and structural channels shown separately for the region indicated by the box in the left panel and magnified view of the overlaid signal in the same region. **(B)** Same region imaged with the same objective lens and microscope after the second expansion step in LICONN. Resolution is increased both for the structural and immunolabeling channels, revealing neurites as separate structures and the periodic protein-density modulation induced by the actin/spectrin cytoskeletal lattice in the prominent neurite in the magnified view. At this resolution, molecular signals can be clearly assigned to individual pre- and post-synapses. Same dataset as in Fig. 1D.

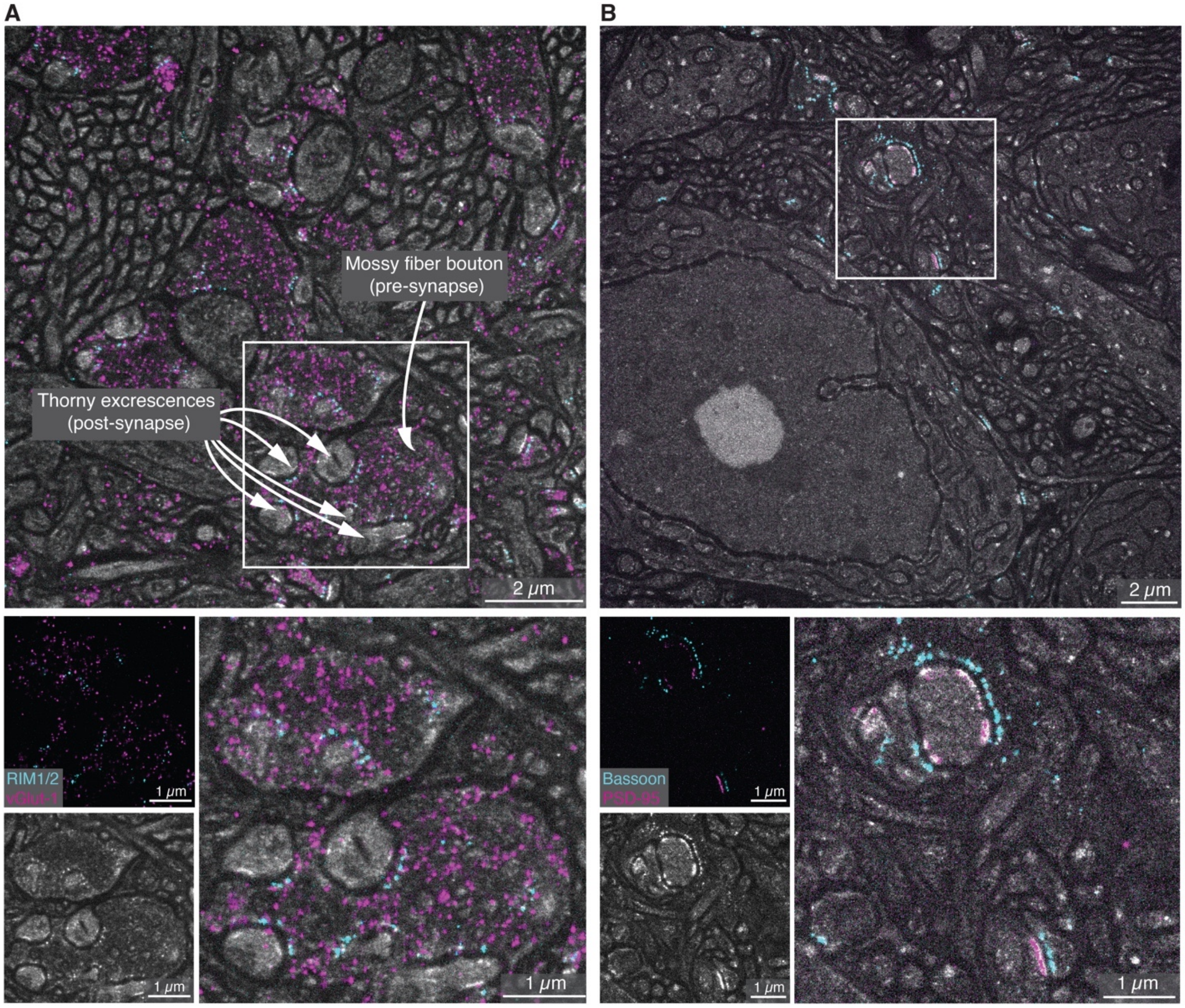


**Fig. S21. Synaptic labeling in mossy fiber boutons in the CA3 stratum lucidum. (A)** Mossy fiber boutons forming synapses with complex spines (thorny excrescences) at proximal dendrites of CA3 pyramidal neurons, interspersed with bundles of mossy fibers (non-myelinated axons of DG granule cells). Single plane of a LICONN imaging volume including the structural channel (gray) and immuno-labeling for vGlut1 (vesicular glutamate transporter 1, magenta), a synaptic vesicle marker in excitatory synapses, and the active zone marker RIM1/2 (cyan). *Bottom*: Immunolabeling and structural channels displayed separately and as overlay for the two mossy fiber boutons and their postsynaptic partners in the box in the top panel. **(B)** Similar measurement with immunolabeling for pre-synaptic Bassoon and the post-synaptic scaffolding protein PSD-95, present at excitatory synapses. Immunolabelling for RIM1/2 and vGlut1 is representative of $n$=2 technical replicates and Bassoon/PSD95 of $n$=3 replicates across $n$=2 animals.

**A**

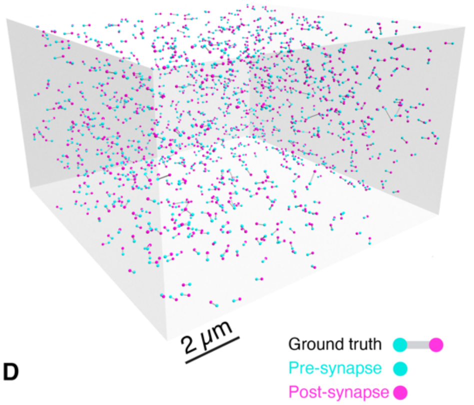


**B**

Immunolabeling based detection

Pre-synapse detection

| | Precision | Recall | F1 |
|---|---|---|---|
| Initial | 0.96 | 0.92 | 0.94 |
| Post-processed | 0.95 | 0.94 | 0.94 |

Post-synapse detection

| | Precision | Recall | F1 |
|---|---|---|---|
| Initial | 0.98 | 0.89 | 0.93 |
| Post-processed | 0.94 | 0.96 | 0.95 |

Full-synapse detection (paired)

| | Precision | Recall | F1 |
|---|---|---|---|
| Initial | 0.93 | 0.82 | 0.87 |
| Post-processed | 0.89 | 0.90 | 0.90 |

**C**

Deep-Learning based detection

Pre-synapse detection

| | Precision | Recall | F1 |
|---|---|---|---|
| Initial | 0.95 | 0.91 | 0.93 |
| Post-processed | 0.95 | 0.93 | 0.94 |

Post-synapse detection

| | Precision | Recall | F1 |
|---|---|---|---|
| Initial | 0.97 | 0.86 | 0.92 |
| Post-processed | 0.93 | 0.93 | 0.93 |

Full-synapse detection (paired)

| | Precision | Recall | F1 |
|---|---|---|---|
| Initial | 0.94 | 0.82 | 0.87 |
| Post-processed | 0.90 | 0.89 | 0.89 |

**D**

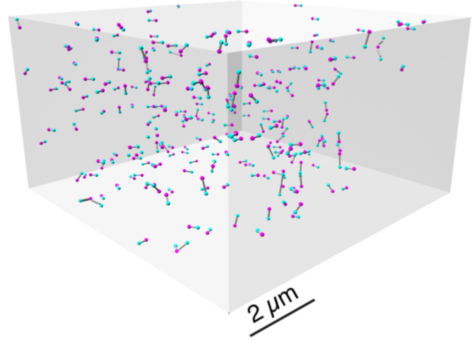


**E**

Immunolabeling based detection

Pre-synapse detection

| | Precision | Recall | F1 |
|---|---|---|---|
| Initial | 0.92 | 0.89 | 0.90 |
| Post-processed | 0.92 | 0.93 | 0.93 |

Post-synapse detection

| | Precision | Recall | F1 |
|---|---|---|---|
| Initial | 0.99 | 0.87 | 0.92 |
| Post-processed | 0.97 | 0.94 | 0.95 |

Full-synapse detection (paired)

| | Precision | Recall | F1 |
|---|---|---|---|
| Initial | 0.94 | 0.83 | 0.88 |
| Post-processed | 0.92 | 0.90 | 0.91 |

**F**

Deep-Learning based detection

Pre-synapse detection

| | Precision | Recall | F1 |
|---|---|---|---|
| Initial | 0.97 | 0.87 | 0.92 |
| Post-processed | 0.97 | 0.91 | 0.94 |

Post-synapse detection

| | Precision | Recall | F1 |
|---|---|---|---|
| Initial | 1.00 | 0.86 | 0.93 |
| Post-processed | 0.98 | 0.93 | 0.95 |

Full-synapse detection (paired)

| | Precision | Recall | F1 |
|---|---|---|---|
| Initial | 0.97 | 0.83 | 0.90 |
| Post-processed | 0.95 | 0.90 | 0.92 |

**Fig. S22. Immunolabeling and deep-learning based detection of excitatory synapses. (A)** Rendering of ground truth (proofread) immunolabeling-based excitatory synapse detections in the dataset in Fig. 3G. (**B**) Precision, recall and F1 score of computational immunolabeling-based detection of pre-synapses, post-synapses, and fully assembled synapses, evaluated using the manually generated ground truth displayed in panel (A). Values are given both for the base detection (initial) and with post-processing (see Methods). (**C**) Precision, recall and F1 score for deep-learning-based prediction of pre-synapses, post-synapses, and fully assembled synapses from the structural LICONN channel in the same dataset. The dataset was not included in training. (**D**) Rendering of ground truth (proofread) immunolabeling based excitatory synapse detections in the dataset in Fig. 4L. (**E**) Precision, recall and F1 score of computational immunolabeling-based detection of pre-synapses, post-synapses, and fully assembled synapses, evaluated using the manually generated ground truth displayed in (D). (**F**) Precision, recall and F1 score for deep-learning-based prediction of pre-synapses, post-synapses, and fully assembled synapses from the structural LICONN channel in the same dataset. The dataset was not included in training.

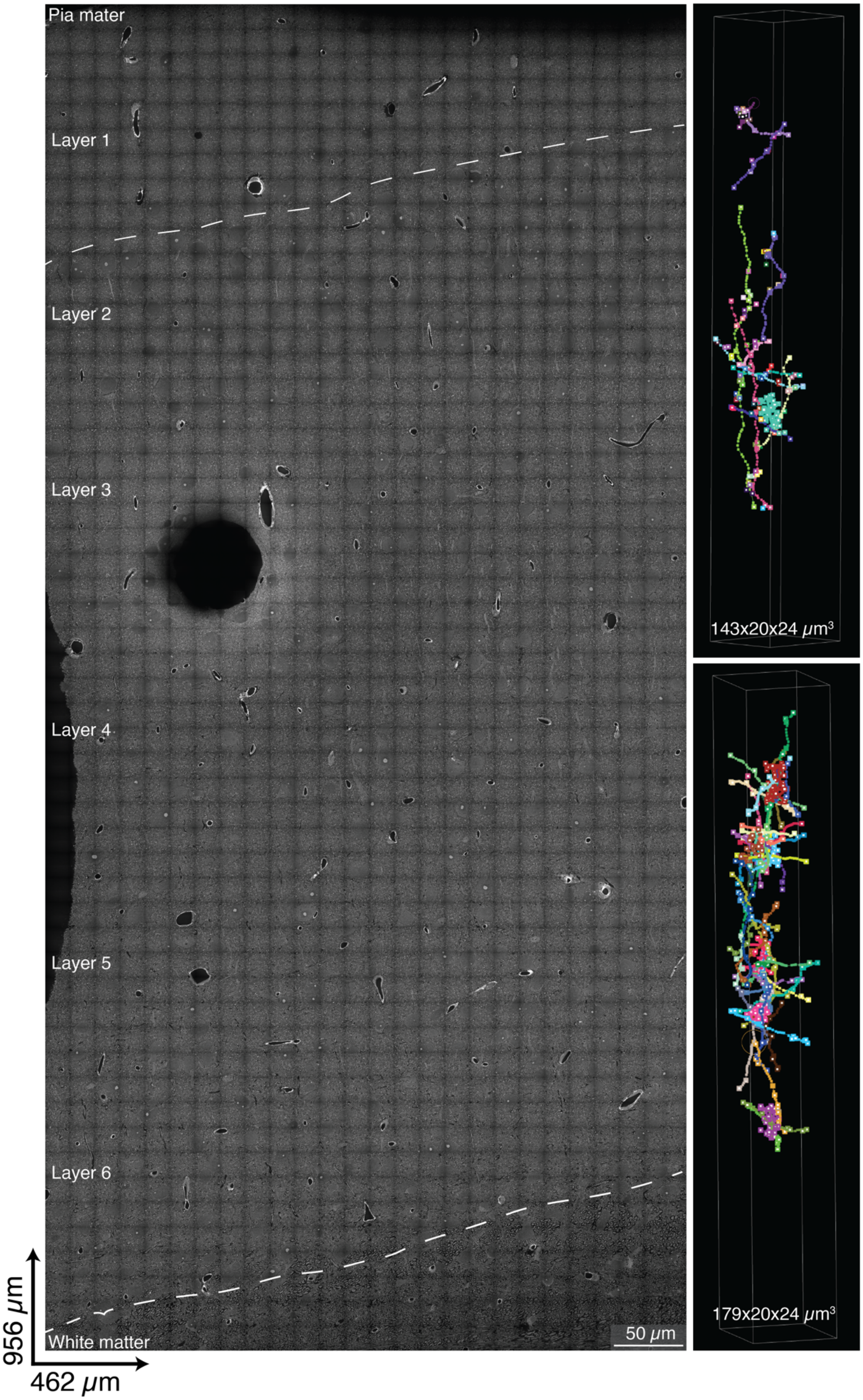

Pia mater
Layer 1
Layer 2
Layer 3
Layer 4
Layer 5
Layer 6
White matter
50 μm
956 μm
462 μm
143x20x24 μm³
179x20x24 μm³

**Fig. S23 Molecularly informed connectivity analysis (manual tracing).** *Left*: Overview of LICONN sample in cortex, spanning from pia mater to the white matter. *Right*: Manual tracings from the same specimen, used for analysis in Fig. 4A-K. The dark circular region in the centre left represents a defect in the tissue-hydrogel hybrid (e.g. air bubble) or in sample mounting, which we occasionally observed. Such defects are not specific to LICONN and did not impede our measurements, such that we did not screen samples for defects before imaging. If required for a particular application, such as scaling to much larger volumes, detecting defects would be possible before LICONN imaging by coarse overview scanning of the samples.

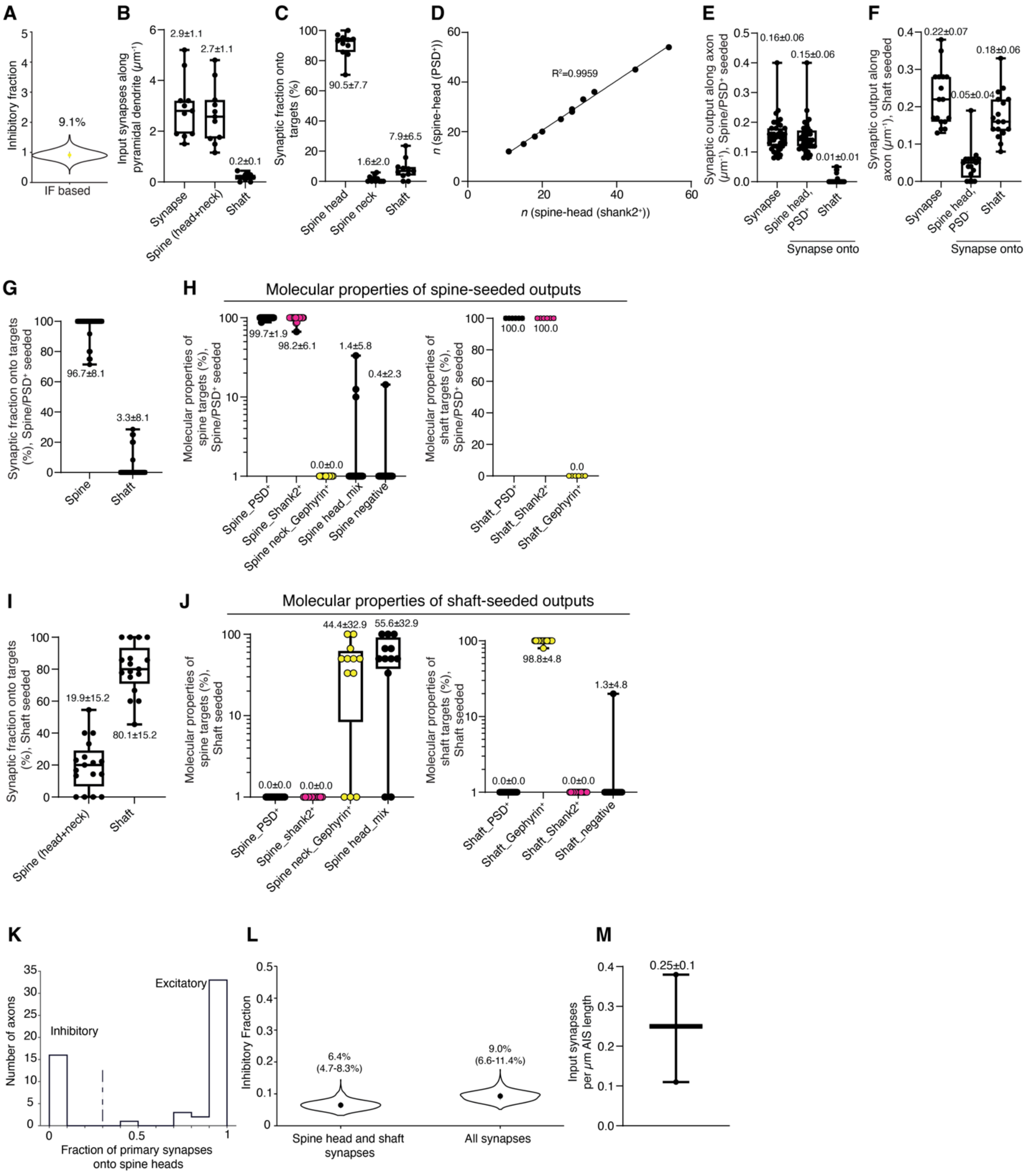


**Fig. S24. Morphology-based connectivity analysis (manual tracing) in mouse somatosensory cortex.** (**A**) Bootstrap analysis of the immunolabeling-based excitation/inhibition ratio measurement in Fig.2F (351 total immunolabeling positive synapses). Distribution across $n = 1000$ bootstrap samples, median 9.1%, 10$^{th}$-90$^{th}$ percentile 7.2-11.3%). (**B-C**) Input properties of spiny dendrites, based on structural identification of synapses (disregarding the molecular information). (B) Density of synaptic inputs onto a spiny dendrite per unit length, including the total number of synapses (335), those onto spines (332), and onto shafts (23). (**C**) Fraction of synaptic inputs according to target location (spine head, spine neck, shaft). (**D**) Number of dendritic spines that feature a PSD vs. number of spines that are positive for Shank2. Linear regression yields $R^2$ of 0.996. (**E-J**) Output target locations of spine- and shaft- seeded axons. Only axons were taken that were seeded by a spine with a single synapse (PSD positive, regardless of presence of immunolabeling signal). (E) Density of synaptic outputs per unit length of spine-seeded axons, according to target location (total, spine head, shaft). (F) Density of synaptic outputs per unit length of shaft-seeded axons, according to target location (total, spine head, shaft). (G) Relative proportion of synaptic output target location for spine-seeded axons. (H) Molecular properties of spine-seeded axons outputs. *Left*: Outputs onto spines. 98% of spine head synapses were Shank2 positive. 1.4% of Shank2 positive spine heads had an additional synapse that was Gephyrin positive. In 0.4% no immunolabeling was present but a synapse could be detected due to the presence of a PSD in the structural channel. *Right*: Output onto shafts. 100% of the shaft outputs of these spine-seeded axons were positive for Shank2. (I) Relative proportion of synaptic output target location for shaft-seeded axons. (J) Molecular properties of shaft-seeded axons outputs. *Left*: Outputs onto spines. 44% of these synapses were onto spine necks, which were Gephyrin positive. 55% of the outputs of these shaft-seeded axons targeted spine heads with a Gephyrin-positive connection while the same spine head was also contacted by an excitatory axon (Shank2 positive connection). *Right*: Output onto shafts. These were mostly Gephyrin-positive (99%). In 1% of cases, we identified a synaptic connection based on the presence of a pre-synaptic bouton with dense projection arrangement (Fig. 1B), which was negative for immunolabeling. (**K**) Distribution of axons (total 55 axons with ≥3 outputs), including both spine-head seeded (38) and shaft-seeded (17), according to the target location of their synaptic outputs (spine head or shaft). Synapses onto spine necks were not considered. The vast majority of axons either had a clear preference for spine heads or shafts, differentiating them into excitatory and inhibitory axons, respectively. The vertical line indicates the threshold for classifying them. Determining the relative number of synapses of these excitatory and inhibitory axons onto dendrites, yielded 6.5% inhibitory synapses. Additionally taking synapses onto spine necks and onto spine heads that received both an excitatory and an inhibitory synapse into account, overall inhibitory fraction increased to 9.4%. (**L**) Bootstrap analysis of same data as in (K). Distribution across $n = 1000$ bootstrap samples. (**M**) Density of input synapses along axonal initial segments for the 3 cells analyzed in Fig. 4J.

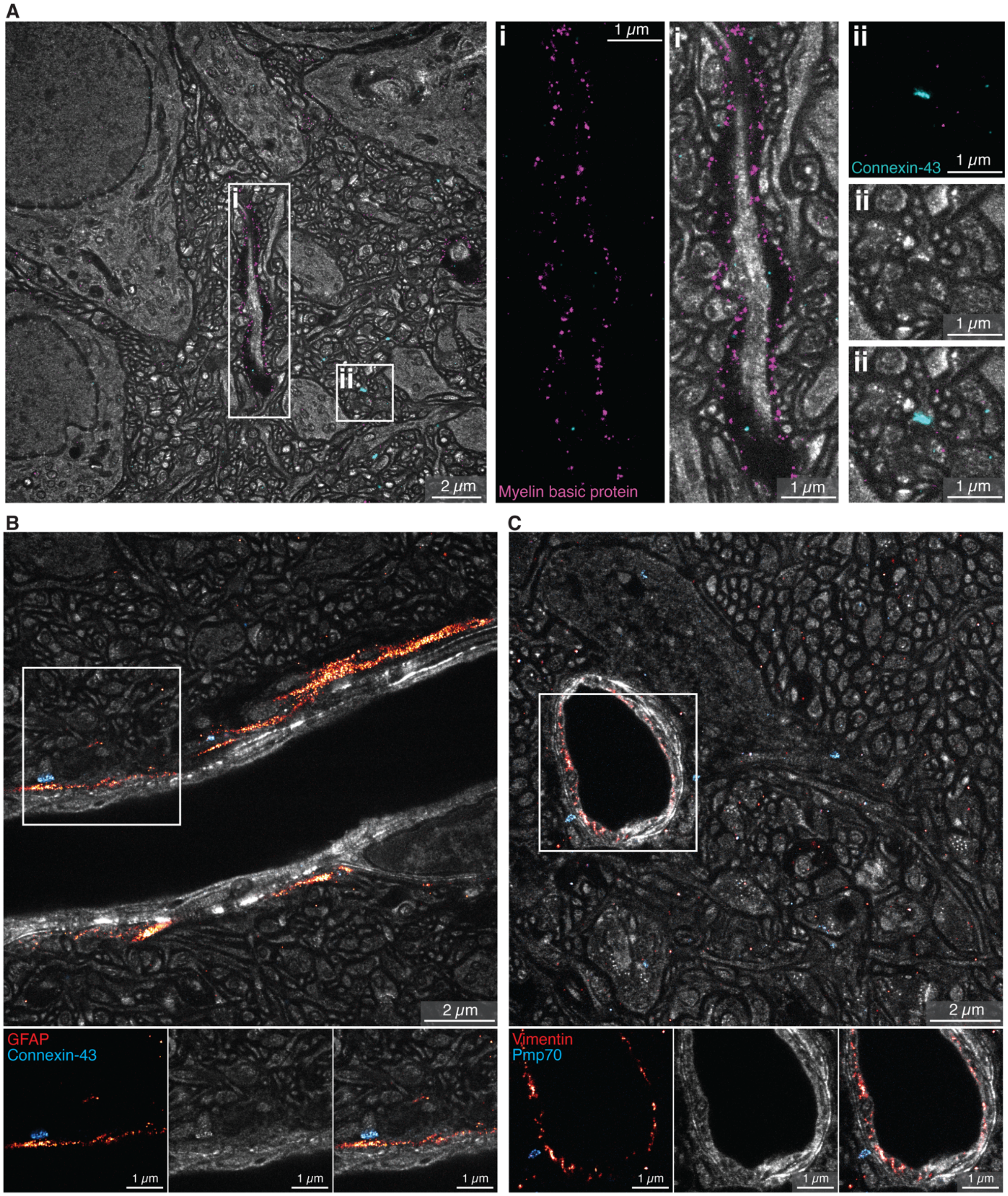


**Fig. S25. Molecular identification of cellular and subcellular structures. (A)** Single plane of a LICONN imaging volume in the hippocampal CA1 stratum radiatum, including the structural channel (gray) and immunolabeling for myelin basic protein (magenta) and the astrocytic gap junction protein Connexin-43 (cyan). Magnified views highlight structures expressing the respective molecular markers: (i) Immunolabeling for myelin basic protein identifies myelin sheaths, where the regions largely devoid of signal in the structural LICONN channel correspond to lipid-rich myelination. (ii) Connexin-43 indicates gap junctions. **(B)** Single plane from LICONN imaging volume in hippocampus, showing the structural channel (gray) and immunolabeling for glial fibrillary acidic protein (GFAP, orange) and Connexin-43 (cyan). GFAP is an intermediate filament protein, commonly used as astrocyte marker. Note the GFAP signal in the immediate vicinity of the capillary traversing the image and the cell nucleus on the right in the periphery of the blood vessel. Panels in the bottom show the immunolabeling and structural channels separately, as indicated by the box. **(C)** Single plane from a LICONN imaging volume in the CA3 stratum lucidum, with additional immunolabeling for vimentin (orange, here highlighting the endothelial lining of a blood capillary) and the peroxisomal marker Pmp70 (70 kDa peroxisomal membrane protein). Note that peroxisomes do not produce pronounced contrast in the structural (protein density) channel but can be identified by specific labeling. The displayed combinations of immunolabellings for MBP/Cnx-43 were replicated $n$=1 times, RIM1/2 and vGlut1 $n$=2 times, GFAP/Cnx-43 $n$=2 times, Vimentin/Pmp70 $n$=1 times.

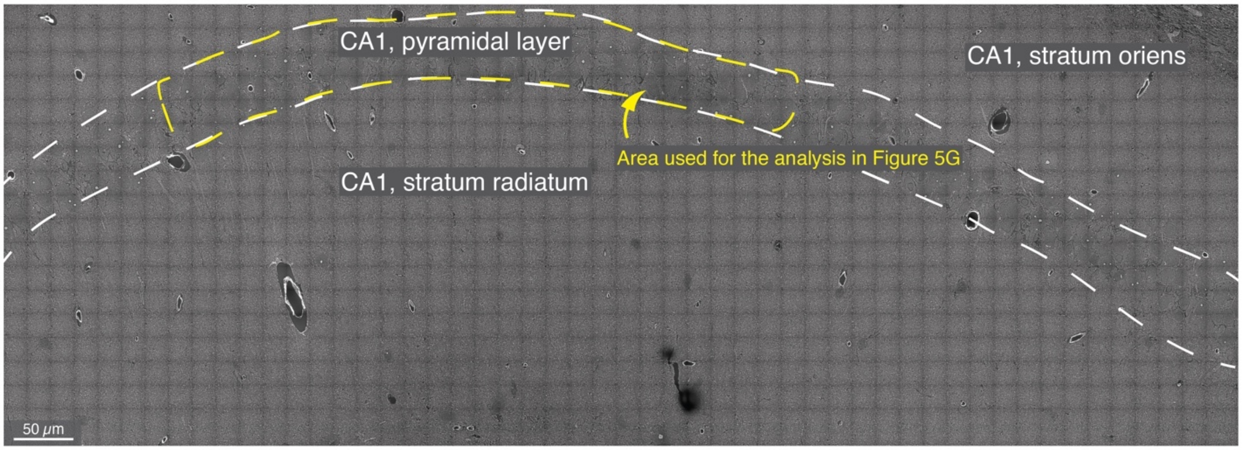


**Fig. S26. LICONN analysis in *Hnrnpu*$^{+/-}$ haploinsufficent mice.** Overview in the hippocampal CA1 region, tiled from high-resolution spinning disc confocal images. The region used for analysis of cilia length in Fig. 5 is indicated by the yellow dashed line.

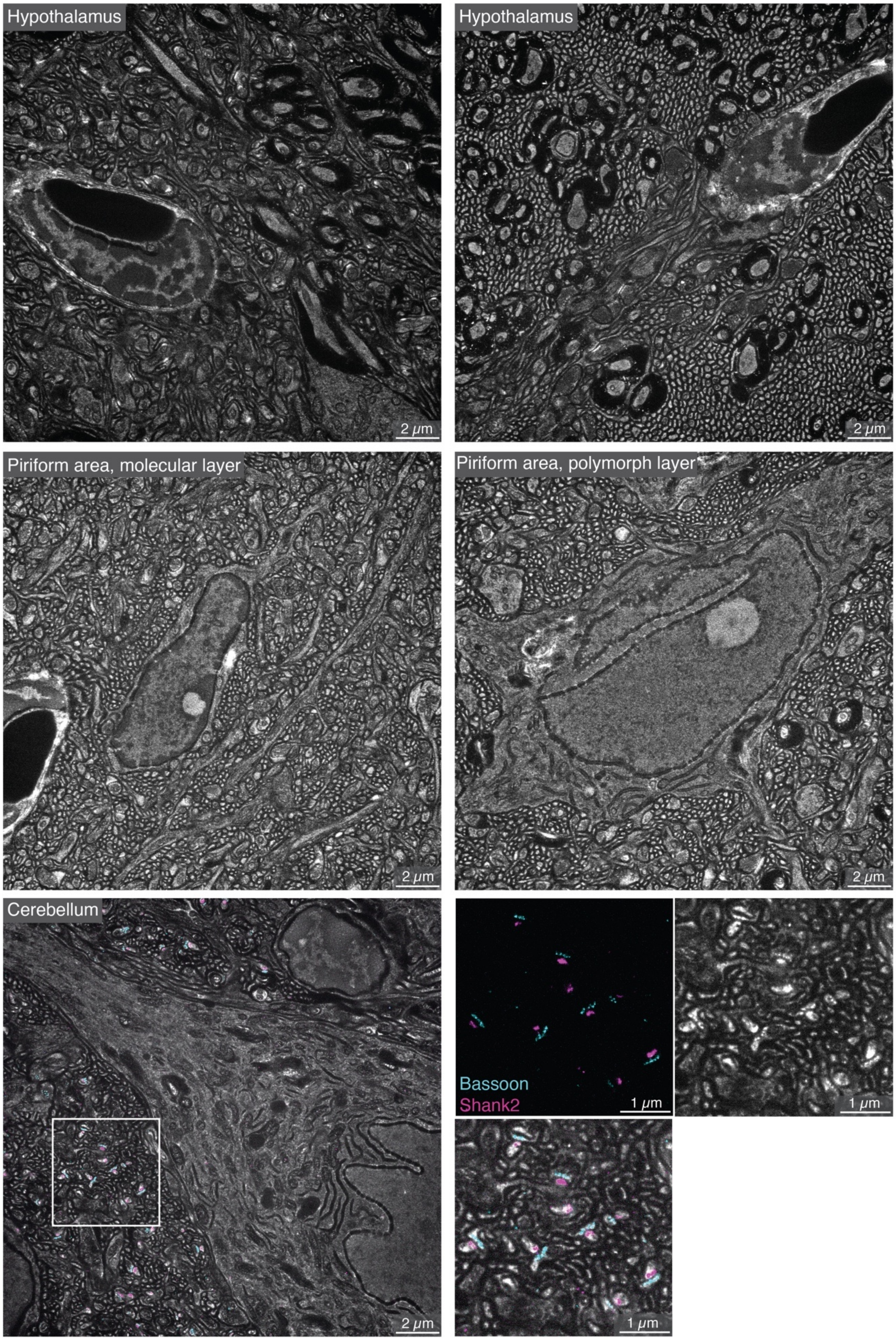
Hypothalamus
2 μm
Hypothalamus
2 μm
Piriform area, molecular layer
2 μm
Piriform area, polymorph layer
2 μm
Cerebellum
2 μm
Bassoon
Shank2
1 μm
1 μm
1 μm

**Fig. S27. LICONN analysis in diverse brain regions.** Single example planes from LICONN imaging volumes in hypothalamus, piriform area, and cerebellum. In the cerebellum, additional immuno-labeling for Bassoon and Shank2 was applied.

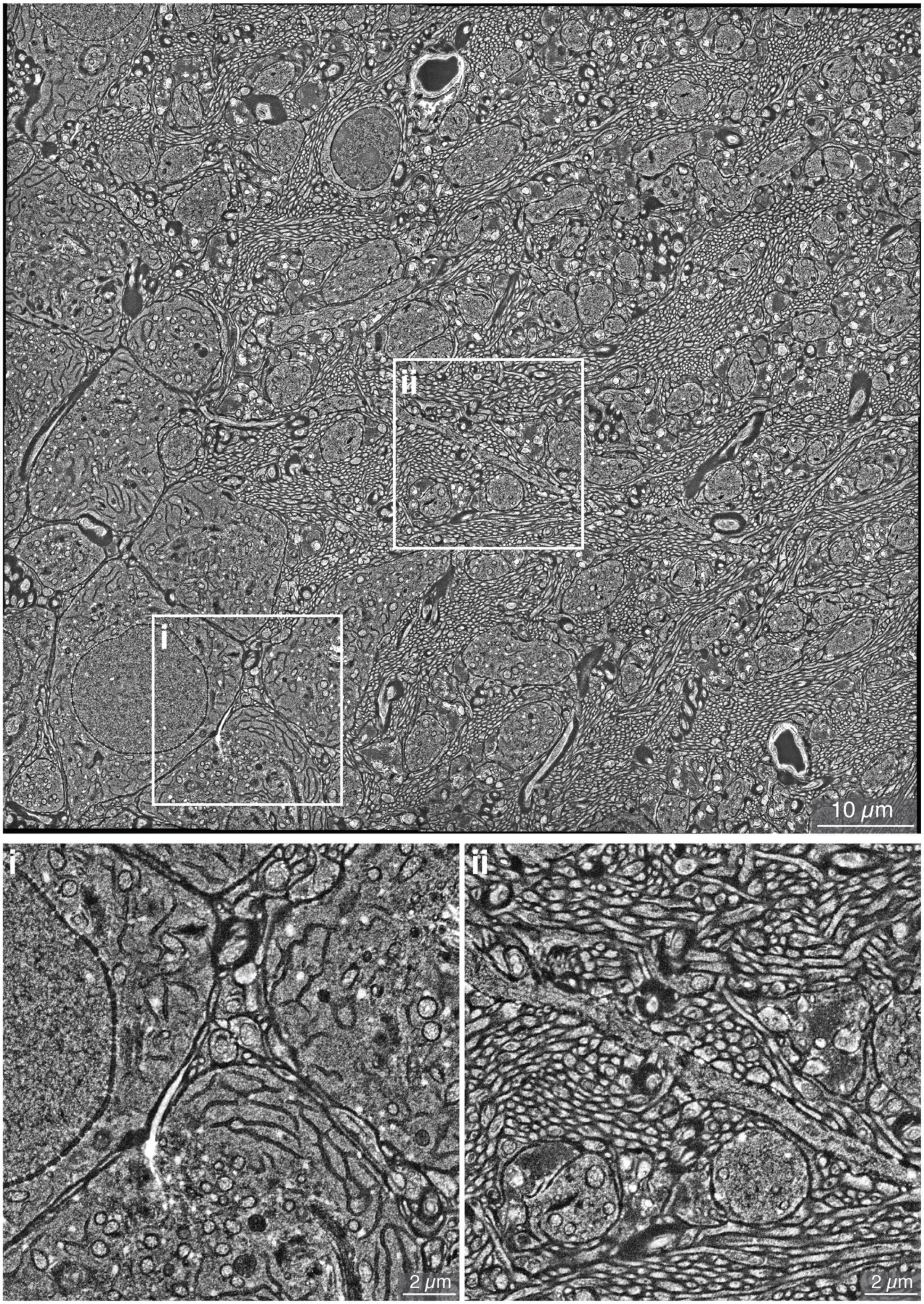
ii
i
10 μm
i
ii
2 μm
2 μm

**Fig. S28. LICONN imaging in the hippocampal CA3 stratum lucidum.** Single plane overview image with magnified views, illustrating the complex arrangement of cellular structures including a primary cilium and intracellular organelle diversity in (i) and mossy fiber bundles and an example of multiple shaft synapses in (ii). Overview imaging in CA3 stratum lucidum was performed in *n*=2 replicates, single tile measurements were performed in multiple technical replicates.

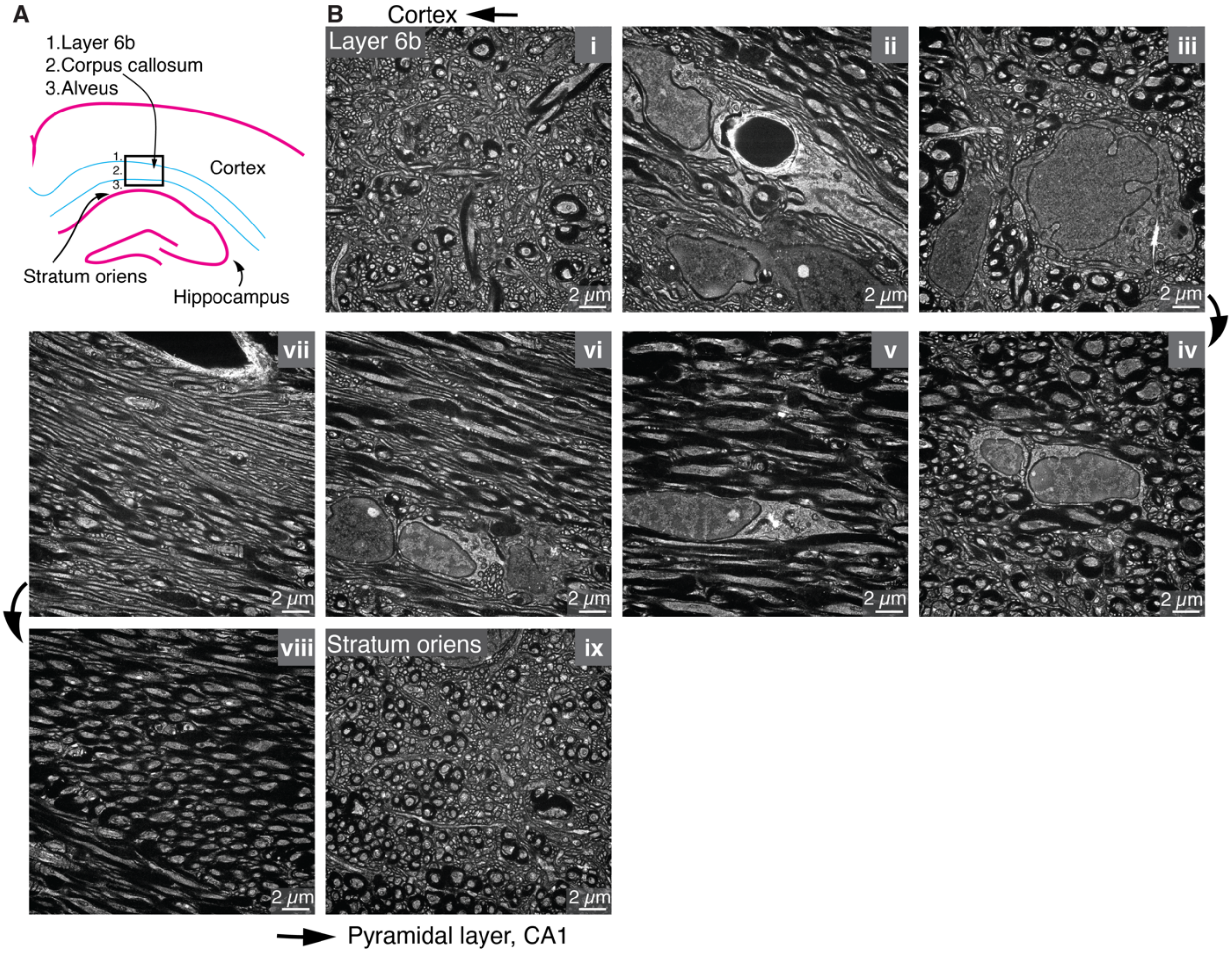


**Fig. S29. LICONN imaging in white matter. (A)** Schematic of a mouse coronal section with the region of imaging (box). (**B**) Single planes from individual LICONN imaging volumes spanning deep cortex, corpus callosum, alveus and hippocampus according to the schematic, revealing the distinct organization in the various regions. Imaging of the various layers was performed in one replicate.

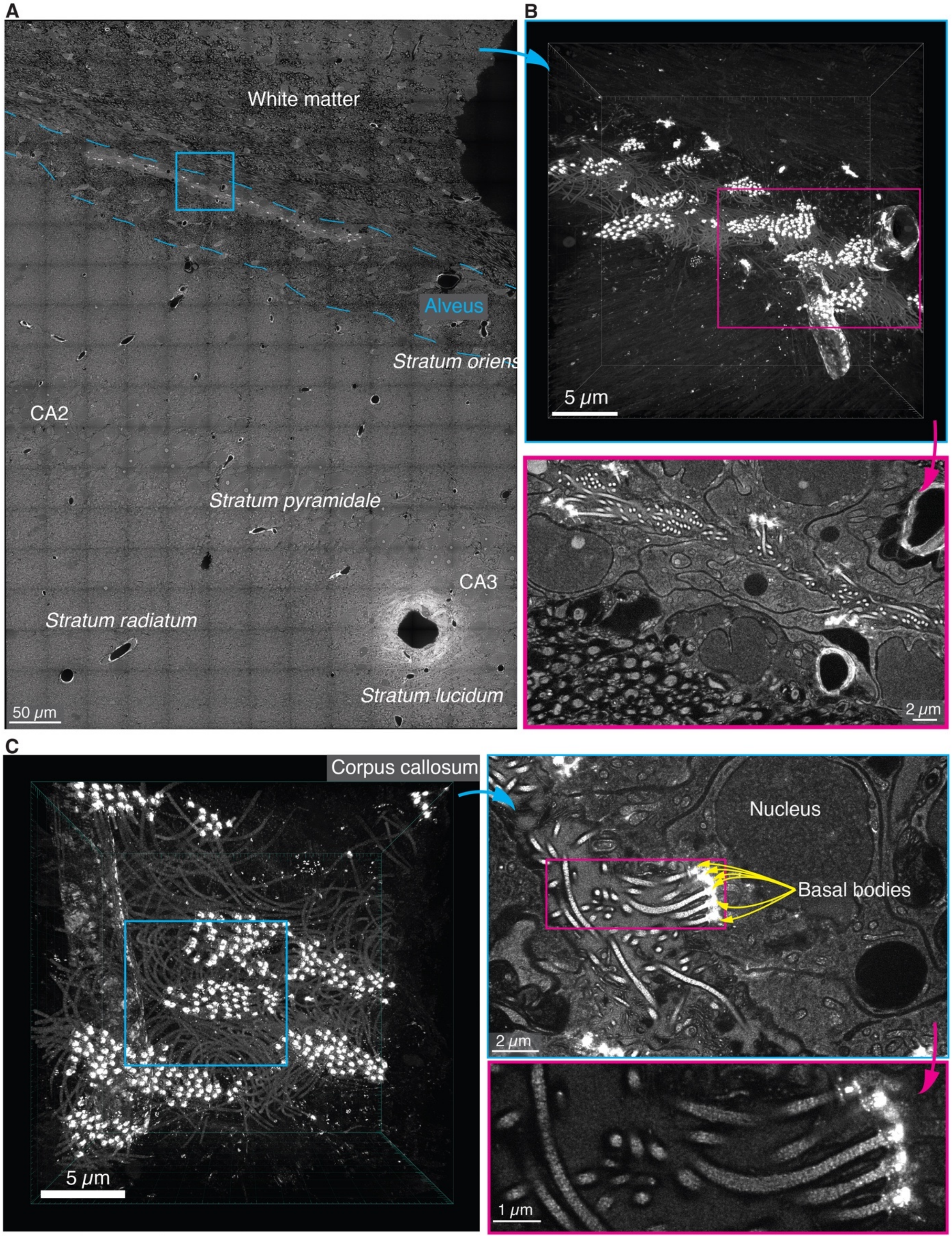


**Figure S30. Muli-ciliated cells at the transition of corpus callosum and alveus. (A)** *Left*: Regional overview image and 3D-rendering of a LICONN imaging volume (*right, top*, recorded with a 20x objective lens) of the region indicated by the white box in the overview, highlighting multi-ciliated cells. The bright structures correspond to basal bodies. *Right, bottom*: Single imaging planes from the same volume as indicated by the magenta box in the upper panel. Intensity lookup tables in the individual panels are adjusted to either emphasize the intensely labeled basal bodies or the local cellular architecture. **(B)** *Left*: Rendering of a LICONN imaging volume recorded with a high-NA objective lens in the same region from a different specimen. *Right*: Single imaging plane showing multi-ciliated cells and magnified view of cilia with intensity lookup table adjusted to visualize the axonemes of cilia. Representative of replicates in $n$=2 animals.

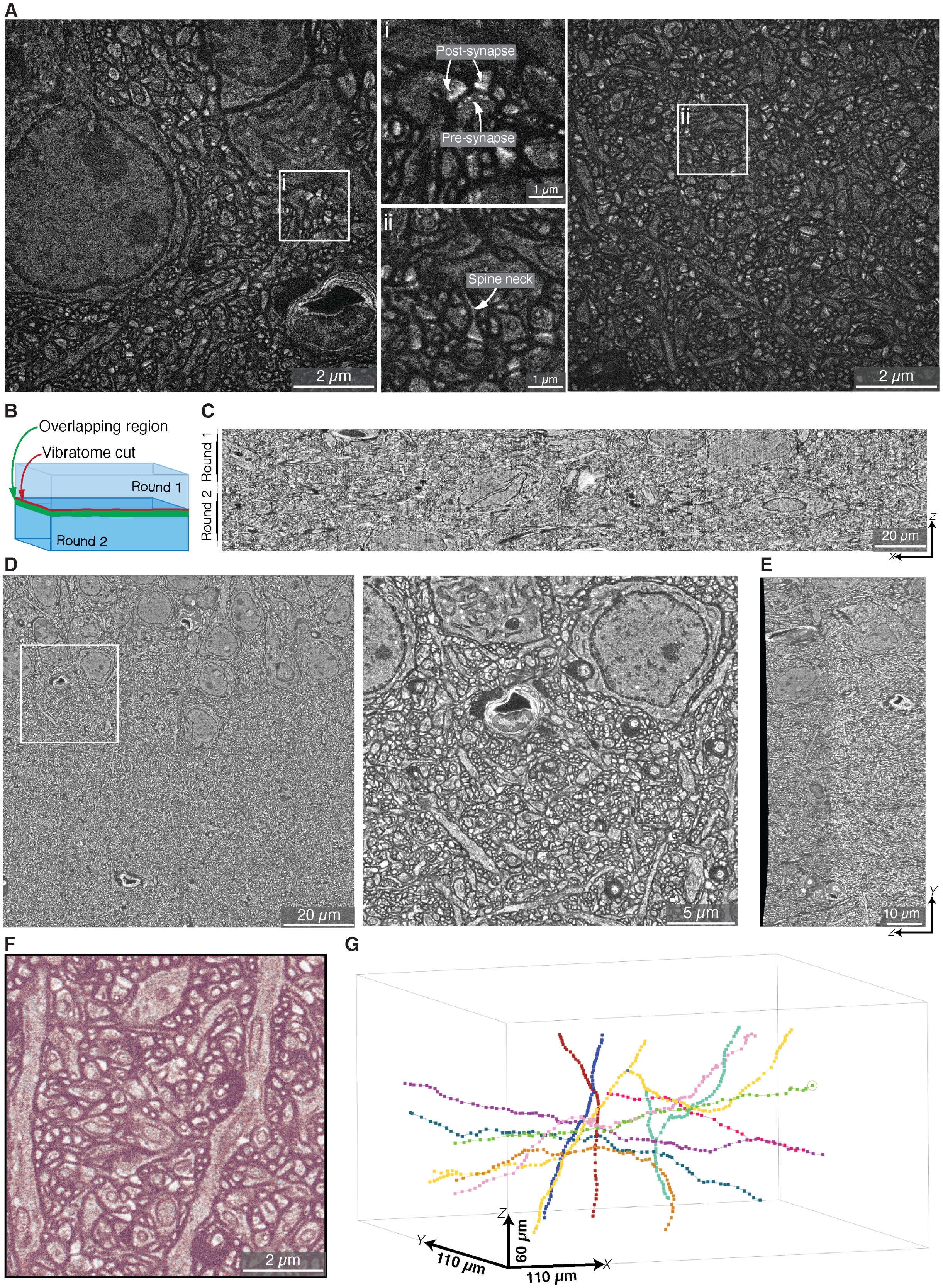


**Fig. S31: Scaling up in axial direction through iterative imaging and sectioning of the LICONN tissue-hydrogel hybrid.** (**A**) LICONN applied to 300 μm thick brain tissue slices from hippocampus, CA1 stratum radiatum. Single imaging planes in sub-regions of a multi-tile volume with magnified views of the boxed regions. Data quality in such thicker biological samples is equivalent to LICONN in thinner tissue sections. (**B**) Schematic of iterative block-face imaging and sectioning of the LICONN tissue/hydrogel hybrid. In imaging round 1, a multi-tile volume is acquired, then most of the imaged hydrogel layer is removed with a vibratome, and in round 2 a second multi-tile imaging volume deeper in the tissue is acquired. A region of continuous overlap is recorded in both imaging rounds, enabling lossless stitching and fusion of imaging volumes across rounds. (**C**) Axial (*xz*-)view of the full LICONN volume of panel A, fused from 72 subvolumes arranged on a 6x6x2 grid in 3D. Data were acquired in two imaging rounds at increasing depth (data displayed after coarse stitching step). The first imaging round had an axial extent of 466 µm (30.2 µm native tissue scale) and the subsequent vibratome cut was performed 320 µm below the hydrogel surface. The second imaging round had an axial extent of 353 µm (23 µm native tissue scale). Total multi-round acquisition time for the full volume was 3 h 36 min, including image acquisition for round 1 (2 h 04 min), sample handling and vibratome cutting between imaging rounds (15 min), and image acquisition for round 2 (1 h 17 min). (**D**) Single plane overview of the same volume in lateral (*xy*-)direction (109 x 109 $\mu m^2$ native tissue scale) after complete volume fusion. The magnified view of the boxed region focuses on a region with tile borders, showing seamless fusion (region adjacent to the view in the left image in panel A). (**E**) Single-plane axial (*xz*-)view after volume fusion in the same dataset, with seamless fusion across the two individual imaging rounds with vibratome cutting in between. (**F**) Single-plane lateral (*xy*-)view of a subarea in the axial overlap region between imaging rounds, displayed after volume fusion. Data from rounds 1 and 2 are color coded in red and green. Homogeneous coloring indicates voxel-exact matching of volumes from individual imaging rounds after the fusion process. (**G**) 3D-illustration of manually traced axons in the final stitched and fused volume, demonstrating traceability across borders between imaging rounds.

**Movie S1.**

**Manual segmentation of LICONN volume**. Fly-through of the original data in Fig. 1G, with step-by-step appearance of manually annotated segments.

**Movie S2.**

**Automated segmentation.** Rendering of original data in Fig. 2A with a subset of neurite segments from the FFN segmentation with comprehensive proofreading of neuronal structures.

**Movie S3.**

**Excitatory synaptic inputs and outputs of a cortical neuron based on deep-learning prediction of synapses.** Rotation of rendering in Fig. 4O.
